# Bimolecule detection for Extracellular Vesicle Screening

**DOI:** 10.1101/2020.07.23.217018

**Authors:** Hisako Kaneda, Yui Ida, Ryusuke Kuwahara, Izumi Sato, Takanari Nakano, Haruhiko Tokuda, Tsuyoshi Sato, Takayuki Murakoshi, Koichi Honke, Norihiro Kotani

**Author notes:** To whom correspondence should be addressed: Norihiro Kotani: Department of Biochemistry, Saitama Medical University, 38 Morohongo, Moroyama-machi, Iruma-gun, Saitama 350-0495, Japan.

## Abstract

Extracellular vesicle (EV) has been investigated for use in clinical testing in recent years. Specific EV surface proteins provide distinguishing characteristics, but are insufficient for more detailed classification of EVs. Here, we suggest a novel “Bimolecular surface antigen expressed in EV” (BiEV) as a potential indicator for more efficient EV screening. A BiEV can be identified using a previously developed method, enzyme-mediated activation of radical sources, to label the components proximal (within 20 nm) to a given molecule. We examined the screening of cancer cell-secreted EV (cEV) included in serum EVs from a model mouse for lung cancer. The cEV-specific BiEVs were first identified by through a comparison of serum EVs from wild-type and lung cancer mice, showing that the CHL1-SLC4A1 bimolecule was a significant candidate for cEV-specific BiEV. Enzyme-linked immunosorbent assay quantification of CHL1-SLC4A1 BiEV appeared to suggest a potential for cancer screening of these mice. Using the same protocols, we found that CHL1-caspase 14 BiEV was significantly elevated in lung cancer patients compared with healthy persons. A BiEV strategy may be able to make a contribution to more effective EV screening, resulting in novel biological and clinical applications.

## Introduction

Extracellular vesicle (EV) is lipid membrane vesicles 40 to 1000 nm in size secreted extracellularly by various cells. They are classified into several types based on their properties, secretory processes, and size [1–3]. Exosomes measuring approximately 50–150 nm and secreted through a multivesicular endosome contribute to not only the secretion of intracellular molecules, but also the transmissive mechanism that transports them to other cells for intercellular communication [1–3]. Because EVs are secreted from various cells and possess a variety of properties, it is important to distinguish among them. Serum EV screening in particular is a valuable clinical tool and provides an alternative to serum proteins secreted from cancer cells. As a result, attempts are being made to diagnose various diseases by classifying and detecting disease-specific EVs [4]. To our knowledge, the most recent advanced strategy involves the detection of non-coding RNAs (*e.g.*, miRNAs) in EVs [5], which may be able to detect large numbers of cancers with fairly high accuracy. Similarly, specific EV surface proteins expressed in these EVs (generally membrane proteins and/or membrane-attached proteins) have been investigated as targets for cancer screening [6]. Their potential to be used as indicators for the characterization of EVs has been investigated using single surface proteins. However, many of these single proteins, including marker proteins for EVs, are known to be ubiquitous proteins, and as such may be insufficient for distinguishing and classifying EVs.

Cell membrane or membrane-attached proteins non-randomly form heterocomplexes accompanied by a fluid biological membrane known as a “lipid raft” structure [7]. We reported that the membrane heterocomplex “<u>B</u>imolecular <u>C</u>ancer <u>T</u>arget” (BiCAT) could be a potential lung cancer (LC) antigen consisting of a hetero bimolecular complex [8]. In the BiCAT strategy, because cancer specific-antigens are defined by two molecules, the specificity is theoretically higher than that of a single-molecule antigen. Because EVs are composed of a lipid bilayer biological membrane, a heterocomplex, including a bimolecule, was considered to be an attractive indicator antigen to distinguish EVs.

To employ bimolecules in EV screening, it is essential to identify which molecules form the specific bimolecules in EVs. Proximity labeling [9,10] have recently attracted research resources to analyze these bimolecules. Among them, we developed a simple and physiological method, called enzyme-mediated activation of radical source (EMARS) [11,12], that labels only bimolecular partner proteins with fluorescein followed by analysis using an antibody array and/or typical proteome strategy [13]. To date, EMARS has been applied to various studies involving molecular complexes on cell membranes [14–20].

Here, we propose a “<u>Bi</u>molecular surface antigen expressed in <u>EV</u>” (BiEV) as a new type of indicator to provide diversity in EV screening. Transgenic mice with *EML4-ALK* onco-fusion genes [21,22] were used as an animal model for the spontaneous onset of LC. We focused on BiEVs in cancer cell-secreted EVs (cEVs) of this mouse model, which includes exosomes and ectosomes, raised from several LC cell pathways resulting in secreted to serum. In addition, serum EVs derived from 12 serum samples of healthy persons and LC patients were purified and used for BiEV analysis by the same protocols. The experiments revealed specific BiEV antigens in both mouse and human LC cell-secreted EVs, suggesting that BiEV in cEVs can serve as indicators in EV screening and biomarkers for LC screening.

## Material and Methods

### Animals

C57BL/6J mice used as wild type mice were obtained by self-breeding of the pair mice purchased from Japan CLEA Co. (Tokyo, Japan), and were maintained under specific pathogen-free conditions (SPF) at 20-25 °C in accordance with the Saitama Medical University Animal Experiment Committee (protocol number 30M059). The *EML4-ALK* transgenic mice [22] (generous gift from Professor Hiroyuki Mano and Dr. Manabu Sone, National Cancer Center Research Institute) were maintained under SPF conditions at room temperature in accordance with Saitama Medical School Animal Experiment Committee (protocol number 2697 and 2701) and the Saitama Medical University Safety Committee for Recombinant DNA Experiments (protocol number 1356). All animals were maintained under a 12 hour light/12 hour dark exposure and free access to food and water.

### Human serum samples

The serum samples of healthy persons and lung cancer patients were obtained from the Medical Genome Center Biobank, National Center for Geriatrics and Gerontology and BioBank Japan Project supported by Japan Agency for Medical Research and Development (AMED) [23,24] in Japan. The handling of human serum and its use in research are performed in accordance with the Declaration of Helsinki (as revised in Fortaleza, Brazil, October 2013), and the permission and the guidelines of the Ethics Review Committee of Saitama Medical University (protocol number 850).

### Cell culture and EV isolation

The LK2 human squamous cell lung carcinoma cell lines (RIKEN CELL BANK, Ibaragi, Japan) were cultured in RPMI 1640 medium (Wako Chemicals, Osaka, Japan) supplemented with 5% fetal bovine serum (FBS; GIBCO) at 37°C under humidified air containing 5% CO_2_. For the collection of secreted EVs from LK2 cells, the medium was replaced with new serum free medium, ASF Medium 104 (Ajinomoto, Tokyo, Japan), and then cultured cells continuously cells for 2 days. The secreted EVs were partially purified from cultured medium by using ExoQuick-TC Exosome Precipitation Solution (20 μl/100 μl serum; System Biosciences, CA) at 4°C for overnight. The precipitation containing EVs was solubilized with 100 μl PBS, and then used for further analysis.

### Preparation of HRP-conjugated antibody for EMARS reaction

The mouse and human anti-CHL1 antibodies (AF2147 and MAB2126; R&D systems, MN) were partially reduced and bound to HRP using a peroxidase labeling kit SH (Dojindo, Kumamoto, Japan). The binding capacity of the HRP-labeled antibody was evaluated as described previously [8].

### Purification and EMARS reaction for serum EVs

The serum EVs from mouse and human serum were purified by using the combination of polymer precipitation and size exclusion chromatography. In order to determine which fractions should be collected for EV purification in size exclusion chromatography, mouse serum and blue dextran (Sigma-Aldrich Japan, Tokyo, Japan) were subjected to the short column with 2 ml Sephacryl S-500 resin (GE Healthcare Life Sciences, IL) equilibrated with PBS. The resins were pre-treated with yeast extract solution (0.1 g/100 ml H_2_O for 1 hour; BD Biosciences, NJ) to avoid non-specific binding of EVs to resins. Then, the sample fractions were eluted with 100 µl PBS/fraction. The protein concentration of each fraction was measured using micro BCA protein assay kit (Thermo Fisher Scientific, MA) with VarioSkan Flash microplate reader (Thermo Fisher Scientific, MA) at 562 nm. The blue dextran was measured by with VarioSkan Flash microplate reader at 300 nm. The serum (50-100 μl) was treated with ExoQuick Exosome Precipitation Solution (25.2 μl/100 μl serum; System Biosciences, CA) on ice for 30 min. The precipitation (hereinafter referred to as precipitated EV) was solubilized with 100 μl PBS, and then added 0.25 µg of HRP-conjugated anti-mouse or human CHL1 antibody with gentle mixed. After incubation at room temperature for 20 min, the precipitated EV mixture was subjected to short column with 2ml Sephacryl S-500 equilibrated with PBS. The step-by-step elution was carried out under gravity flow with PBS (100 μl PBS/fraction). The fractions No. 6 to 8, which contained EVs with HRP-conjugated CHL1 antibody, were collected depend on each experiment, and applied to Nanosep® centrifugal devices (30K; Pall Life Sciences, MI) followed by centrifugation at 8,000 g for 5 min to concentrate EV particles. The concentrated EV particles on the filters were washed with 100 μl PBS once by centrifugation, and then fluorescein-tyramide (FT) reagent as EMARS reaction reagent [8] was added to the filter with gentle mixing. After 10 min incubation at room temperature, remaining FT reagent was removed by centrifugation at 8,000 g for 5 min followed by wash with PBS once by centrifugation. The EV proteins after EMARS reaction on the filter were collected with 30 μl of 50 mM Tris-HCl (pH 7.4) containing 1% SDS by gentle pipetting and transferred to new tube. To remove excess FT reagent, the collected sample was subjected to mini column with approx. 1ml Sephadex G-50 (GE Healthcare Life Sciences, IL), which was equilibrated with 50 mM Tris-HCl (pH 7.4) containing 1% SDS. The eluted samples (approx. 30 μl) were subsequently used for the analysis of EMARS products, such as immunoprecipitation, western blot, and ELISA described below. In the case of direct detection of EMARS products on SDS-PAGE gel, the gel was directly subjected to ChemiDoc MP Image analyzer (BIO-RAD, CA) with the gel fluorescent detection mode (blue light filter for fluorescein). For loading controls, the gel after exposure was stained with Coomassie Brilliant Blue solution.

### Purification and Enrichment of EMARS products

EMARS products were purified and enriched with anti-fluorescein antibody-conjugated Sepharose as described previously [8]. Briefly, the flow-through fraction of G-50 column described above was diluted with 300 μL of NP-40 lysis buffer. Then, 20 μl of prepared anti-fluorescein antibody Sepharose, which was produced by the conjugation reaction between anti-fluorescein antibody (600101096; Rockland Immunochemicals, PA or 6400-01; Southern Biotech, AL) and NHS-activated Sepharose 4 Fast Flow (GE Healthcare Life Sciences, IL), was added to the sample, and mixed with rotation at 4°C for overnight. After washing resins, appropriate solutions (reducing SDS-sample buffer for western blot or 1 % SDS solution containing MPEX PTS reagent (GL Science, Tokyo, Japan) for MS analysis) was added to Sepharose beads, and heated at 95°C for 5 min to elute the fluorescein-labeled molecules from resins.

### Proteomic Analysis of EMARS products by using mass spectrometry analysis

Proteomic analysis in this study was performed by using nano liquid chromatography-electrospray ionization mass spectrometry (nano LC-ESI-MS/MS). In the case of proteomic analysis for BiEVs in mouse serum EVs, we received the technical support of MS analysis (Laboratory of Cytometry and Proteome Research, Nanken-Kyoten and Research Core Center, Tokyo Medical and Dental University). Briefly, the peptide samples of EMARS products derived from serum EVs were purified under same protocols as described above, and were injected into a nano-UHPLC system (Bruker Daltonics, MA). Mass analysis was performed on a maXis-4G-CPR (Bruker Daltonics, MA) mass spectrometer equipped with a nano-ESI source. An L-column ODS (0.1×150 mm, 3 μm particle size, CERI, Tokyo, Japan) was used, preceded by a Magic C18AQ UHPLC NanoTrap column (5 μm particle size, 200 Å pore diameter). The peptides were eluted from the column using a 10% to 35% acetonitrile gradient over 50 min. The eluted peptides were directly electro-sprayed into the spectrometer, with MS/MS spectra acquired in a data-dependent mode.

In the case of proteomic analysis for BiEVs in human serum EVs, we received the technical support of MS analysis (the Natural Science Center for Basic Research and Development (N-BARD), Hiroshima University). The peptide samples of EMARS products prepared as well as mouse samples were injected into an Ultimate 3000 RSLCnano (Thermo Fisher Scientific, MA). Mass analysis was performed on a LTQ Orbitrap XL mass spectrometer equipped with a nano-ESI source (Thermo Fisher Scientific, MA). A NIKKYO nano HPLC capillary column (3μm C18, 75 μm I.D.×120 mm; Nikkyo Technos) was used, preceded by a C18 PepMap100 column (300 μm I.D × 5mm; Thermo Fisher Scientific, MA). The peptides were eluted from the column using a 4% to 35% acetonitrile gradient over 97 min. The eluted peptides were directly electro-sprayed into the spectrometer, with precursor scan and data-dependent MS/MS scan.

The raw data was analyzed using Proteome Discoverer ver. 2.2 software (Thermo Fisher Scientific, MA) with the following parameters: Cys alkylation: iodoacetamide, Digestion: Trypsin, Species: Homo sapiens or Mus musculus, FDR (False discovery rate): strict 1%, relaxed 5%, respectively.

### Cryo-electron microscopy

For cryoEM imaging of EVs, the grids were prepared by applying 3 μl sample on glow-discharged holey carbon grids (Quantifoil Mo 2/2 200 mesh). The sample was ice-embedded using a LEICA EM GP2 (Leica Microsystems, Tokyo, Japan) at 100% humidity and 10 °C. Images were collected on an FEI Talos Arctica microscope (Thermo Fisher Scientific, MA) at 200 kV using a Falcon III detector in liner mode and a Volta phase plate. Each data was collected at a dose rate of 10e^−^/pixel/sec for 4 s, with a total accumulated dose of ~ 40 e^−^/Å^2^.

### Western blot analysis

For western blot analysis, the solubilized samples (purified EVs etc.) with reducing SDS sample buffer or the eluted samples from purified and enriched resins described above were subjected to 6 to 12 % SDS-PAGE by using typical SDS-running buffer or Rapid Running Buffer Solution (Nacalai Tesque, Kyoto, Japan) in order to adjust to molecular mass of each candidate protein. The gels were subsequently blotted to an Immobilon®-P PVDF Membrane (Millipore, MA). After blocking with 5% skim milk solution, transferred membrane was reacted with first antibodies as follows: anti-mouse and human CHL1 antibody (AF2147 and MAB2126; R&D systems, MN; 1 μg/ml), HRP-conjugated anti-mouse and human CHL1 antibody (described above), anti-α2 integrin antibody (ab133557; Abcam; 1 μg/ml), anti-β1 integrin antibody (610467; BD transduction laboratories, NJ; 0.25 μg/ml), anti-FGFR3 antibody (c-15; Santa cruz, TX; 0.2 μg/ml 5% BSA-PBS), anti-TSG101 antibody (EXOAB-TSG101-1; System Biosciences, CA; 1:500), anti-CD63 antibody (EXOAB-CD63A-1; System Biosciences, CA; 1:500), anti-CD5L antibody (AF2834; R&D systems, MN; 0.4 μg/ml), anti-Pregnancy Zone Protein (PZP) antibody (PAG324Ra01; CLOUD-CLONE, TX; 0.5 μg/ml), anti-SLC4A1 antibody (18566-1-AP; PROTEINTECH, IL; 0.6 μg/ml), anti-Thrombospondin 1 (THBS1) antibody (PAA611Hu01; CLOUD-CLONE, TX; 0.5 μg/ml), and anti-caspase 14 antibody (MAB8215; R&D systems, MN; 0.5 μg/ml). The reaction with first antibody was carried out at room temperature for 1 hour to overnight. Then, the appropriate secondary antibody, HRP-conjugated anti-mouse IgG (Promega, WI; 1:3000-5000), HRP-conjugated anti-rabbit IgG (Promega, WI; 1:3000-5000), HRP-conjugated anti-goat IgG (Santa Cruz, TX; 1:3000-5000), Rabbit or Mouse TrueBlot®: anti-Rabbit or Mouse IgG HRP (Rockland Immunochemicals, PA; 1:1000), or HRP-conjugated anti-rat IgG (Santa Cruz; 1:3000-5000) were reacted at room temperature for 1 hour. After antibody treatment, the membranes were developed with an Immobilon Western Chemiluminescent HRP Substrate (Millipore, MA). The membrane was exposed and analyzed using ChemiDoc MP Image analyzer (BIO-RAD, CA). The molecular weight marker was a Pre-stained Protein Markers, Broad Range (Nacalai Tesque, Kyoto, Japan). For loading controls, SDS-PAGE gel was stained with Coomassie Brilliant Blue solution. In the case of membrane stripping, the membranes were treated with stripping solution (Wako Chemicals, Osaka, Japan) at room temperature for 20 min, re-blocking, and re-stained with appropriate antibody.

### ELISA for CHL1 proteins in serum or EVs

The mouse serum CHL1 level was measured with direct ELISA method using HRP-conjugated anti-mouse and human CHL1 antibody as described previously [25]. In the case of the human CHL1 level in serum and serum EVs by sandwich ELISA, diluted serum or precipitated serum EVs described above were applied to the ELISA plates after coated with capture antibody, anti-human CHL1 antibody (10143-MM05; Sino Biological, Beijing, China), followed by the detection with HRP-conjugated anti-human CHL1 antibody (described above). The relative index (CHL1) for the measurement of mouse serum CHL1 levels were determined based on the value of standard sample described previously [25]. In human CHL1 ELISA, the recombinant human CHL1 partial protein (10143-H08H; Sino Biological, Beijing, China) was used as a standard.

### ELISA for fluorescein-labeled proteins

For the sandwich ELISA to measure fluorescein-labeled SLC4A1 or caspase 14, 96 well ELISA plate (#3369; CORNING) was coated with anti-mouse and human SLC4A1 antibody (18566-1-AP; PROTEINTECH, IL; 1 μg/ml PBS) or anti-human caspase 14 antibody (H00023581-M01; Abnova, Taipei, Taiwan; 0.6 μg/ml PBS) at 4 °C for overnight. After blocking with 1% BSA-PBS (or 5% skim milk-PBS in caspase 14) solution at 37 °C for 1 hour, the corresponding EMARS products containing fluorescein-labeled SLC4A1 or fluorescein-labeled caspase 14 were added and incubated at 37 °C for 1 hour. Each well was washed gently with PBST (0.05 % tween-PBS solution), then treated with HRP-conjugated anti-fluorescein antibody (1 μg/ml), which was prepared from anti-fluorescein antibody (600101096; Rockland Immunochemicals, PA) using a peroxidase labeling kit NH_2_ (Dojindo, Kumamoto, Japan) according to the manufacture’s instruments, at 37 °C for 1 hour. In the case of the detection of SLC4A1 molecule itself, recombinant human SLC4A1 poly peptides (CSB-EP021663HU; CAUABIO THECHNOLOGY, Wuhan, China) and HRP-conjugated anti-SLC4A1 antibody, which was prepared using a Zenon rabbit IgG HRP Labeling Kit (Thermo Fisher Scientific, MA) according to the manufacturer’s instrument, was used. The development was carried out with SureBlue/TMB Peroxidase Substrate (100 μl /each well; Sera care) at room temperature for appropriate minutes (3-10 min). After stopping the reaction using 2M HCl solution, the absorbance (O.D. 450 nm) was measured using Varioskan Flash microplate reader (Thermo Fisher Scientific, MA). The standard sample of fluorescein-labeled SLC4A1 (100 μl) was prepared as the EMARS products derived from 100 μl serum in large tumor-bearing *EML4-ALK* transgenic mouse (30 weeks old male). To compensate the differences among ELISA plates, the relative values were determined based on the value of standard sample as “BiEV index”. The “BiEV index (SLC4A1)” used in this study showed how many microliters corresponds to the above standard sample. In the case of fluorescein-labeled caspase 14, the “BiEV index (caspase 14)” was used based on the fluorescein-labeled recombinant caspase 14 protein (11856-H07E; Sino Biological, Beijing, China) labeled with NHS-fluorescein (Thermo Fisher Scientific, MA). Caspase 14 protein (2.5 μg) and 8 μg NHS-fluorescein were mixed into 20 μl PBS, and then incubate at room temperature for 50 min. The reaction mixture was applied to mini column with approx. 0.5 ml Sephadex G-50 (GE Healthcare Life Sciences, IL) in order to remove excess NHS-fluorescein.

### ELISA for CHL1 proteins in serum or EVs

The mouse serum CHL1 level was measured with direct ELISA method using HRP-conjugated anti-mouse and human CHL1 antibody as described previously [25]. In the case of the human CHL1 level in serum and serum EVs by sandwich ELISA, diluted serum or precipitated serum EVs described above were applied to the ELISA plates after coated with capture antibody, anti-human CHL1 antibody (10143-MM05; Sino Biological, Beijing, China), followed by the detection with HRP-conjugated anti-human CHL1 antibody (described above). The relative index (CHL1) for the measurement of mouse serum CHL1 levels were determined based on the value of standard sample described previously [25]. In human CHL1 ELISA, the recombinant human CHL1 partial protein (10143-H08H; Sino Biological, Beijing, China) was used as a standard.

### Statistical analysis

The statistical analyses and receiver operating characteristic (ROC) curve analysis for the data of ELISA measurement were performed using R software (The R Foundation for Statistical Computing, Austria) and EZR (Saitama Medical Center, Jichi Medical University, Japan) [26], which is a graphical user interface for R. Statistical significance test of BiEV index of fluorescein-labeled SLC4A1 and caspase 14 were performed with Mann–Whitney test. We used a statistical significance level of 0.05 or smaller. Similarly, the correlation between tumor mass and BiEV index (SLC4A1) was indicated using Spearman’s rank correlation coefficient.

## Results

### Purification of serum EVs from *EML4-ALK* transgenic mice

A typical BiEV analysis of serum EVs is summarized in Figure 1. Western blot analysis of CD63 confirmed that EVs could be crudely purified from mouse serum by a polymer precipitation method (Figure 2A). CHL1, which is the molecule constituting LC BiCAT [8], was also detected at higher levels in serum EVs from *EML4-ALK* transgenic mice (Figure S1A). EMARS products from precipitated serum EVs (crude EVs) were detectable in CHL1 probe (+) samples, but not in EMARS probe (−) samples, indicating that EMARS reacts well with CHL1 probes and no serum factors induced non-specific EMARS reactions (Figure 2B). However, fluorescein-labeled EMARS products contained so many serum proteins (Data not shown) that size-exclusion chromatography was subsequently performed. The No. 6 to 8 fractions, fractionated by Sephacryl S-500 resin, were most suitable for collection of serum EVs with limited serum proteins (Figure 2C). The detection of TSG101 and ubiquitinated TSG101 [27], a typical EV marker, supported a similar conclusion (Figure S1B). Cryo-electron microscopy revealed many spherical particles measuring dozens of nanometers in size, with a lipid bilayer in fraction No. 6 and 8 samples (Figure. 2D), with each fraction containing EVs approximately 20– 100 nm in diameter.

**Figure 1.**
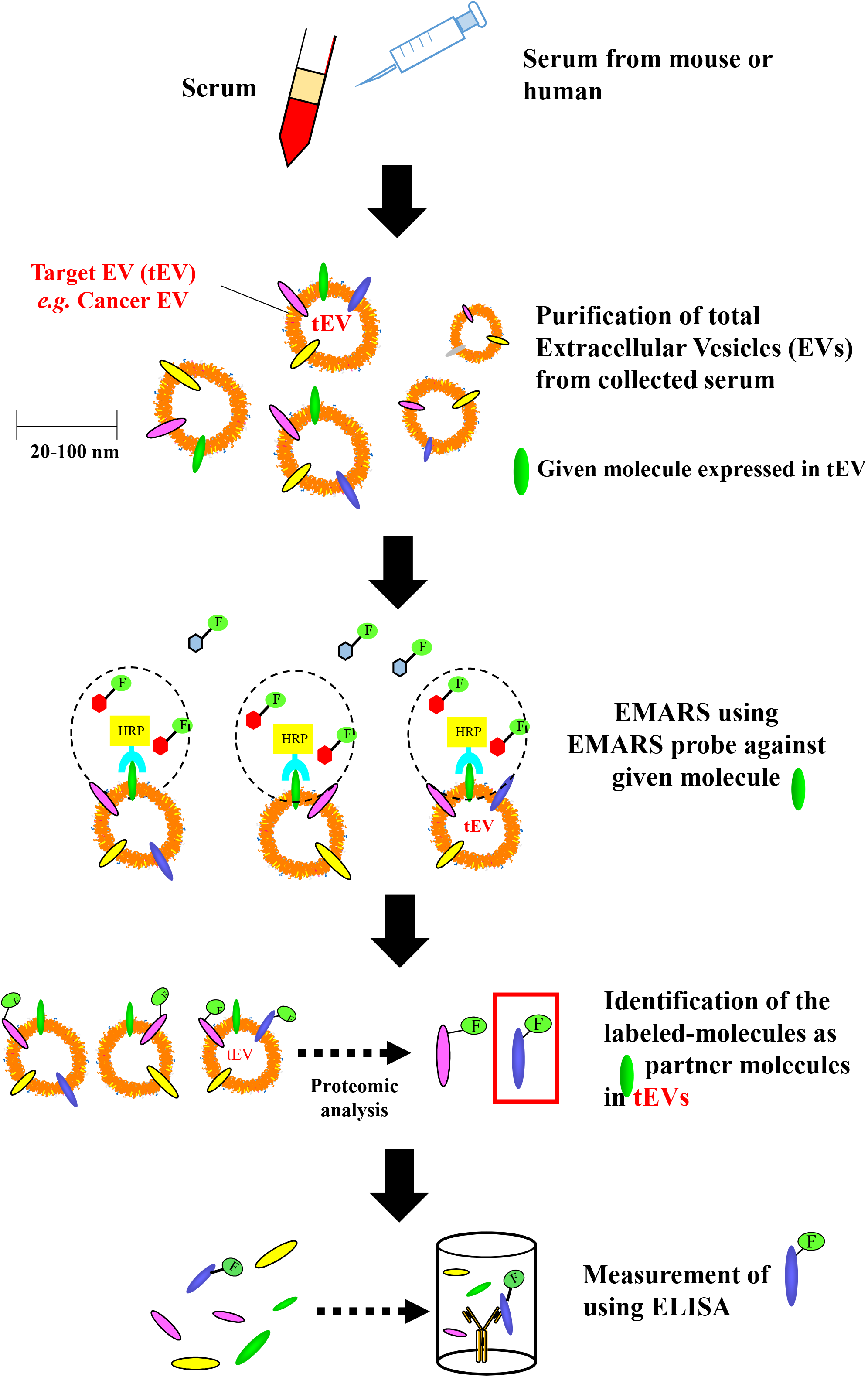
Overview of BiEV analysis for serum EVs. Schematic illustration of BiEV analysis for serum EVs. The target EV (tEV) indicates the EV to be detected and investigated. Before the EMARS reaction, serum EVs were purified using a combination of polymer-based precipitation and size-exclusion chromatography. To determine the amount of tEV in the sample by measuring the BiEV levels, appropriate BiEV partner molecules labeled with fluorescein were measured by a sandwich ELISA.

**Figure. 2.**
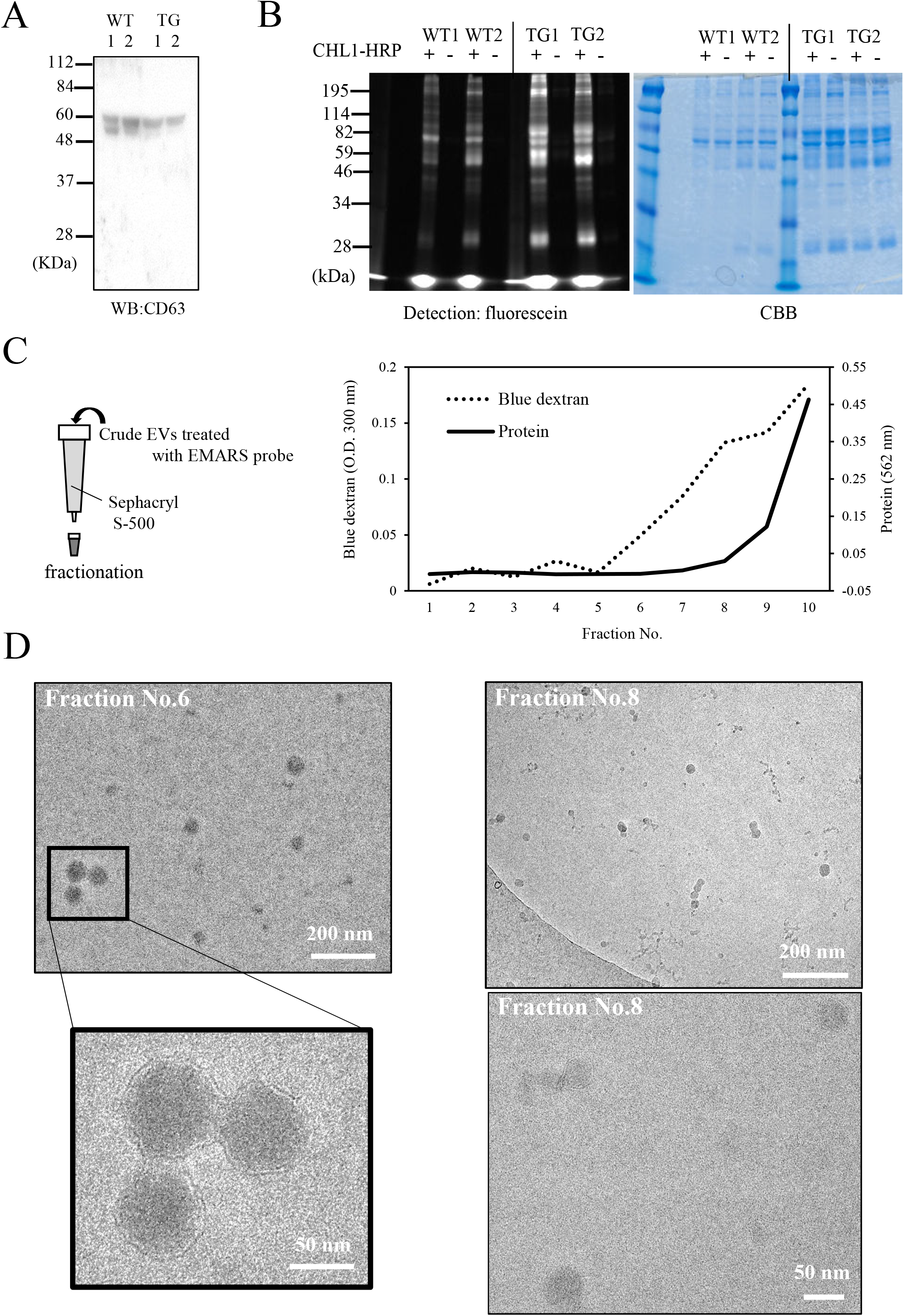
Purification of EVs from serum. (*A*) Western blot analysis of CD63 EV marker in precipitated EV samples. The precipitated EVs derived using ExoQuick Solution were prepared from the serum collected from 2 mice of wild-type (WT) and *EML4-ALK* transgenic (TG) mouse as described in the “*Materials and methods*”. (*B*) Analysis of EMARS products obtained from EMARS reaction for crude mouse serum EVs. EMARS reaction was carried out directly in the precipitated serum EVs from two of WT and TG mice with or without HRP-conjugated CHL1 probe. The EMARS products were subsequently subjected to SDS-PAGE (10% gel) with fluorescein detection and CBB staining. (*C*) Fractionation using Sephacryl S-500 chromatography. Blue dextran (an indicator of void volume) and mouse serum (an indicator of protein elution) were used for preliminary experiments. The dotted line indicates absorbance of blue dextran. The solid line indicates protein concentration measured using a BCA protein kit. (*D*) Morphological observation of serum EVs (fraction No. 6 and No. 8) using cryo-electron microscopy. Lower panel of fraction No.6 is an enlarged view of a part of the upper panel. Scale bar; 200 nm (upper panel) and 50 nm (lower panel).

### EMARS reactions for purified serum EVs from *EML4-ALK* transgenic mice

We first classified sera from three groups: wild-type (WT) mice, *EML4-ALK* transgenic mice with a large tumor (TL; tumor weight above 0.15 g), and *EML4-ALK* transgenic mice with a small tumor (TS; tumor weight 0.1 g or less). To obtain experimental results averaged over some mice, a 10 μL aliquot of the serum from 10 animals in each group was mixed in equal proportions, and then used for the following experiments. EMARS products were detected in the sample from TL mice in contrast to WT and TS mice (Figure 3A “IP”). The band pattern of TS samples was similar to that of the WT sample (Figure 3A), indicating that cancer EVs expressing CHL1 were poorly contained in TS compared with TL. TSG101 and CD63 molecules were moderately abundant in the TL sample compared with the other two samples (Figure S2A and S2B), suggesting that the signals of EMARS products in the TL sample were due to a moderate increase in CHL1-expressed cEVs secreted from large tumor tissues. This supports the fact that fluorescein-labeled CHL1 molecules raised from self-labeling by EMARS with the CHL1 probe were abundant in TL (Figure S2C).

**Figure 3.**
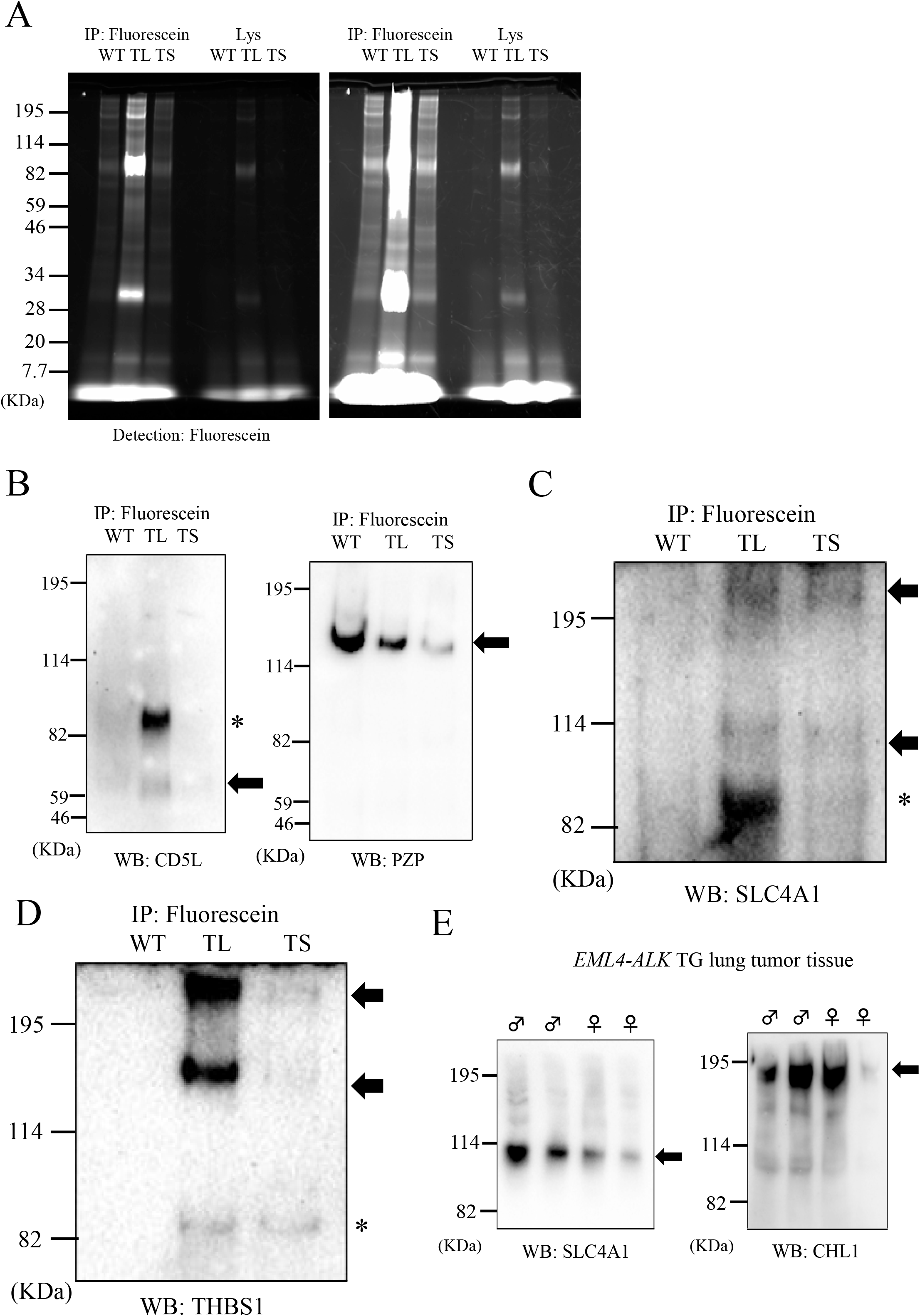
BiEV analysis for serum EVs from *EML4-ALK* transgenic mouse. (*A*) The EMARS products for serum EVs from *EML4-ALK* transgenic mouse. To average experimental results over each group, an aliquot of the serum (10 μL each) from 10 animals in each group (WT), large lung tumor-bearing (TL), and small lung tumor-bearing mice (TS) was mixed in equal proportions, and then applied to EV purification and EMARS. The EMARS products were concentrated and purified by immunoprecipitation with the anti-fluorescein antibody Sepharose. The resulting samples were subjected to SDS-PAGE analysis with fluorescence detection. “IP” indicates the immunoprecipitated samples, and “Lys” indicates the lysate samples before immunoprecipitation. The right panel indicates the same gel as the left panel, but exposed for a longer time. (*B*) Confirmation of candidate partner molecules (CD5L and PZP) with mouse CHL1 in EVs. The EMARS products of WT, TL, and TS were respectively applied to immunoprecipitation (anti-fluorescence antibody Sepharose) and western blot analysis with anti-CD5L (left panel) antibody. After the stripping as described in the “*Materials and methods*”, the membranes were re-stained with anti-PZP antibody (right panel). Arrows indicate the detected band of CD5L and PZP proteins (including predicted dimer). Asterisk indicates unknown bands (predicted as non-specific or partial fragments). (*C, D*) Confirmation of candidate partner molecules (SLC4A1 and THBS1) with mouse CHL1 in EVs. The western blot analysis was performed with anti-SLC4A1 antibody (*C*). After stripping, the membranes were re-stained with anti-THBS1 antibody (*D*). Arrows indicate the detected band of SLC4A1 and THBS1 proteins (including predicted dimers). Asterisks indicate unknown bands (predicted as non-specific or partial fragments). (*E*) Expression of SLC4A1 (left column) and CHL1 proteins (right column) in tumor tissues from two male and two female *EML4-ALK* transgenic mice. The fragments of lung cancer tissues were mashed and washed gently with PBS, and then lysed with SDS-PAGE sample buffer directly. The resulting samples were subjected to Western blot analysis with anti-SLC4A1 antibody and anti-CHL1 antibody. Arrows indicate the detected band of monomer SLC4A1 and CHL1 proteins.

### Identification of partner molecules of CHL1 in serum cEVs

In small tumor-bearing mice, specific BiEVs in serum cEVs may be difficult to detect due to the small amounts of labeled molecules. Candidates of BiEV partners of CHL1 suitable for small tumor screening in *EML4-ALK* transgenic mouse were examined by mass spectrometry (MS) proteomics, using both TL and TS EMARS products, as summarized in Table 1 (with raw data provided in Table S1). We selected four candidate molecules (CD5L, PZP, THBS1, and SLC4A1). Western blot analysis using anti-CD5L antibodies revealed moderate bands at approximately 60 kDa in the TL samples, but no obvious bands were detected in the other two samples (Figure 3B; left panel). After stripping, the same membrane was re-stained with anti-PZP antibodies. PZP (approximately 130 kDa) was detected strongly in the WT sample (Figure 3B; right panel). The results indicate that some CHL1 BiEVs were also expressed in the serum EVs from WT mice. In contrast, western blot analysis using anti-SLC4A1 and anti-thrombospondin-1 (THBS1) antibodies revealed moderate bands in both TL and TS samples, but not in WT samples (Figure 3C and 3D). In addition, we found that SLC4A1 and CHL1 molecules were expressed in LC tissue from *EML4-ALK* transgenic mice (Figure 3E). We therefore postulated that the measurement of CHL1-SLC4A1 BiEV in serum EVs may be an effective indicator for cEV screening in *EML4-ALK* transgenic mice.

**Table 1.**
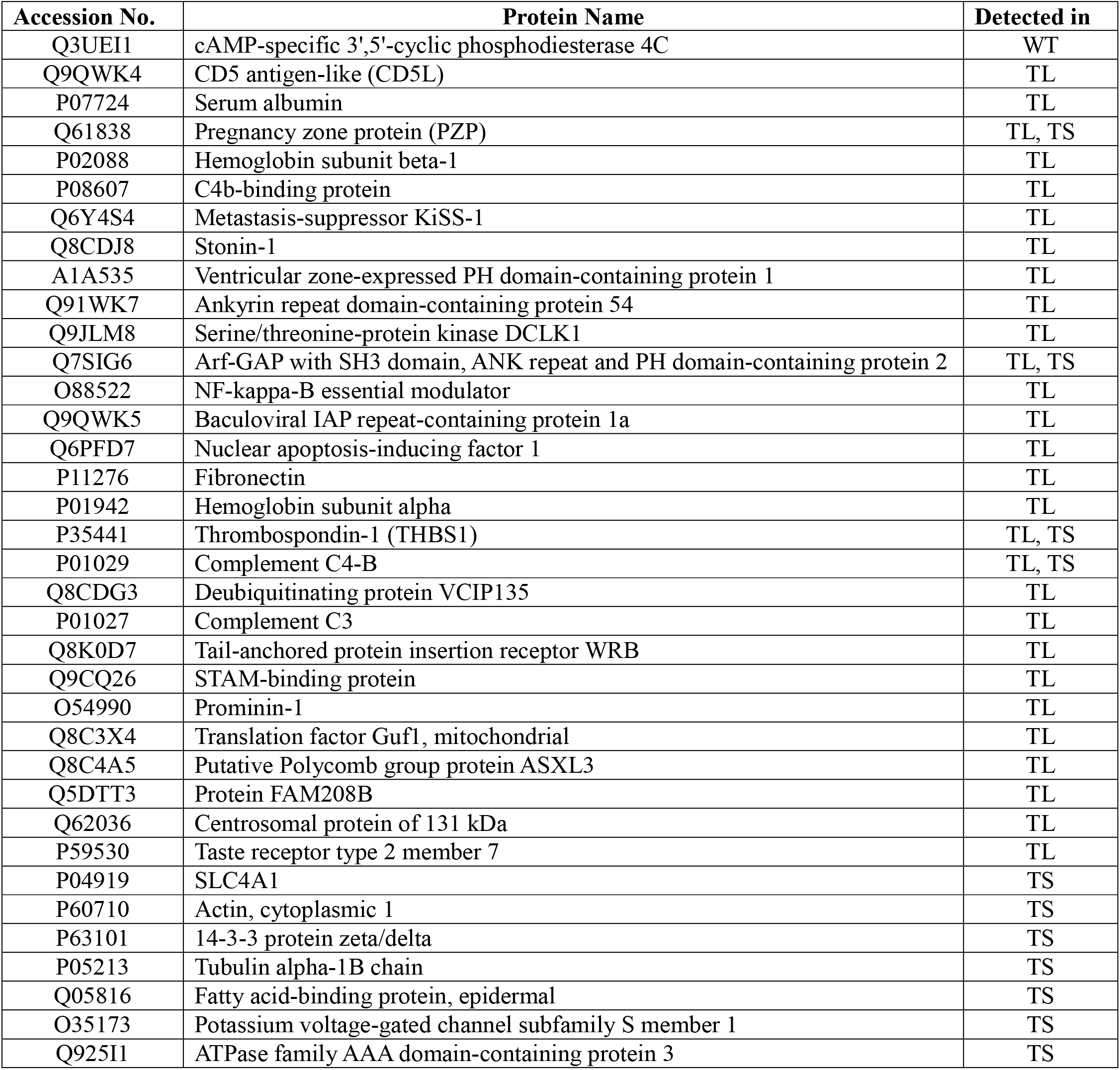
Candidates for the partners of CHL1 in serum EVs from WT, TL, and TS mice.

### Measurement of CHL1-SLC4A1 BiEVs using a sandwich ELISA

The serum CHL1 level has been identified as a significant indicator of somewhat larger lung tumor-bearing *EML4-ALK* transgenic mice, however, it was an insufficient indicator for small lung tumor-bearing *EML4-ALK* transgenic mice [25]. To determine whether the CHL1-SLC4A1 BiEV in serum EVs was elevated in *EML4-ALK* transgenic mouse, and TS mice in particular, measurement of fluorescein-labeled SLC4A1, which indicated an abundance of CHL1-SLC4A1 BiEV, was performed using a sandwich enzyme-linked immunosorbent assay (ELISA). Using recombinant SLC4A1 we determined that our ELISA could perform sensitive measurements of SLC4A1 (at least approx. 300 pg to 2000 pg/mL with linearity) as indicated by a correlation coefficient of 0.9995 in the calibration curve (Figure S3A). However, fluorescein-labeled SLC4A1, which was made by a fluorescein-labeling reagent, did not perform well due to an unknown mechanism. We used EMARS products containing moderate amounts of fluorescein-labeled SLC4A1 obtained from one TL mouse serum EV as a standard. It was measured using horseradish peroxidase– conjugated anti-fluorescein antibodies (with a correlation coefficient of 0.995; Figure S3B). To compare the results of each ELISA experiment in different plates, a BiEV index (SLC4A1) was based on the measured value of the standard substance (Figure S3B).

The BiEV index (SLC4A1) from WT mice and *EML4-ALK* transgenic mice (detailed information for each is provided in Table S2) is summarized in Figure 4A. Whereas CHL1 serum levels of small tumor-bearing mice were not significantly different from that of WT mice (Figure S3C), the BiEV index (SLC4A1) in small tumor-bearing mice (TS) had a tendency to be higher overall compared with that in WT mice (Figure 4A). A significant difference was observed in the mean value of CHL1-SLC4A1 BiEV levels between WT and small tumor-bearing mice (*p* value: 8.9 × 10^−3^). Based on the receiver operating characteristic (ROC) curve of each BiEV index (SLC4A1), the area under the curve (AUC) and cut-off value were calculated as 0.782 and 0.238, respectively (Figure 4B). In addition, a Spearman’s rank correlation analysis indicated a slight correlation between the BiEV index (SLC4A1) and tumor mass (Figure 4C), suggesting that cEVs expressing CHL1-SLC4A1 BiEV were not secreted as much in the small tumor state, resulting in a moderate proportional correlation with tumor size. In contrast to increased CHL1-SLC4A1 BiEV in small tumor-bearing mice, a decrease in averaged SLC4A1 protein expression levels in whole-serum EVs was observed (Figure 4D).

**Figure 4.**
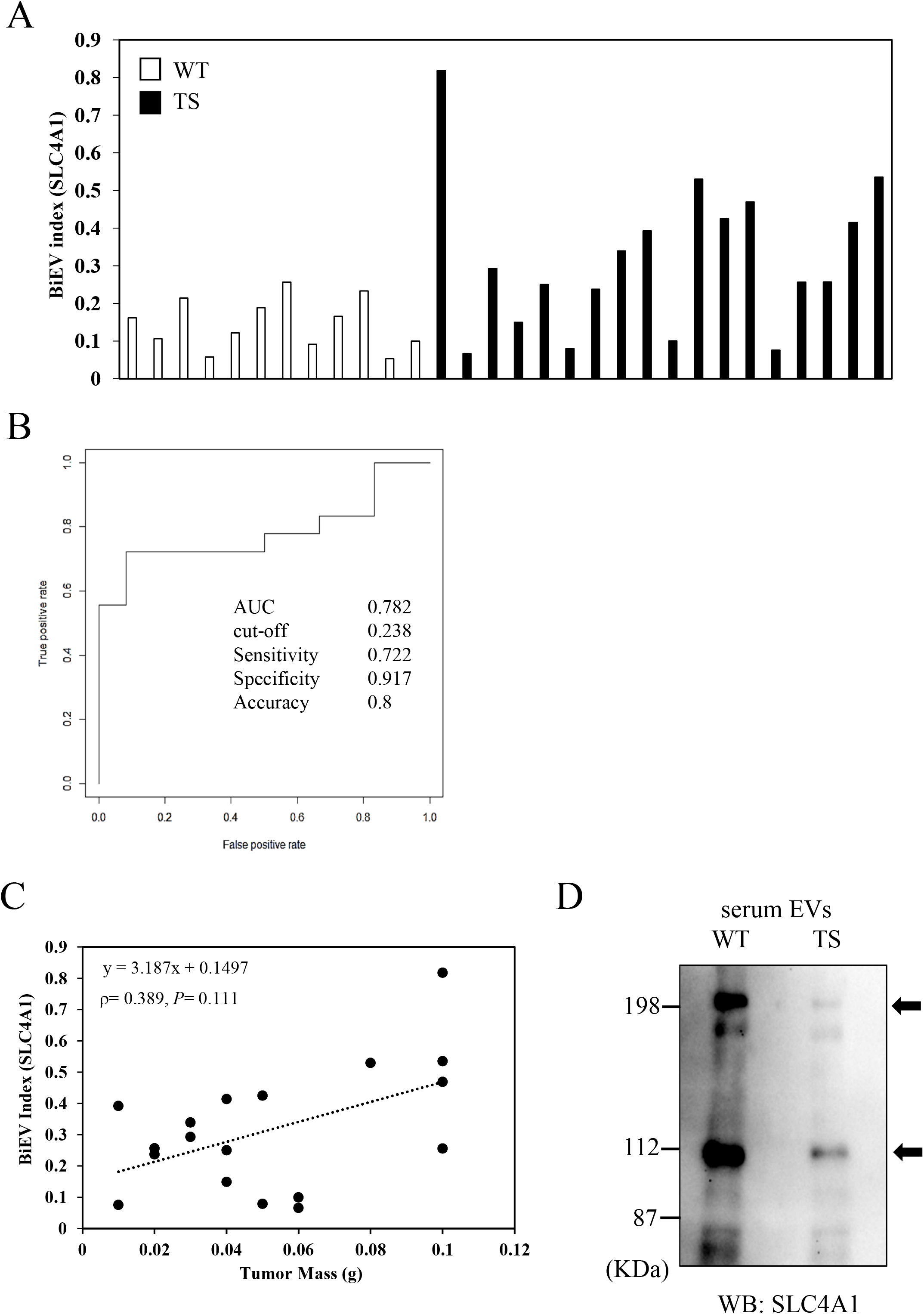
Measurement of CHL1-SLC4A1 BiEV in serum EVs. (*A*) Measurement of fluorescein-labeled SLC4A1 using a sandwich ELISA. The serum EVs from 12 wild-type mice (open bar) and 18 small lung tumor-bearing *EML4-ALK* transgenic mice (closed bar) were applied to EMARS reactions followed by ELISA measurements, respectively. The EMARS products containing fluorescein-labeled SLC4A1 were added to anti-SLC4A1 antibody-coated ELISA plates. A BiEV index (SLC4A1) was calculated based on the value of fluorescein-labeled SLC4A1 in standard samples. (*B*) ROC curve for BiEV indexes. AUC was calculated as 0.782. (*C*) A scatter-plot of tumor mass versus BiEV index. Correlation between BiEV index and lung tumor mass of TS was estimated using Spearman’s rank correlation coefficient (0.389: *p* = 0.111). (*D*) Western blot analysis of SLC4A1 in whole-serum EVs from WT and TS. An aliquot of the serum from 12 animals in the wild-type group and 18 animals in small lung tumor-bearing *EML4-ALK* transgenic mice was mixed in equal proportions followed by EV purification. Arrows indicate the detected band of SLC4A1.

### BiEV analysis and measurement in lung cancer patients

Because BiEV was found to be useful for screening serum EV in mice, a similar analysis was performed using serum EVs from healthy person (H) and LC patients. To examine whether CHL1 alone is a significant difference between H and LC patients, serum CHL1 levels or CHL1-expressing serum EVs were measured by ELISA (a correlation coefficient of 0.9878 in a calibration curve: Figure S4A) using a previously described technique [25]. In both measurements, no significant differences between H and LC were found (Figure S4B and S4C), suggesting that BiEV analysis may be needed as well as *EML4-ALK* transgenic mice.

As with the model mouse experiments, an aliquot of the serum from healthy person and LC patients (detailed information is provided in Table S3) was mixed in equal proportions, and then used for the following experiments. EMARS products using CHL1 EMARS probe were detected in both serum EVs, although the band pattern seemed to be no critical differences between H and LC samples because of only small amount of cEV in LC sera, as with the WT and TS mice (Figure 5A). To determine whether fluorescein-labeled SLC4A1 as identified in the model mouse experiments could be detected in human LC samples, EMARS products from H and LC were subjected to SLC4A1 western blot analysis and ELISA quantification. No crucial differences between H and LC (Figure S5A and S5B) were seen, indicating that identification of more suitable BiEV should be needed in the human samples. The candidates of BiEV partners of human CHL1 were identified by MS proteomics (summarized in Table 2, with raw data provided in Table S4 and a summary of mass scores in Table S5).

**Table 2.** Candidates for the partners of CHL1 in serum EVs from lung cancer patients.

| Accession No. | Protein Name | Also detected in LK2 EVs* |
| --- | --- | --- |
| P15924 | Desmoplakin | ○ |
| P02788 | Lactotransferrin | ○ |
| P31947 | 14-3-3 protein sigma | ○ |
| P04792 | Heat shock protein beta-1 |  |
| P07355 | Annexin A2 | ○ |
| P27482 | Calmodulin-like protein 3 |  |
| P47929 | Galectin-7 | ○ |
| Q08554 | Desmocollin-1 | ○ |
| Q01469 | Fatty acid-binding protein 5 | ○ |
| P37802 | Transgelin-2 |  |
| P04083 | Annexin A1 | ○ |
| P06733 | Alpha-enolase |  |
| P09211 | Glutathione S-transferase P |  |
| Q13885 | Tubulin beta-2A chain |  |
| P63104 | 14-3-3 protein zeta/delta | ○ |
| P31944 | Caspase-14 | ○ |
| P68366 | Tubulin alpha-4A chain |  |
| P58107 | Epiplakin |  |
| P01833 | Polymeric immunoglobulin receptor | ○ |
| P68363 | Tubulin alpha-1B chain |  |
| P29508 | Serpin B3 | ○ |
| P68371 | Tubulin beta-4B chain |  |
| P60174 | Triosephosphate isomerase |  |
| Q9HCY8 | Protein S100-A14 |  |
| P31949 | Protein S100-A11 | ○ |
| Q96DA0 | Zymogen granule protein 16 homolog B | ○ |
| Q9NZT1 | Calmodulin-like protein 5 |  |
| P02545 | Prelamin-A/C |  |
| P68104 | Elongation factor 1-alpha 1 | ○ |
| P11142 | Heat shock cognate 71 kDa protein | ○ |

\* “LK2 EVs” were purified from the culture supernatant of LK2 lung cancer cell line which expresses CHL1.

**Figure 5.**
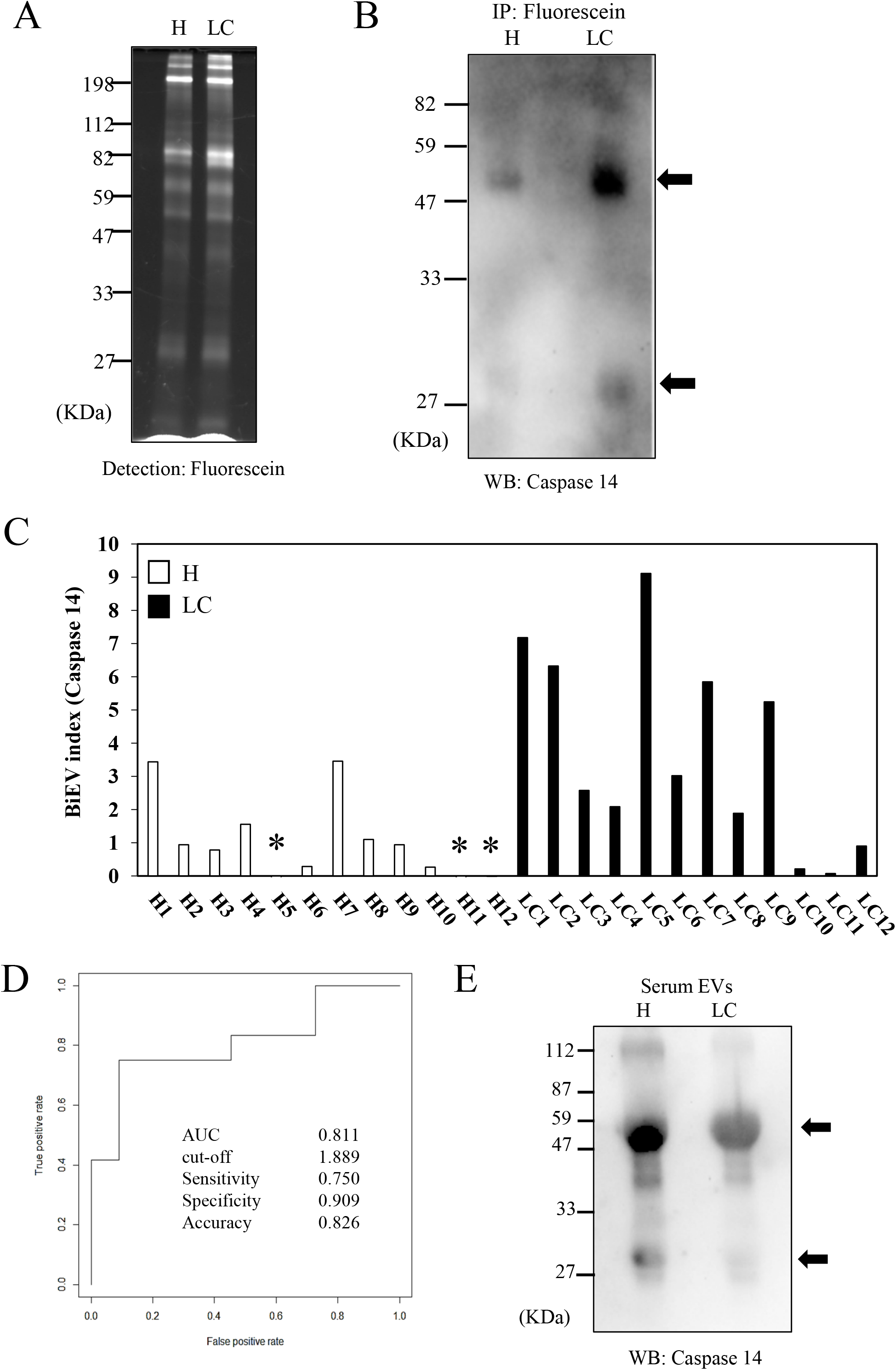
Measurement of CHL1-caspase 14 BiEV in serum EVs from lung cancer patients. (*A*) EMARS products purified from serum EVs of healthy person (H) and lung cancer (LC) patients. Fifty microliters of mouse serums was collected from the H and LC groups, and utilized in EV purification followed by EMARS reactions. To average experimental results over each group, an aliquot of the serum (10 μL each) from 5 H and 5 LC was mixed each in equal proportions. The EMARS products were subjected to SDS-PAGE analysis with fluorescence detection. (*B*) Confirmation of caspase 14 as a partner molecule with CHL1 identified by MS proteomics. The H and LC samples were applied respectively to immunoprecipitation (anti-fluorescence antibody Sepharose) and western blot analysis with anti-caspase 14 antibodies. Arrows indicate the detected band of caspase 14 proteins (including predicted dimer). (*C*) Measurement of fluorescein-labeled caspase 14 using a sandwich ELISA. Serum EVs from 12 H (open bar) and 12 LC (closed bar) were applied to EMARS reactions followed by ELISA measurements, respectively. The EMARS products containing fluorescein-labeled caspase 14 were added to anti-caspase 14 antibody-coated ELISA plates. “BiEV index (caspase 14)” was calculated based on the value of fluorescein-labeled recombinant caspase 14 made by fluorescein-labeling regent. The values are shown as the average of three independent ELISA experiments using the same samples. The detail data of H and LC persons is provided in Table S3. Asterisks indicate the samples were below detection limit. (*D*) ROC curve for BiEV indexes. The AUC was calculated as 0.811. (*E*) Western blot analysis of caspase 14 in whole-serum EVs from H and LC. An aliquot of the serum (2 μL each) from 12 persons in H and LC was mixed in equal proportions followed by EV purification with precipitation protocol. Arrows indicate the detected band of caspase 14.

In the first screening using immunoprecipitation and western blot analysis, several candidates were examined (Fatty acid-binding protein 5, Galectin 7, caspase 14) and revealed that caspase 14, which is family member of caspase involved in keratinocyte differentiation [28], was detected at significant levels as a partner of CHL1 BiEV in serum EVs from LC (Figure 5B) [29,30]. Fluorescein-labeled caspase 14 was also detected in EMARS products of secreted EVs from LK2 culture cells (Table 2), suggesting that CHL1–caspase 14 BiEV is a typical indicator of cEV in LC. We therefore postulated that the measurement of CHL1–caspase 14 BiEV in serum EVs is a suitable indicator for the cEV screening in LC.

To measure fluorescein-labeled caspase 14 in EMARS products using sandwich ELISA, a fluorescein-labeled caspase 14 standard was prepared using recombinant caspase 14. The caspase 14 ELISA system was suitable for sensitive measurement of the fluorescein-labeled caspase 14 standard with a correlation coefficient of 0.9878 in a calibration curve (Figure S5C). To compare the results among each ELISA experiment in different plates, a BiEV index (caspase 14) was set based on the measured value (ng/mL) of the fluorescein-labeled caspase 14 standard.

The BiEV index (caspase 14) from the H and LC serum EVs is summarized in Figure 5C. LC EVs had a tendency to be higher overall than that in H EVs. The mean value of the BiEV index (caspase 14) between the H and LC EVs was significantly different (*p* value: 0.019). Based on the ROC curve of each BiEV index, the calculated AUC and cut-off value were 0.811 and 1.889, respectively (Figure 5D). No increase in total protein expression levels of caspase 14 molecules in whole-serum EVs between H and LC serum EVs was found (Figure. 5E).

## Discussion

Taking into consideration the potential and expectation for bimolecular strategies identified by previous studies [8], we examine the possibility of applying BiEV to EV screening and clinical applications. Because several functional EVs, including exosomes, have been investigated as potential secretable biomarkers for precision medicine [31]. We focused on a characterization of serum EVs as a BiEV carrier to develop a new strategy for EV screening. It was once assumed that partner molecules of CHL1 in cell membranes and EVs were similar because EVs may contain BiCATs [8], which exist in cancer cell membranes as BiEVs. Despite the presence of three BiCATs in transport vesicles, as confirmed by electron microscopy [8], this was not the case with BiCATs in serum cEV (Figure S1A). This likely because the transport vesicles described in the BiCAT report were not involved in EV formation, or the membrane components constituting EVs in *EML4-ALK* transgenic mouse were originally different or altered cancer cell membranes.

We chose to independently examine BiEVs in serum cEV using EMARS. In an analysis of serum EVs, purification of serum EVs (Figure 2C) was an important sequence, because EMARS reactions labeled many serum proteins utilizing insufficient purification samples [32,33]. Although several purification methods are available for EVs, we selected size-exclusion chromatography [34] because of its simplicity of handling and low costs. Our gel-purification system was useful for collecting and purifying many EV particles (Figure 2D). However, in purified EVs, some soluble serum proteins were detected by the EMARS reaction. These serum proteins may be adsorbed by the EV itself in the bloodstream and/or in the EV secretion pathway from cancer cell and may be present in the proximity of CHL1. Whether these are simply adsorbed or have the property of adsorbing specifically to EV surface molecules is unknown.

A previous study has shown that CHL1 expression in serum EVs was increased in large tumor-bearing mice [25], and in this study, CHL1 BiEV was greatly increased in large tumor-bearing mice (Figure 3). This is the result of upregulated secretions of EVs expressing CHL1, increased expression of CHL1 on EVs, or both. In any case, purified and detected EVs were considered to contain EVs secreted from *EML4-ALK* LC cells. CHL1 BiEV can therefore be used as an indicator of cEV, but confirmation is needed. The candidate partner molecules for CHL1 BiEV in Table 1 have been identified by MS, and another confirmation technique, such as immunoprecipitation or western blot experiments, is required to identify whether they are true partner molecules. Among the molecules listed in Table 1, some candidates BiEV partner molecules did not form BiEVs specifically in serum EVs from LC. In contrast, CHL1-THBS1 and CHL1-SLC4A1 BiEV were clearly expressed in serum EV even in LC model mice, particularly in individuals with small tumors (Figure 3C and 3D).

In this study, we preferentially analyzed SLC4A1, a membrane protein, and compared it with THBS1. However, because several studies have reported THBS1 is expressed in exosomes and is also involved in cancer cell migration, invasion, metastasis, and epithelial-mesenchymal transition [35–37], further study is warranted.

ELISA quantification revealed that the CHL1-SLC4A1 BiEV in mouse serum EVs are applicable biomarkers for screening small tumors in *EML4-ALK* transgenic mice (Figure 4) compared with the serum CHL1 level, which is a single biomarker for LC cells, as reported previously [25]. Tumor size in *EML4-ALK* transgenic mouse was partially correlated with the value of CHL1-SLC4A1 BiEV (Figure 4C) supporting the evidence for direct secretion of such EVs in cancer cells. These advantages allowed us to discuss not only the sensitivity of BiEV analysis for typical EV screening but also its applicability to cancer screening in human cancer patients, who usually present with small tumor masses. The SLC4A1 molecule is a multi-pass transmembrane chloride bicarbonate exchange transporter [38]. In addition, it has also been found to be markedly upregulated in several cancer cells [39]. However, the average expression level of SLC4A1 in the whole-serum EVs was lower in LC model mice that in WT mice (Figure 4D). The amount of fluorescein-labeled SLC4A1, which is indicated as CHL1-SLC4A1 BiEV, should be quantified using this BiEV analysis.

The application of BiEV analysis in human serum EV screening is an important issue. According to our study, CHL1-SLC4A1 BiEV detected in mouse LC did not appear to be significant in human LC patients (Figure S5A and S5B). It is therefore necessary to identify the appropriate BiEV according to animal species as well as cancer type. In human screening, it is assumed that the appropriate BiEV differs by tissue type within a cancer type. In the case of LC, small cell carcinoma, non-small cell carcinoma, adenocarcinoma, squamous cell carcinoma, and among others are listed as categories. In consideration of experimental efficiency and the value of human serum sources, our experiments were performed by integrating serum, but it will be important to study these factors in the future.

In this study, we have shown that measurement of CHL1–caspase 14 BiEV may be useful for screening cEV in LC patients. Caspase 14 is a caspase family protease [30] and is not a membrane protein but expressed in desmosomes [40] and urinary exosomes [41], demonstrating that caspase 14 could be secreted and attached with membrane. Interestingly, CHL1–caspase 14 BiEV has also been detected in EVs secreted from LK2 human LC culture cell line (Table 2), suggesting that EVs expressing CHL1–caspase 14 BiEV were directly secreted from LC cells in patients. LK2 cell secretion experiments with serum-free medium indicated that CHL1–caspase 14 BiEV is not be formed by binding of free caspase 14 in patient serum but display caspase 14 on the EV surface during the intracellular process of EV formation in LC cells. Similar to SLC4A1 in mice, the high expression levels of caspase 14 itself has been observed in cancer cells [42–44], but expression of caspase 14 in healthy EVs and cancer EVs exhibited no significant differences as a whole (Figure 5E). This also demonstrates that fluorescein-labeled caspase 14 should be quantified in BiEV analysis.

BiEV strategies have some problems to overcome. Although the cause is unknown, in some cases, neither CHL1-SLC4A1 nor CHL1–caspase 14 BiEV were increased in serum EVs even in *EML4-ALK* transgenic mouse and LC patients (Figure 4 and 5). Because cancer screening does not perform multiple tests of biomarkers, it is necessary to search for appropriate BiEVs to produce high AUC scores in ROC analyses (Figure 4B and 5E). In addition, serum EV purification and EMARS reactions of the BiEV strategy were slightly complicated and time consuming. As a solution, a novel ELISA technique is required. Because serum EVs were approximately 20 to 100 nm in size according to cryo-electron microscopy (Figure 2D), a more reasonable fluorescein-labeling distance (within 20 nm [8]) would allow a BiEV to be measured directly by the ExoScreen method, which easily detects and quantifies proximal molecules in an EV particle [45]. To use BiEV information in the clinical field, development of such a system will be necessary.

Taken together, the BiEV strategy proved useful for the screening in EV classification. Considerable advantages should become apparent if these BiEVs are used as a novel biomarker (cancer EV antigen) for cancer screening or at least molecular diagnosis.

## Abbreviations

EV: extracellular vesicle
cEV: cancer cell-secreted EV
tEV: 
cEV: target EV
BiEV: bimolecular surface antigen expressed in EV
EMARS: enzyme-mediated activation of radical source
BiCAT: Bimolecular Cancer Target
CHL1: close homolog of L1
LC: lung cancer
MS: mass spectrometry

## Acknowledgments

We thank Professor Hiroyuki Mano and Dr. Manabu Soda for providing *EML4-ALK* transgenic mice; Medical Genome Center Biobank, National Center for Geriatrics and Gerontology and BioBank Japan Project (supported by AMED) for providing human serum samples; Laboratory of Cytometry and Proteome Research, Tokyo Medical and Dental University, and Ms. Amimoto of the Natural Science Center for Basic Research and Development (N-BARD), Hiroshima University for MS technical assistance; Professor Tsumoru Shintake for helping with electron microscopic analysis; Kochi University’s experimental training equipment facility and the Saitama Medical University Biomedical Research Center for providing general technical assistance. This work was supported by Grants-in-aid for Scientific Research in Japan (No. JP15K07941, and No. JP18K06663 to N. K.; JP17H04409 to T. S.), Mizutani foundation research grants (to N. K.), and Japan science and technology grants (to N. K.). We also thank enago group for editing a draft of this manuscript.

## Potential Conflict of Interest

The authors declare that they have no conflicts of interest with the contents of this article.

---

**Figure S1.**
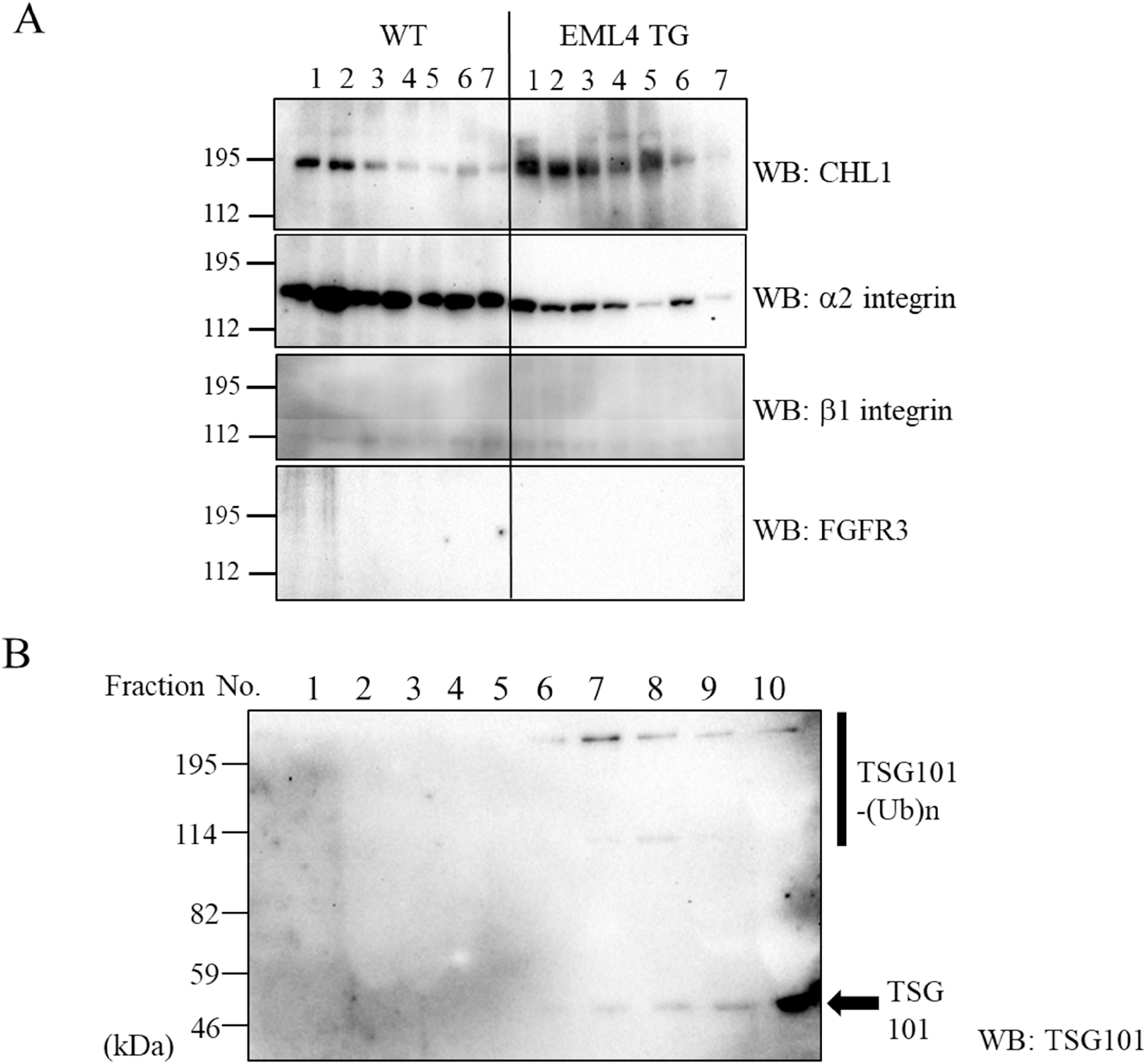
Characterization of crude and purified serum EVs. (*A*) Protein expression of precipitated EVs. The serum collected from seven mice of wild-type (WT) and *EML4-ALK* transgenic mouse (TL group) was subjected to Western blot analysis with anti-CHL1, anti-α2 integrin, anti-β1 integrin, and anti-FGFR3 antibodies, which were cancer cell membrane BiCAT molecules previously reported. (*B*) Western blot analysis for TSG101 antigen detection in Sephacryl S-500 fractions. The fractions were concentrated with *Nanosep*® centrifugal unit, and then subjected to Western blot analysis with anti-TSG101 antibody. TSG101-(Ub)n indicates ubiquitinated TSG101.

**Figure S2.**
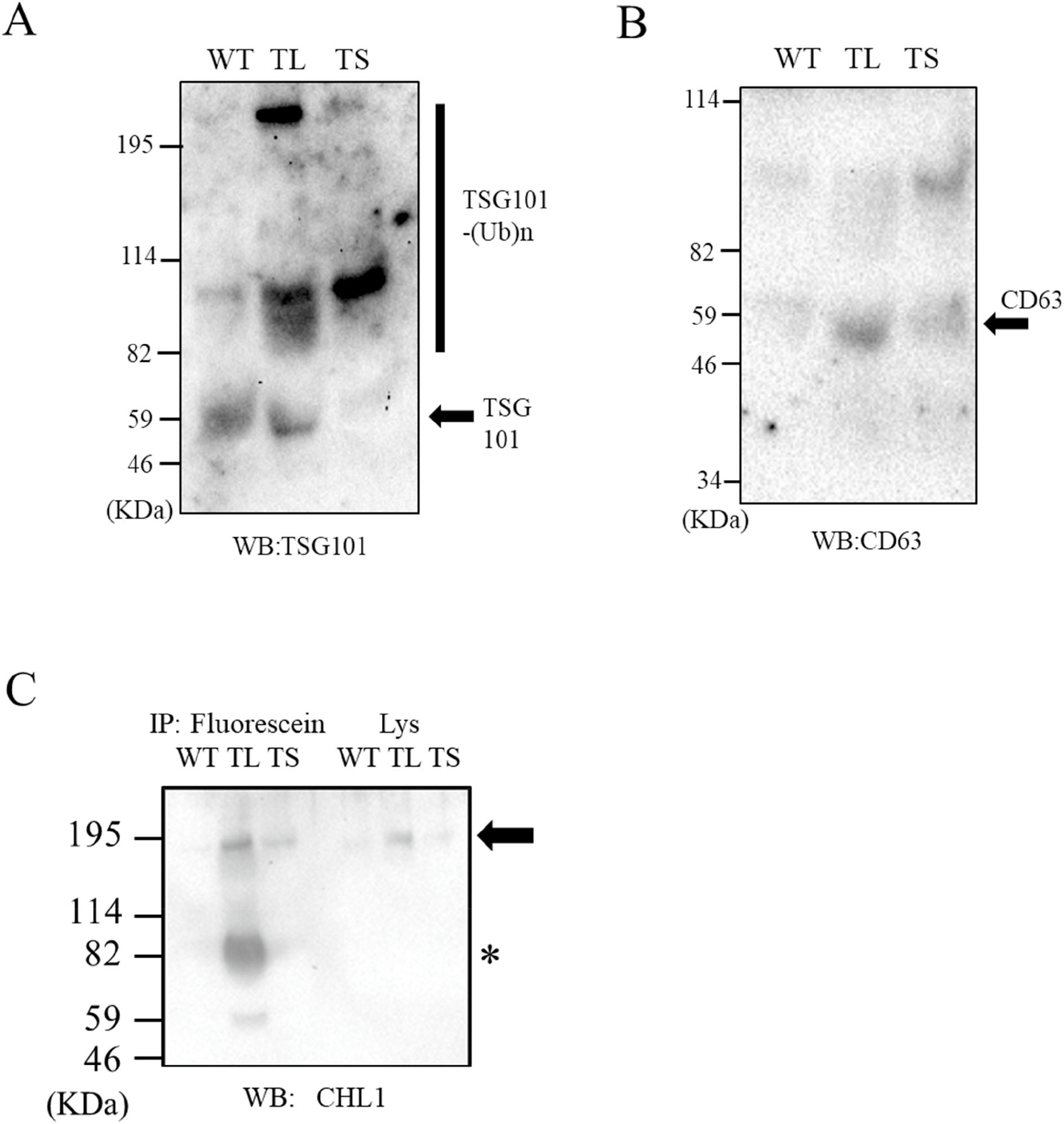
Characterization of serum EVs from WT, TL, and TS mice. (*A, B*) Western blot analysis of TSG101 (*A*) and CD63 (*B*) in serum EVs from WT, TL, and TS using anti-TSG101 and CD63 antibody. Arrows indicate the detected band of monomer TSG101 and CD63. The black bar indicates wide range of molecular weight due to an ubiquitination in TSG101 (TSG101-(Ub)n). (*C*) Western blot analysis of EMARS products with anti-CHL1 antibody. The EMARS products were concentrated and purified by immunoprecipitation with the anti-fluorescein antibody Sepharose. The resulting samples were subjected to SDS-PAGE analysis with fluorescence detection. “IP” indicates the immunoprecipitated samples, and “Lys” indicates the lysate samples before immunoprecipitation. Arrow indicates the detected band of CHL1. Asterisk indicates unknown bands (predicted as non-specific or partial fragments).

**Figure S3.**
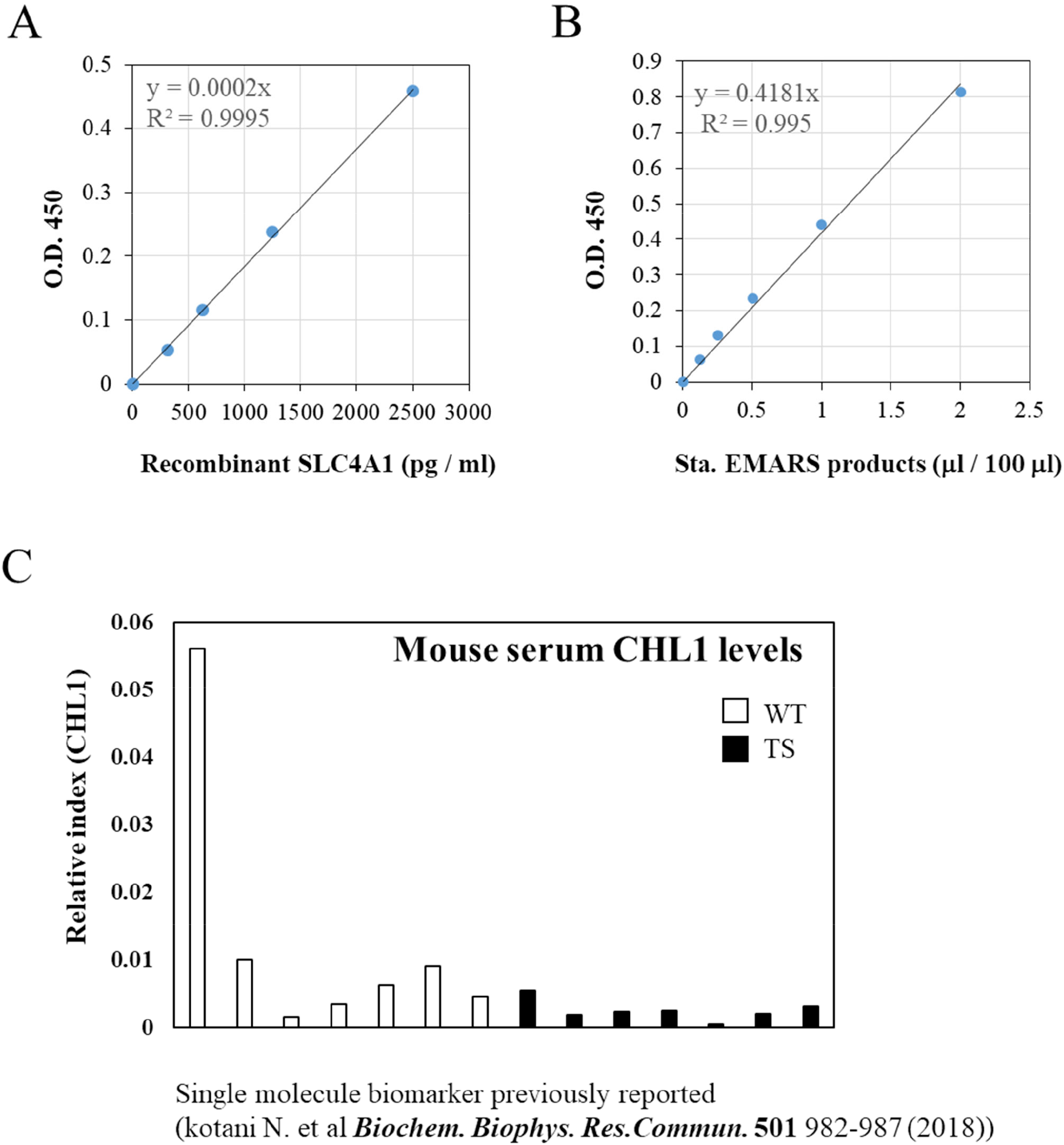
Characterization of ELISA system for SLC4A1. (*A, B*) Calibration curve of sandwich ELISA for the detection of both SLC4A1 partial proteins and fluorescein-labeled SLC4A1. The detection of several concentrations of recombinant SLC4A1 partial protein using HRP-labeled anti-SLC4A1 antibody which is prepared using Zenon system is summarized in (*A*). The detection of several concentrations of self-made standard materials containing fluorescein-labeled SLC4A1 using HRP-labeled anti-fluorescein antibody is summarized in (*B*). (*C*) Comparison of serum CHL1 levels between wild-type (WT) and small tumor-bearing *EML4-ALK* transgenic (TS) mice by using previously established ELISA system for CHL1 measurement. There were no significant differences between them.

**Figure S4.**
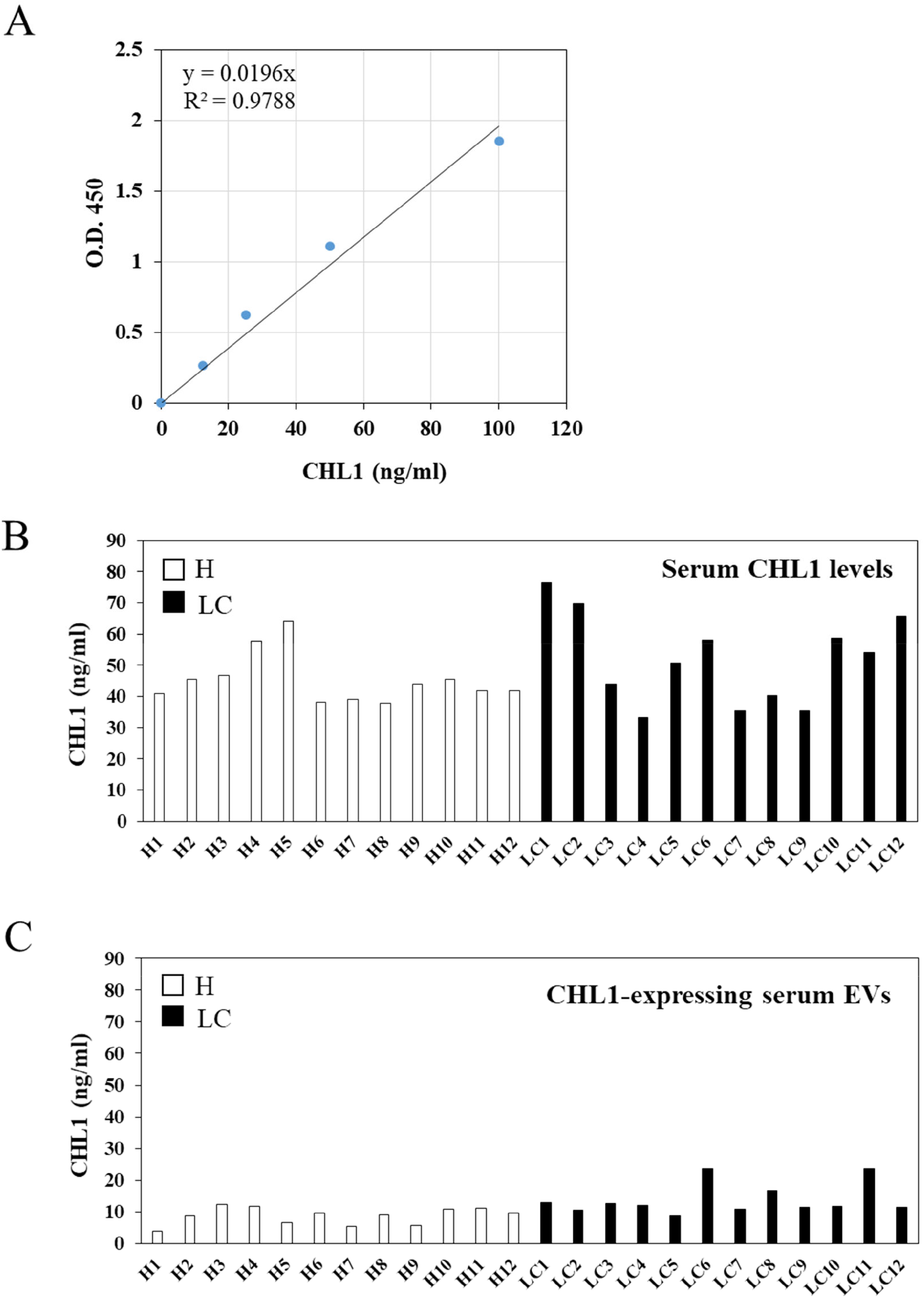
Characterization of ELISA system for human CHL1. (*A*) Calibration curve of sandwich ELISA for the detection of both human CHL1. The detection of several concentrations of recombinant human CHL1 partial protein using HRP-labeled anti-CHL1 antibody. (*B*) Comparison of serum CHL1 levels between H (open bar) and LC (closed bar) by using ELISA system for human CHL1 measurement. There were no significant differences between them. (*C*) Comparison of CHL1-expressing EVs between H (open bar) and LC (closed bar). The serum EVs were purified by using ExoQuick Solution followed by ELISA measurement of CHL1 levels. There were also no significant differences between them.

**Figure S5.**
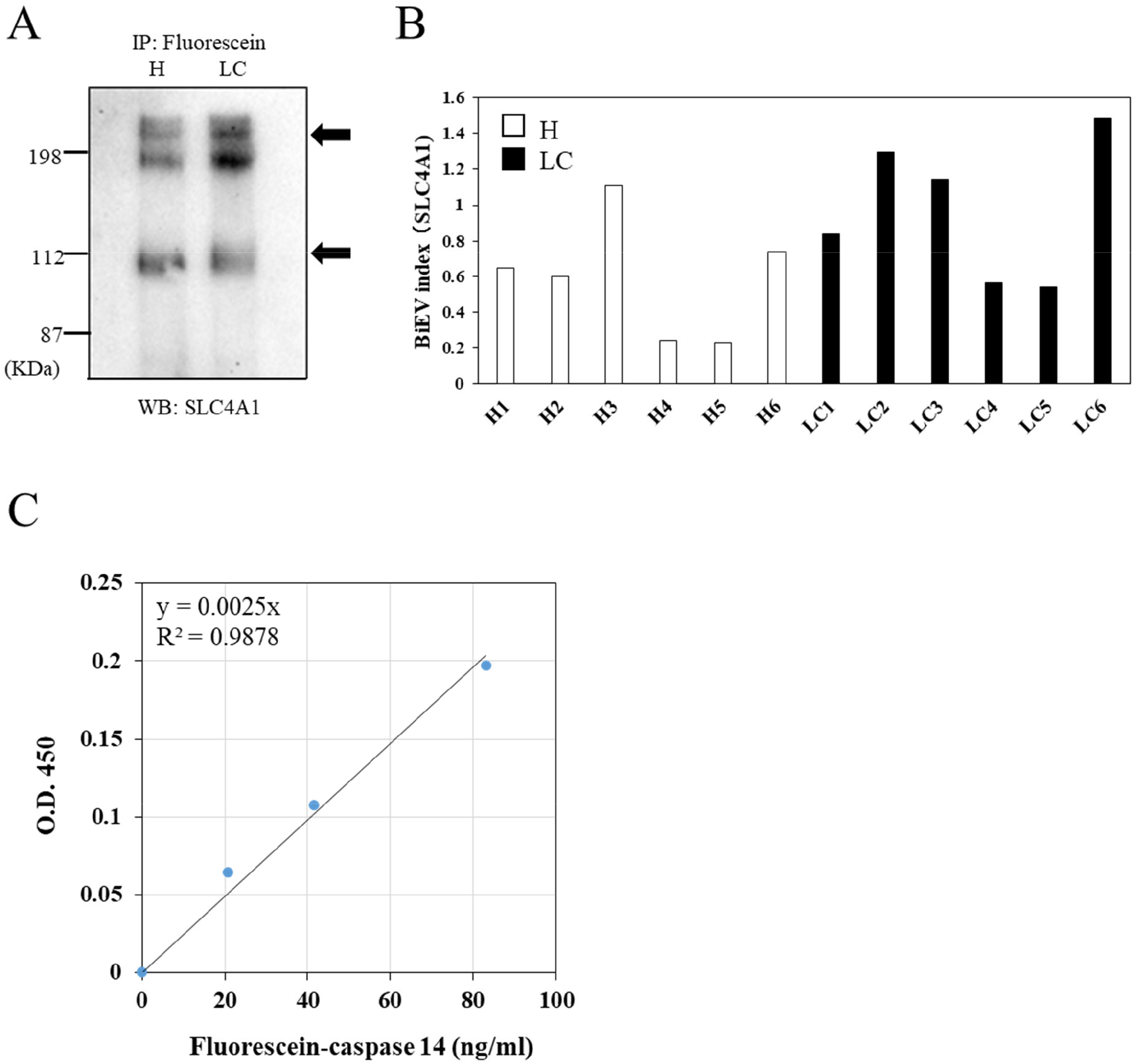
Comparison of CHL1-SLC4A1 BiEV in human serum EVs and characterization of ELISA system for fluorescein labeled-caspase 14. (*A*) The healthy person (H) and lung cancer patients (LC) EAMRS samples were respectively applied to immunoprecipitation (anti-fluorescence antibody Sepharose) and western blot analysis with anti-SLC4A1 antibody. Arrows indicate the detected band of human SLC4A1 proteins (including predicted dimer). (*B*) Measurement of fluorescein-labeled human SLC4A1 using sandwich ELISA system used in mouse fluorescein-labeled SLC4A1. The serum EVs from 6 H (open bar) and 6 LC (closed bar) were applied to EMARS reaction followed by ELISA measurement, respectively. The EMARS products containing fluorescein-labeled SLC4A1 were added to anti-SLC4A1 antibody-coated ELISA plates. “BiEV index (SLC4A1)” was calculated based on the value of fluorescein-labeled SLC4A1 in standard samples as described in “*Materials and methods*”. (*C*) Calibration curve of sandwich ELISA for the detection of fluorescein-labeled caspase 14. The detection of several concentrations of fluorescein-labeled caspase 14 standard was performed using HRP-labeled anti-fluorescein antibody.

## Notes

### Competing Interest Statement

The authors have declared no competing interest.

### Summary of Updates

Figure revised

