## Supplementary material for "Bimolecule detection for Extracellular Vesicle Screening": Table S1

### Table S1 Raw data of MS analysis for mouse BiEVs (WT)

#### Protein Report

BRUKER

##### Project Info

| | | | |
|---|---|---|---|
| **Name:** | NAWA | **Date:** | October 6, 2015 |

##### Sample Info & Protocols

| | | | |
|---|---|---|---|
| | | **Date:** | March 1, 2018 |
| **Name:** | 2018-3 | | |

##### Search Result Info

| Search Result | Location | Search Engine | Database | Ident. Compounds |
|---|---|---|---|---|
| maXis_mouse-2016_Mascot_2018-03-02 16:30:42 | /NAWA/2018-3/0301-kotani-wt_Tray01-E1_01_6598.mgf | Mascot, 2.6.0 | SwissProt, SwissProt_2017_07.fasta | 11/1543 |

| | | | |
|---|---|---|---|
| **Protein 1:** | Ig mu chain C region OS=Mus musculus GN=Ighm PE=1 SV=2 | | |
| **Accession:** | IGHM_MOUSE | **Score:** | 411.52 |
| **Database:** | SwissProt | **MW [kDa]:** | 49.90 |
| **Seq. Coverage [%]:** | 14.30 % | **pI:** | 6.56 |
| | | **No. of Peptides:** | 5 |
| **Modification(s):** | Carbamidomethyl | | |

| 10 | 20 | 30 | 40 | 50 | 60 | 70 | 80 | 90 | 100 | 110 | 120 |
|---|---|---|---|---|---|---|---|---|---|---|---|
| SQSFPNVFPL | VSCESPLSDK | NLVAMGCLAR | DFLPSTISFT | WNYQNNTEVI | QGIRTFPTLR | TGGKYLATSQ | VLLSPKSILE | GSDEYLVCKI | HYGGKNRDLH | VPIPAVAEMN | PNVNVFVPPR |
| 130 | 140 | 150 | 160 | 170 | 180 | 190 | 200 | 210 | 220 | 230 | 240 |
| DGFSGPAPRK | SKLICEATNF | TPKPITVSWL | KDGKLVESGF | TTDPVTIENK | GSTPQTYKVI | STLTISEIDW | LNLNVYTCRV | DHRGLTFLKN | VSSTCAASPS | TDILTFTIPP | SFADIFLSKS |
| 250 | 260 | 270 | 280 | 290 | 300 | 310 | 320 | 330 | 340 | 350 | 360 |
| ANLTCLVSNL | ATYETLNISW | ASQSGEPLET | KIKIMESHPN | GTFSAKGVAS | VCVEDWNNRK | EFVCTVTHRD | LPSPQKKFIS | KPNEVHKHPP | AVYLLPPARE | QLNLRESATV | TCLVKGFSPA |
| 370 | 380 | 390 | 400 | 410 | 420 | 430 | 440 | 450 | 460 | | |
| DISVQWLQRG | QLLPQEKYVT | SAPMPEPGAP | GFYFTHSILT | VTEEEWNSGE | TYTCVVGHEA | LPHLVTERTV | DKSTGKPTLY | NVSLIMSDTG | GTCY | | |

| Cmpd. | m/z meas. | Δ m/z [ppm] | z | Rt [min] | Score | P | Range | Sequence | Modification |
|---|---|---|---|---|---|---|---|---|---|
| 715 | 552.7840 | -4.16 | 2 | 38.26 | 35.36 | 0 | 21-30 | K.NLVAMGCLAR.D | Carbamidomethyl: 7 |
| 756 | 660.3794 | -4.12 | 2 | 39.55 | 94.75 | 0 | 65-76 | K.YLATSQVLLSPK.S | |
| 822 | 756.8633 | -3.81 | 2 | 41.64 | 76.81 | 0 | 77-89 | K.SILEGSDEYLVCK.I | Carbamidomethyl: 12 |

| Cmpd. | m/z meas. | Δ m/z [ppm] | z | Rt [min] | Score | P | Range | Sequence | Modification |
|---|---|---|---|---|---|---|---|---|---|
| 783 | 875.4448 | -4.71 | 2 | 40.45 | 121.38 | 0 | 155-170 | K.LVESGFTTDPVTIENK.G | |
| 1286 | 802.4054 | -11.93 | 2 | 56.79 | 83.22 | 0 | 356-369 | K.GFSPADISVQWLQR.G | |

**Protein 2:** Keratin, type II cytoskeletal 73 OS=Mus musculus GN=Krt73 PE=1 SV=1

| | | | |
|---|---|---|---|
| **Accession:** | K2C73_MOUSE | **Score:** | 88.96 |
| **Database:** | SwissProt | **MW [kDa]:** | 58.90 |
| **Seq. Coverage [%]:** | 4.30 % | **pI:** | 8.36 |
| | | **No. of Peptides:** | 2 |

| 10 | 20 | 30 | 40 | 50 | 60 | 70 | 80 | 90 | 100 | 110 | 120 |
|---|---|---|---|---|---|---|---|---|---|---|---|
| MNRQFTCKPG | VANRGFSGCS | AVLSGGSSSS | YRAAGKGLSG | GFSSRSLYSL | RSPRSISFNV | ASSSGRTGGY | GFGRNRASGF | AGSMFGSGAL | GPSNPSLCLP | GGIHQVTVNK | SLLAPLNVEL |
| 130 | 140 | 150 | 160 | 170 | 180 | 190 | 200 | 210 | 220 | 230 | 240 |
| DPEIQKVRAQ | EREQIKALNN | KFASFIDKVR | FLEQQNQVLQ | TKWELLQQLD | LSNCRRNLEP | VYEAHISSLQ | KQLDSLSGDR | VRLDSELRGM | RDAVEDCKKR | YEEEINKRTT | AENEFVVLKK |
| 250 | 260 | 270 | 280 | 290 | 300 | 310 | 320 | 330 | 340 | 350 | 360 |
| DVDAAYMSKV | ELQAKVDALD | GEIKFLKCLY | EGEITQMQSH | ISDTSVVLSM | DNNRNLDLDS | IIAEVRAQYE | DIALKSKAEA | EMVYQTKFQE | LQLAAGRHGD | DLKHTRNEIS | ELTRLIQRLR |
| 370 | 380 | 390 | 400 | 410 | 420 | 430 | 440 | 450 | 460 | 470 | 480 |
| SEIESVKKQC | SNLETAIADA | EQRGDCALKD | ARAKLDELER | ALHQAKEELA | RMLREHQELM | SMKLALDIEI | ATYRKLLEGE | ECRMSGEHTS | AVSISVISSS | APGTVGAGTS | FGFSSAGTYG |
| 490 | 500 | 510 | 520 | 530 | 540 | | | | | | |
| YRQSSVAGGY | GILSGGCVTG | SGNCSPRGDT | KNRLGSASEF | KEVSGKTLAL | GSPSKKTMR | | | | | | |

| Cmpd. | m/z meas. | Δ m/z [ppm] | z | Rt [min] | Score | P | Range | Sequence | Modification |
|---|---|---|---|---|---|---|---|---|---|
| 803 | 679.3494 | -29.97 | 2 | 41.14 | 34.09 | 0 | 295-306 | R.NLDLDSIIAEVR.A | |
| 1129 | 639.3527 | -9.24 | 2 | 51.40 | 54.87 | 0 | 424-434 | K.LALDIEIATYR.K | |

#### Protein Report

**Protein 3:** Ig kappa chain C region OS=Mus musculus PE=1 SV=1

| | | | |
|---|---|---|---|
| **Accession:** | IGKC_MOUSE | **Score:** | 83.86 |
| **Database:** | SwissProt | **MW [kDa]:** | 11.80 |
| **Seq. Coverage [%]:** | 13.20 % | **pI:** | 5.23 |
| | | **No. of Peptides:** | 1 |

| 10 | 20 | 30 | 40 | 50 | 60 | 70 | 80 | 90 | 100 | 110 |
|---|---|---|---|---|---|---|---|---|---|---|
| ADAAPTVSIF | PPSSEQLTSG | GASVVCFLNN | FYPKDINVKW | KIDGSERQNG | VLNSWTDQDS | KDSTYSMSST | LTLTKDEYER | HNSYTCEATH | KTSTSPIVKS | FNRNEC |

| Cmpd. | m/z meas. | Δ m/z [ppm] | z | Rt [min] | Score | P | Range | Sequence | Modification |
|---|---|---|---|---|---|---|---|---|---|
| 791 | 767.8650 | -5.02 | 2 | 40.73 | 83.86 | 0 | 62-75 | K.DSTYSMSSTLTLTK.D | |

**Protein 4:** Keratin, type II cytoskeletal 5 OS=Mus musculus GN=Krt5 PE=1 SV=1

| | | | |
|---|---|---|---|
| **Accession:** | K2C5_MOUSE | **Score:** | 75.50 |
| **Database:** | SwissProt | **MW [kDa]:** | 61.70 |
| **Seq. Coverage [%]:** | 2.10 % | **pI:** | 7.59 |
| | | **No. of Peptides:** | 1 |

| 10 | 20 | 30 | 40 | 50 | 60 | 70 | 80 | 90 | 100 | 110 | 120 |
|---|---|---|---|---|---|---|---|---|---|---|---|
| MSRQSSVSFR | SGGSRSFSAA | SAITPSVSRT | SFSSVSRSGG | GGGGRISLGG | ACGAGGYGSR | SLYNVGGSKR | ISYSSGGGSF | RNQFGAGGFG | FGGGAGSGFG | FGGGAGSGFG | FGGGAGFGGG |
| 130 | 140 | 150 | 160 | 170 | 180 | 190 | 200 | 210 | 220 | 230 | 240 |
| YGGAGFPVCP | PGGIQEVTVN | QNLLTPLNLQ | IDPTIQRVRT | EEREQIKTLN | NKFASFIDKV | RFLEQQNKVL | DTKWALLQEQ | GTKTIKQNLD | PLFEQYINNL | RRQLDGVLGE | RGRLDSELRN |
| 250 | 260 | 270 | 280 | 290 | 300 | 310 | 320 | 330 | 340 | 350 | 360 |
| MQDLVEDYKN | KYEDEINKRT | TAENEFVMLK | KDVDAAYMNK | VELEARVDAL | MDEINFMKMF | FDAELSQMQT | HVSDTSVVLS | MDNNRSLDLD | SIIAEVKAQY | EDIANRSRTE | AESWYQTKYE |
| 370 | 380 | 390 | 400 | 410 | 420 | 430 | 440 | 450 | 460 | 470 | 480 |
| ELQQTAGRHG | DDLRNTKHEI | SEMNRMIQRL | RSEIDNVKKQ | CANLQNAIAE | AEQRGELALK | DARNKLTELE | EALQKAKQDM | ARLLREYQEL | MNTKLALDVE | IATYRKLLEG | EECRLSGEGV |
| 490 | 500 | 510 | 520 | 530 | 540 | 550 | 560 | 570 | 580 | 590 | |
| GPVNISVVTN | SVSSGYGGGS | SIGVGSGFGG | GLGSGFAGGL | GPRFTRGGGG | LGLGSGLSVG | GSGFSAGSSQ | GGMSFGSGGG | SGSSVKFVST | TSSSRRSFKS | | |

### Protein Report

| Cmpd. | m/z meas. | Δ m/z [ppm] | z | Rt [min] | Score | P | Range | Sequence | Modification |
|---|---|---|---|---|---|---|---|---|---|
| 1347 | 651.8553 | -9.13 | 2 | 58.95 | 75.50 | 0 | 326-337 | R.SLDLDSIIAEVK.A | |

**Protein 5:** Keratin, type II cytoskeletal 79 OS=Mus musculus GN=Krt79 PE=1 SV=2

**Accession:** K2C79_MOUSE

**Database:** SwissProt

**Seq. Coverage [%]:** 2.30 %

**Score:** 69.30

**MW [kDa]:** 57.50

**pI:** 7.55

**No. of Peptides:** 1

| 10 | 20 | 30 | 40 | 50 | 60 | 70 | 80 | 90 | 100 | 110 | 120 |
|---|---|---|---|---|---|---|---|---|---|---|---|
| MRSSLSRQTF | STKGGFSSNS | ASGGGGSRMR | TSYSSVTMSR | GSGGGGGVRS | GSSSGGFGSR | SLYNLGGKNI | SVSMACGASS | GRALGGFGSG | AYVGLGASRQ | TFGPVCPPGG | IQEVTVNQSL |
| 130 | 140 | 150 | 160 | 170 | 180 | 190 | 200 | 210 | 220 | 230 | 240 |
| LTPLNVEIDP | EIQRVRTQER | EQIKTLNNKF | ASFIDKVRFL | EQQNKVLETK | WALLQEQSQN | TGVARSLEPF | FENYLSTLRR | QLDTKQSERG | RLDMELRNVQ | DNLEDFKNKY | EDEINKRTAL |
| 250 | 260 | 270 | 280 | 290 | 300 | 310 | 320 | 330 | 340 | 350 | 360 |
| ENEFVLLKKD | VDAAYMGRMD | LHGKVDSLTQ | EIDFLQQLFE | MELSQVQTNV | SDTNVILSMD | NNRNLDLDSI | IAEVKAQYEL | IAQKSRAEAE | SWYQTKYEEL | QVTAGKHGDS | LRDTKNEIAE |
| 370 | 380 | 390 | 400 | 410 | 420 | 430 | 440 | 450 | 460 | 470 | 480 |
| LTRTTQRLQG | EVDAAKKQCQ | QLQTAIAEAE | QNGEMALKDA | KKKLGDLDTA | LHQAKEDLAR | MLREYQDLVS | VKLALDMEIA | TYRKLLESEE | SRMSGDCPSA | ISISVTGNST | SVCAGGTAGF |
| 490 | 500 | 510 | 520 | 530 | 540 | | | | | | |
| GNGLSLGGAG | GASKGGFGSS | VSYGAAKGGQ | VSGGTSILRK | TTTVKTSSRR | Y | | | | | | |

| Cmpd. | m/z meas. | Δ m/z [ppm] | z | Rt [min] | Score | P | Range | Sequence | Modification |
|---|---|---|---|---|---|---|---|---|---|
| 1344 | 665.3580 | -13.03 | 2 | 58.84 | 69.30 | 0 | 304-315 | R.NLDLDSIIAEVK.A | |

**Protein 6:** Keratin, type II cytoskeletal 2 epidermal OS=Mus musculus GN=Krt2 PE=1 SV=1

**Accession:** K22E_MOUSE

**Database:** SwissProt

**Seq. Coverage [%]:** 1.40 %

**Score:** 40.12

**MW [kDa]:** 70.90

**pI:** 8.26

**No. of Peptides:** 1

| 10 | 20 | 30 | 40 | 50 | 60 | 70 | 80 | 90 | 100 | 110 | 120 |
|---|---|---|---|---|---|---|---|---|---|---|---|
| MSCQISCRSR | RGGGGGGGGG | FRGFSSGSAV | VSGGSRRSNT | SFSCISRHGG | GRGGSGGGGF | GSQSLVGLGG | YKSISSSVAG | NSGGYGGSSF | GGSSGFGGGR | GFGGGQGFGG | SGGFGGGSGF |
| 130 | 140 | 150 | 160 | 170 | 180 | 190 | 200 | 210 | 220 | 230 | 240 |
| GGGQGFGGGS | RFGGGSGFGG | GGFGGGSFGG | GRFGGGPGGF | GGPGGFPGGG | IHEVSVNQSL | LQPLDVKVDP | EIQNVKSQER | EQIKTLNNKF | ASFIDKVRFL | EQQNQVLRTK | WELLQQLDVG |
| 250 | 260 | 270 | 280 | 290 | 300 | 310 | 320 | 330 | 340 | 350 | 360 |
| SRTTNLDPIF | QAYIGMLKKQ | VDRLSAERTS | QESELNNMQD | LVEDFKKKYE | DEINKRTSAE | NDFVTIKKDV | DSCYMDKTEL | QARLDILAQE | VNFLRTLYDA | ELSQLQQDVT | DTNVILSMDN |
| 370 | 380 | 390 | 400 | 410 | 420 | 430 | 440 | 450 | 460 | 470 | 480 |
| NRNLDLDSII | AEVQNQYEMI | AHKSKAESEE | LYHSKYEELQ | VTAVKHGDSL | KEIKMEISEL | NRTIQRLQGE | ISHVKKQCKG | VQDSIADAEQ | RGEHAIKDAR | GKLTDLEEAL | QQCREDLARL |
| 490 | 500 | 510 | 520 | 530 | 540 | 550 | 560 | 570 | 580 | 590 | 600 |
| LRDYQELMNT | KLSLDVEIAT | YRKLLEGEEC | RMSGDFSDNV | SVSITSSTIS | SSVASKTGFG | SGGQSSGGRG | SYGGRGGGGG | GGSTYGSGGR | SSGSRGSGSG | SGGGGYSSGG | GSRGGSGGGY |
| 610 | 620 | 630 | 640 | 650 | 660 | 670 | 680 | 690 | 700 | 710 | |
| GSGGGSRGGS | GGGYGSGGGS | GSGGGYSSGG | GSRGGSGGGG | VSSGGGSRGG | SSSGGGSRGG | SSSGGGGYSS | GGGSRGGSSS | GGAGSSSEKG | GSGSGEGCGS | GVTFSFR | |

| Cmpd. | m/z meas. | Δ m/z [ppm] | z | Rt [min] | Score | P | Range | Sequence | Modification |
|---|---|---|---|---|---|---|---|---|---|
| 539 | 590.3008 | -26.47 | 2 | 32.95 | 40.12 | 0 | 396-405 | K.YEELQVTAVK.H | |

### Table S1. Raw data of MS analysis for mouse BiEVs (TL)

#### Protein Report

BRUKER

##### Project Info

| | | | |
|---|---|---|---|
| **Name:** | NAWA | **Date:** | October 6, 2015 |

##### Sample Info & Protocols

| | | | |
|---|---|---|---|
| | | **Date:** | March 1, 2018 |
| **Name:** | 2018-3 | | |

##### Search Result Info

| Search Result | Location | Search Engine | Database | Ident. Compounds |
|---|---|---|---|---|
| maXis_mouse-2016_Mascot_2018-03-02 16:16:47 | /NAWA/2018-3/0302-kotani-D_Tray01-E3_01_6616.mgf | Mascot, 2.6.0 | SwissProt, SwissProt_2017_07.fasta | 94/1489 (FDR:4.72%) |

| | | | |
|---|---|---|---|
| **Protein 1:** | Ig mu chain C region OS=Mus musculus GN=Ighm PE=1 SV=2 | | |
| **Accession:** | IGHM_MOUSE | **Score:** | 1152.31 |
| **Database:** | SwissProt | **MW [kDa]:** | 49.90 |
| **Seq. Coverage [%]:** | 44.70 % | **pI:** | 6.56 |
| | | **No. of Peptides:** | 18 |
| **Modification(s):** | Carbamidomethyl, Oxidation | | |

| 10 | 20 | 30 | 40 | 50 | 60 | 70 | 80 | 90 | 100 | 110 | 120 |
|---|---|---|---|---|---|---|---|---|---|---|---|
| SQSFPNVFPL | VSCESPLSDK | NLVAMGCLAR | DFLPSTISFT | WNYQNNTEVI | QGIRTFPTLR | TGGKYLATSQ | VLLSPKSILE | GSDEYLVCKI | HYGGKNRDLH | VPIPAVAEMN | PNVNVFVPPR |
| 130 | 140 | 150 | 160 | 170 | 180 | 190 | 200 | 210 | 220 | 230 | 240 |
| DGFSGPAPRK | SKLICEATNF | TPKPITVSWL | KDGKLVESGF | TTDPVTIENK | GSTPQTYKVI | STLTISEIDW | LNLNVYTCRV | DHRGLTFLKN | VSSTCAASPS | TDILTFTIPP | SFADIFLSKS |
| 250 | 260 | 270 | 280 | 290 | 300 | 310 | 320 | 330 | 340 | 350 | 360 |
| ANLTCLVSNL | ATYETLNISW | ASQSGEPLET | KIKIMESHPN | GTFSAKGVAS | VCVEDWNNRK | EFVCTVTHRD | LPSPQKKFIS | KPNEVHKHPP | AVYLLPPARE | QLNLRESATV | TCLVKGFSPA |
| 370 | 380 | 390 | 400 | 410 | 420 | 430 | 440 | 450 | 460 | | |
| DISVQWLQRG | QLLPQEKYVT | SAPMPEPGAP | GFYFTHSILT | VTEEEWNSGE | TYTCVVGHEA | LPHLVTERTV | DKSTGKPTLY | NVSLIMSDTG | GTCY | | |

| Cmpd. | m/z meas. | Δ m/z [ppm] | z | Rt [min] | Score | P | Range | Sequence | Modification |
|---|---|---|---|---|---|---|---|---|---|
| 690 | 552.7835 | -4.98 | 2 | 38.03 | 51.12 | 0 | 21-30 | K.NLVAMGCLAR.D | Carbamidomethyl: 7 |
| 359 | 560.7804 | -5.97 | 2 | 27.96 | 37.10 | 0 | 21-30 | K.NLVAMGCLAR.D | Carbamidomethyl: 7; Oxidation: 5 |

| Cmpd. | m/z meas. | Δ m/z [ppm] | z | Rt [min] | Score | P | Range | Sequence | Modification |
|---|---|---|---|---|---|---|---|---|---|
| 483 | 367.7126 | -2.10 | 2 | 31.77 | 28.62 | 0 | 55-60 | R.TFPTLR.T | |
| 729 | 660.3771 | -7.61 | 2 | 39.24 | 76.40 | 0 | 65-76 | K.YLATSQVLLSPK.S | |
| 797 | 756.8616 | -5.98 | 2 | 41.33 | 91.47 | 0 | 77-89 | K.SILEGSDEYLVCK.I | Carbamidomethyl: 12 |
| 1304 | 842.4405 | -10.40 | 3 | 58.28 | 95.43 | 0 | 98-120 | R.DLHVPIPAVAEMNPNVNVFVPPR.D | |
| 179 | 452.2164 | -7.17 | 2 | 22.08 | 52.65 | 0 | 121-129 | R.DGFSGPAPR.K | |
| 1142 | 740.0627 | -9.28 | 3 | 52.79 | 58.62 | 0 | 133-151 | K.LICEATNFTPKPITVSWLK.D | Carbamidomethyl: 3 |
| 765 | 875.4447 | -4.83 | 2 | 40.33 | 146.64 | 0 | 155-170 | K.LVESGFTTDPVTIENK.G | |
| 679 | 339.7123 | -1.66 | 2 | 37.69 | 18.91 | 0 | 204-209 | R.GLTFLK.N | |
| 643 | 753.3401 | -4.77 | 2 | 36.66 | 89.13 | 0 | 287-299 | K.GVASVCVEDWNNR.K | Carbamidomethyl: 6 |
| 119 | 426.2193 | -2.50 | 3 | 19.42 | 32.60 | 1 | 300-309 | R.KEFVCTVTHR.D | Carbamidomethyl: 5 |
| 202 | 574.7762 | -5.65 | 2 | 22.94 | 50.17 | 0 | 301-309 | K.EFVCTVTHR.D | Carbamidomethyl: 4 |
| 663 | 665.8803 | -7.29 | 2 | 37.20 | 59.36 | 0 | 328-339 | K.HPPAVYLLPPAR.E | |
| 389 | 554.2881 | -2.32 | 2 | 28.86 | 54.73 | 0 | 346-355 | R.ESATVTCLVK.G | Carbamidomethyl: 7 |
| 1250 | 802.4093 | -7.11 | 2 | 56.52 | 92.16 | 0 | 356-369 | K.GFSPADISVQWLQR.G | |
| 244 | 456.7583 | -6.09 | 2 | 24.53 | 39.26 | 0 | 370-377 | R.GQLLPQEK.Y | |
| 1175 | 1183.0425 | -11.52 | 2 | 53.85 | 116.72 | 0 | 433-454 | K.STGKPTLYNVSLIMSDTGGTCY.- | Carbamidomethyl: 21 |

| | | | |
|---|---|---|---|
| **Protein 2:** | Serum albumin OS=Mus musculus GN=Alb PE=1 SV=3 | | |
| **Accession:** | ALBU_MOUSE | **Score:** | 565.94 |
| **Database:** | SwissProt | **MW [kDa]:** | 68.60 |
| **Seq. Coverage [%]:** | 19.70 % | **pI:** | 5.75 |
| | | **No. of Peptides:** | 10 |
| **Modification(s):** | Carbamidomethyl | | |

| | | | | | | | | | | | |
|---|---|---|---|---|---|---|---|---|---|---|---|
| 10 | 20 | 30 | 40 | 50 | 60 | 70 | 80 | 90 | 100 | 110 | 120 |
| MKWVTFLLLL | FVSGSAFSRG | VFRREAHKSE | IAHRYNDLGE | QHFKGLVLIA | FSQYLQKCSY | DEHAKLVQEV | TDFAKTCVAD | ESAANCDKSL | HTLFGDKLCA | IPNLRENYGE | LADCCTKQEP |
| 130 | 140 | 150 | 160 | 170 | 180 | 190 | 200 | 210 | 220 | 230 | 240 |
| ERNECFLQHK | DDNPSLPPFE | RPEAEAMCTS | FKENPTTFMG | HYLHEVARRH | PYFYAPELLY | YAEQYNEILT | QCCAEADKES | CLTPKLDGVK | EKALVSSVRQ | RMKCSSMQKF | GERAFKAWAV |
| 250 | 260 | 270 | 280 | 290 | 300 | 310 | 320 | 330 | 340 | 350 | 360 |
| ARLSQTFPNA | DFAEITKLAT | DLTKVNKECC | HGDLLECADD | RAELAKYMCE | NQATISSKLQ | TCCDKPLLKK | AHCLSEVEHD | TMPADLPAIA | ADFVEDQEVC | KNYAEAKDVF | LGTFLYEYSR |
| 370 | 380 | 390 | 400 | 410 | 420 | 430 | 440 | 450 | 460 | 470 | 480 |
| RHPDYSVSLL | LRLAKKYEAT | LEKCCAEANP | PACYGTVLAE | FQPLVEEPKN | LVKTNCDLYE | KLGEYGFQNA | ILVRYTQKAP | QVSTPTLVEA | ARNLGRVGTK | CCTLPEDQRL | PCVEDYLSAI |
| 490 | 500 | 510 | 520 | 530 | 540 | 550 | 560 | 570 | 580 | 590 | 600 |
| LNRVCLLHEK | TPVSEHVTKC | CSGSLVERRP | CFSALTVDET | YVPKEFKAET | FTFHSDICTL | PEKEKQIKKQ | TALAELVKHK | PKATAEQLKT | VMDDFAQFLD | TCCKAADKDT | CFSTEGPNLV |
| 610 | | | | | | | | | | | |
| TRCKDALA | | | | | | | | | | | |

| Cmpd. | m/z meas. | Δ m/z [ppm] | z | Rt [min] | Score | P | Range | Sequence | Modification |
|---|---|---|---|---|---|---|---|---|---|
| 1396 | 740.4231 | -12.25 | 2 | 61.57 | 40.91 | 0 | 45-57 | K.GLVLIAFSQYLQK.C | |
| 596 | 575.3072 | -6.82 | 2 | 35.21 | 62.46 | 0 | 66-75 | K.LVQEVTDFAK.T | |
| 635 | 478.7671 | -8.10 | 2 | 36.41 | 19.93 | 0 | 98-105 | K.LCAIPNLR.E | Carbamidomethyl: 2 |
| 191 | 716.3119 | -5.36 | 2 | 22.54 | 75.58 | 0 | 287-298 | K.YMCENQATISSK.L | Carbamidomethyl: 3 |
| 231 | 459.2380 | -5.58 | 3 | 24.11 | 37.15 | 0 | 299-309 | K.LQTCCDKPLLK.K | Carbamidomethyl: 4, 5 |
| 1026 | 740.3949 | -8.76 | 2 | 48.89 | 91.33 | 0 | 422-434 | K.LGEYGFQNAILVR.Y | |
| 590 | 720.3908 | -7.61 | 2 | 34.99 | 77.97 | 0 | 439-452 | K.APQVSTPTLVEAAR.N | |
| 1480 | 831.9193 | -12.42 | 2 | 64.62 | 89.11 | 0 | 470-483 | R.LPCVEDYLSAILNR.V | Carbamidomethyl: 3 |
| 841 | 628.3111 | -9.54 | 3 | 42.74 | 54.77 | 0 | 509-524 | R.RPCFSALTVDETYVPK.E | Carbamidomethyl: 3 |
| 689 | 486.7852 | -9.49 | 2 | 37.94 | 18.41 | 0 | 550-558 | K.QTALAELVK.H | |

**Protein 3:** Pregnancy zone protein OS=Mus musculus GN=Pzp PE=1 SV=3

**Accession:** PZP_MOUSE

**Database:** SwissProt

**Seq. Coverage [%]:** 12.90 %

**Score:** 563.49

**MW [kDa]:** 165.70

**pI:** 6.24

**No. of Peptides:** 13

**Modification(s):** Carbamidomethyl

| | | | | | | | | | | | |
|---|---|---|---|---|---|---|---|---|---|---|---|
| 10 | 20 | 30 | 40 | 50 | 60 | 70 | 80 | 90 | 100 | 110 | 120 |
| MRRNQLPTPA | FLLLFLLLPR | DATTATAKPQ | YVVLVPSEVY | SGVPEKACVS | LNHVNETVML | SLTLEYAMQQ | TKLLTDQAVD | KDSFYCSPFT | ISGSPLPYTF | ITVEIKGPTQ | RFIKKKSIQI |
| 130 | 140 | 150 | 160 | 170 | 180 | 190 | 200 | 210 | 220 | 230 | 240 |
| IKAESPVFVQ | TDKPIYKPGQ | IVKFRVVSVD | ISFRPLNETF | PVVYIETPKR | NRIFQWQNIH | LAGGLHQLSF | PLSVEPALGI | YKVVVQKDSG | KKIEHSFEVK | EYVLPKFEVI | IKMQKTMAFL |
| 250 | 260 | 270 | 280 | 290 | 300 | 310 | 320 | 330 | 340 | 350 | 360 |
| EEELPITACG | VYTYGKPVPG | LVTLRVCRKY | SRYRSTCHNQ | NSMSICEEFS | QQADDKGCFR | QVVKTKVFQL | RQKGHDMKIE | VEAKIKEEGT | GIELTGIGSC | EIANALSKLK | FTKVNTNYRP |
| 370 | 380 | 390 | 400 | 410 | 420 | 430 | 440 | 450 | 460 | 470 | 480 |
| GLPFSGQVLL | VDEKGKPIPN | KNITSVVSPL | GYLSIFTTDE | HGLANISIDT | SNFTAPFLRV | VVTYKQNHVC | YDNWWLDEFH | TQADHSATLV | FSPSQSYIQL | ELVFGTLACG | QTQEIRIHYL |
| 490 | 500 | 510 | 520 | 530 | 540 | 550 | 560 | 570 | 580 | 590 | 600 |
| LNEDIMKNEK | DLTFYYLIKA | RGSIFNLGSH | VLSLEQGNMK | GVFSLPIQVE | PGMAPEAQLL | IYAILPNEEL | VADAQNFEIE | KCFANKVNLS | FPSAQSLPAS | DTHLKVKAAP | LSLCALTAVD |
| 610 | 620 | 630 | 640 | 650 | 660 | 670 | 680 | 690 | 700 | 710 | 720 |
| QSVLLLKPEA | KLSPQSIYNL | LPGKTVQGAF | FGVPVYKDHE | NCISGEDITH | NGIVYTPKHS | LGDNDAHSIF | QSVGINIFTN | SKIHKPRFCQ | EFQHYPAMGG | VAPQALAVAA | SGPGSSFRAM |
| 730 | 740 | 750 | 760 | 770 | 780 | 790 | 800 | 810 | 820 | 830 | 840 |
| GVPMMGLDYS | DEINQVVEVR | ETVRKYFPET | WIWDLVPLDV | SGDGELAVKV | PDTITEWKAS | AFCLSGTTGL | GLSSTISLQA | FQPFFLELTL | PYSVVRGEAF | TLKATVLNYM | SHCIQIRVDL |
| 850 | 860 | 870 | 880 | 890 | 900 | 910 | 920 | 930 | 940 | 950 | 960 |
| EISPDFLAVP | VGGHENSHCI | CGNERKTVSW | AVTPKSLGEV | NFTATAEALQ | SPELCGNKLT | EVPALVHKDT | VVKSVIVEPE | GIEKEQTYNT | LLCPQDTELQ | DNWSLELPPN | VVEGSARATH |
| 970 | 980 | 990 | 1000 | 1010 | 1020 | 1030 | 1040 | 1050 | 1060 | 1070 | 1080 |
| SVLGDILGSA | MQNLQNLLQM | PYGCGEQNMV | LFVPNIYVLN | YLNETQQLTE | AIKSKAINYL | ISGYQRQLNY | QHSDGSYSTF | GNHGGGNTPG | NTWLTAFVLK | AFAQAQSHIF | IEKTHITNAF |
| 1090 | 1100 | 1110 | 1120 | 1130 | 1140 | 1150 | 1160 | 1170 | 1180 | 1190 | 1200 |
| NWLSMKQKEN | GCFQQSGYLL | NNAMKGGVDD | EVTLSAYITI | ALLEMPLPVT | HSAVRNALFC | LETAWASISQ | SQESHVYTKA | LLAYAFALAG | NKAKRSELLE | SLNKDAVKEE | DSLHWQRPGD |
| 1210 | 1220 | 1230 | 1240 | 1250 | 1260 | 1270 | 1280 | 1290 | 1300 | 1310 | 1320 |
| VQKVKALSFY | QPRAPSAEVE | MTAYVLLAYL | TSESSRPTRD | LSSSDLSTAS | KIVKWISKQQ | NSHGGFSSTQ | DTVVALQALS | KYGAATFTRS | QKEVLVTIES | SGTFSKTFHV | NSGNRLLLQE |
| 1330 | 1340 | 1350 | 1360 | 1370 | 1380 | 1390 | 1400 | 1410 | 1420 | 1430 | 1440 |
| VRLPDLPGNY | VTKGSGSGCV | YLQTSLKYNI | LPVADGKAPF | ALQVNTLPLN | FDKAGDHRTF | QIRINVSYTG | ERPSSNMVIV | DVKMVSGFIP | MKPSVKKLQD | QPNIQRTEVN | TNHVLIYIEK |
| 1450 | 1460 | 1470 | 1480 | 1490 | 1500 | | | | | | |
| LTNQTLGFSF | AVEQDIPVKN | LKPAPIKVYD | YYETDEFTVE | EYSAPFSDGS | EQGNA | | | | | | |

| Cmpd. | m/z meas. | Δ m/z [ppm] | z | Rt [min] | Score | P | Range | Sequence | Modification |
|---|---|---|---|---|---|---|---|---|---|
| 236 | 501.7734 | -7.01 | 2 | 24.25 | 13.58 | 0 | 73-81 | K.LLTDQAVDK.D | |
| 1079 | 782.7462 | -9.02 | 3 | 50.63 | 56.01 | 0 | 354-374 | K.VNTNYRPGLPFSGQVLLVDEK.G | |
| 1193 | 677.6756 | -10.07 | 3 | 54.43 | 29.09 | 0 | 502-520 | R.GSIFNLGSHVLSLEQGNMK.G | |

| Cmpd. | m/z meas. | Δ m/z [ppm] | z | Rt [min] | Score | P | Range | Sequence | Modification |
|---|---|---|---|---|---|---|---|---|---|
| 1420 | 836.7986 | -11.68 | 3 | 62.44 | 75.23 | 0 | 588-611 | K.AAPLSLCALTAVDQSVLLLKPEAK.L | Carbamidomethyl: 7 |
| 1185 | 715.3994 | -9.42 | 2 | 54.16 | 11.02 | 0 | 612-624 | K.LSPQSIYNLLPGK.T | |
| 1063 | 706.8750 | -10.28 | 2 | 50.10 | 78.47 | 0 | 625-637 | K.TVQGAFFGVPVYK.D | |
| 636 | 544.7796 | -9.48 | 2 | 36.42 | 47.62 | 0 | 770-778 | K.VPDTITEWK.A | |
| 787 | 649.3414 | -11.16 | 2 | 40.96 | 36.66 | 0 | 1016-1026 | K.AINYLISGYQR.Q | |
| 565 | 497.2616 | -6.42 | 3 | 34.26 | 51.82 | 0 | 1061-1073 | K.AFAQAQSHIFIEK.T | |
| 1421 | 924.0991 | -11.27 | 3 | 62.45 | 25.56 | 0 | 1136-1159 | R.NALFCLETAWASISQSQESHVYTK.A | Carbamidomethyl: 5 |
| 714 | 608.8262 | -9.91 | 2 | 38.74 | 56.95 | 0 | 1323-1333 | R.LPDLPGNYVTK.G | |
| 705 | 545.2951 | -10.09 | 2 | 38.40 | 32.56 | 0 | 1348-1357 | K.YNILPVADGK.A | |
| 1301 | 894.4776 | -11.86 | 2 | 58.16 | 48.92 | 0 | 1358-1373 | K.APFALQVNTLPLNFDK.A | |

**Protein 4:** Fibronectin OS=Mus musculus GN=Fn1 PE=1 SV=4

**Accession:** FINC_MOUSE

**Database:** SwissProt

**Seq. Coverage [%]:** 3.20 %

**Score:** 254.35

**MW [kDa]:** 272.40

**pI:** 5.38

**No. of Peptides:** 6

**Modification(s):** Carbamidomethyl

| | | | | | | | | | | | |
|---|---|---|---|---|---|---|---|---|---|---|---|
| 10 | 20 | 30 | 40 | 50 | 60 | 70 | 80 | 90 | 100 | 110 | 120 |
| MLRGPGPGRL | LLLAVLCLGT | SVRCTEAGKS | KRQAQQIVQP | QSPVAVSQSK | PGCFDNGKHY | QINQQWERTY | LGNALVCTCY | GGSRGFNCES | KPEPEETCFD | KYTGNTYKVG | DTYERPKDSM |
| 130 | 140 | 150 | 160 | 170 | 180 | 190 | 200 | 210 | 220 | 230 | 240 |
| IWDCTCIGAG | RGRISCTIAN | RCHEGGQSYK | IGDKWRRPHE | TGGYMLECLC | LGNGKGEWTC | KPIAEKCFDH | AAGTSYVVGE | TWEKPYQGWM | MVDCTCLGEG | NGRITCTSRN | RCNDQDTRTS |
| 250 | 260 | 270 | 280 | 290 | 300 | 310 | 320 | 330 | 340 | 350 | 360 |
| YRIGDTWSKK | DNRGNLLQCV | CTGNGRGEWK | CERHALQSAS | AGSGSFTDVR | TAIYQPQTHP | QPAPYGHCVT | DSGVVYSVGM | QWLKSQGNKQ | MLCTCLGNGV | SCQETAVTQT | YGGNSNGEPC |
| 370 | 380 | 390 | 400 | 410 | 420 | 430 | 440 | 450 | 460 | 470 | 480 |
| VLPFTYNGRT | FYSCTTEGRQ | DGHLWCSTTS | NYEQDQKYSF | CTDHAVLVQT | RGGNSNGALC | HFPFLYNNRN | YTDCTSEGRR | DNMKWCGTTQ | NYDADQKFGF | CPMAAHEEIC | TTNEGVMYRI |
| 490 | 500 | 510 | 520 | 530 | 540 | 550 | 560 | 570 | 580 | 590 | 600 |
| GDQWDKQHDL | GHMMRCTCVG | NGRGEWACIP | YSQLRDQCIV | DDITYNVNDT | FHKRHEEGHM | LNCTCFGQGR | GRWKCDPIDQ | CQDSETRTFY | QIGDSWEKFV | HGVRYQCYCY | GRGIGEWHCQ |
| 610 | 620 | 630 | 640 | 650 | 660 | 670 | 680 | 690 | 700 | 710 | 720 |
| PLQTYPGTTG | PVQVIITETP | SQPNSHPIQW | NAPEPSHITK | YILRWRPKTS | TGRWKEATIP | GHLNSYTIKG | LTPGVIYEGQ | LISIQQYGHR | EVTRFDFTTS | ASTPVTSNTV | TGETAPYSPV |
| 730 | 740 | 750 | 760 | 770 | 780 | 790 | 800 | 810 | 820 | 830 | 840 |
| VATSESVTEI | TASSFVVSWV | SASDTVSGFR | VEYELSEEGD | EPQYLDLPST | ATSVNIPDLL | PGRKYIVNVY | QISEEGKQSL | ILSTSQTTAP | DAPPDPTVDQ | VDDTSIVVRW | SRPQAPITGY |
| 850 | 860 | 870 | 880 | 890 | 900 | 910 | 920 | 930 | 940 | 950 | 960 |
| RIVYSPSVEG | SSTELNLPET | ANSVTLSDLQ | PGVQYNITIY | AVEENQESTP | VFIQQETTGT | PRSDNVPPPT | DLQFVELTDV | KVTIMWTPPD | SVVSGYRVEV | LPVSLPGEHG | QRLPVNRNTF |
| 970 | 980 | 990 | 1000 | 1010 | 1020 | 1030 | 1040 | 1050 | 1060 | 1070 | 1080 |
| AEITGLSPGV | TYLFKVFAVH | QGRESNPLTA | QQTTKLDAPT | NLQFVNETDR | TVLVTWTPPR | ARIAGYRLTA | GLTRGGQPKQ | YNVGPLASKY | PLRNLQPGSE | YTVTLVAVKG | NQQSPKATGV |
| 1090 | 1100 | 1110 | 1120 | 1130 | 1140 | 1150 | 1160 | 1170 | 1180 | 1190 | 1200 |
| FTTLQPLRSI | PPYNTEVTET | TIVITWTPAP | RIGFKLGVRP | SQGGEAPREV | TSDSGSIVVS | GLTPGVEYTY | TIQVLRDGQE | RDAPIVNRVV | TPLSPPTNLH | LEANPDTGVL | TVSWERSTTP |
| 1210 | 1220 | 1230 | 1240 | 1250 | 1260 | 1270 | 1280 | 1290 | 1300 | 1310 | 1320 |
| DITGYRITTT | PTNGQQGTSL | EEVVHADQSS | CTFENLNPGL | EYNVSVYTVK | DDKESAPISD | TVVPEVPQLT | DLSFVDITDS | SIGLRWTPLN | SSTIIGYRIT | VVAAGEGIPI | FEDFVDSSVG |
| 1330 | 1340 | 1350 | 1360 | 1370 | 1380 | 1390 | 1400 | 1410 | 1420 | 1430 | 1440 |
| YYTVTGLEPG | IDYDISVITL | INGGESAPTT | LTQQTAVPPP | TDLRFTNIGP | DTMRVTWAPP | PSIELTNLLV | RYSPVKNEED | VAELSISPSD | NAVVLTNLLP | GTEYLVSVSS | VYEQHESIPL |
| 1450 | 1460 | 1470 | 1480 | 1490 | 1500 | 1510 | 1520 | 1530 | 1540 | 1550 | 1560 |
| RGRQKTGLDS | PTGFDSSDIT | ANSFTVHWVA | PRAPITGYII | RHHAEHSVGR | PRQDRVPPSR | NSITLTNLNP | GTEYVVSIIA | VNGREESPPL | IGQQATVSDI | PRDLEVIAST | PTSLLISWEP |
| 1570 | 1580 | 1590 | 1600 | 1610 | 1620 | 1630 | 1640 | 1650 | 1660 | 1670 | 1680 |
| PAVSVRYYRI | TYGETGGNSP | VQEFTVPGSK | STATINNIKP | GADYTITLYA | VTGRGDSPAS | SKPVSINYKT | EIDKPSQMQV | TDVQDNSISV | RWLPSTSPVT | GYRVTTTPKN | GLGPSKTKTA |
| 1690 | 1700 | 1710 | 1720 | 1730 | 1740 | 1750 | 1760 | 1770 | 1780 | 1790 | 1800 |
| SPDQTEMTIE | GLQPTVEYVV | SVYAQNRNGE | SQPLVQTAVT | NIDRPKGLAF | TDVDVDSIKI | AWESPQGQVS | RYRVTYSSPE | DGIRELFPAP | DGEDDTAELQ | GLRPGSEYTV | SVVALHDDME |
| 1810 | 1820 | 1830 | 1840 | 1850 | 1860 | 1870 | 1880 | 1890 | 1900 | 1910 | 1920 |
| SQPLIGIQST | AIPAPTNLKF | SQVTPTSFTA | QWIAPSVQLT | GYRVRVNPKE | KTGPMKEINL | SPDSSSVIVS | GLMVATKYEV | SVYALKDTLT | SRPAQGVITT | LENVSPPRRA | RVTDATETTI |
| 1930 | 1940 | 1950 | 1960 | 1970 | 1980 | 1990 | 2000 | 2010 | 2020 | 2030 | 2040 |
| TISWRTKTET | ITGFQVDAIP | ANGQTPVQRS | ISPDVRSYTI | TGLQPGTDYK | IHLYTLNDNA | RSSPVIIDAS | TAIDAPSNLR | FLTTTPNSLL | VSWQAPRARI | TGYIIKYEKP | GSPPREVVPR |
| 2050 | 2060 | 2070 | 2080 | 2090 | 2100 | 2110 | 2120 | 2130 | 2140 | 2150 | 2160 |

| Cmpd. | m/z meas. | Δ m/z [ppm] | z | Rt [min] | Score | P | Range | Sequence | Modification |
|---|---|---|---|---|---|---|---|---|---|
| 240 | 611.2577 | -10.04 | 2 | 24.39 | 30.60 | 0 | 370-379 | R.TFYSCTTEGR.Q | Carbamidomethyl: 5 |
| 1401 | 979.5056 | -11.80 | 2 | 61.74 | 85.76 | 0 | 958-975 | R.NTFAEITGLSPGVTYLFK.V | |
| 931 | 652.3660 | -9.32 | 2 | 45.63 | 66.87 | 0 | 1077-1088 | K.ATGVFTTLQPLR.S | |
| 86 | 441.9068 | -5.18 | 3 | 18.23 | 11.55 | 0 | 1116-1128 | K.LGVRPSQGGEAPR.E | |
| 1433 | 953.5292 | -15.01 | 2 | 62.88 | 42.41 | 0 | 1375-1391 | R.VTWAPPPSIELTNLLVR.Y | |
| 724 | 502.2953 | -10.22 | 2 | 39.01 | 17.16 | 0 | 1473-1481 | R.APITGYIIR.H | |

**Protein 5:** Ig kappa chain C region OS=Mus musculus PE=1 SV=1

**Accession:** IGKC_MOUSE | **Score:** 221.81

**Database:** SwissProt | **MW [kDa]:** 11.80

**Seq. Coverage [%]:** 34.00 % | **pI:** 5.23

**No. of Peptides:** 4

**Modification(s):** Oxidation

```
         10         20         30         40         50         60         70         80         90        100        110
ADAAPTVSIF PPSSEQLTSG GASVVCFLNN FYPKDINVKW KIDGSERQNG VLNSWTDQDS KDSTYSMSST LTLTKDEYER HNSYTCEATH KTSTSPIVKS FNRNEC
```

| Cmpd. | m/z meas. | Δ m/z [ppm] | z | Rt [min] | Score | P | Range | Sequence | Modification |
|---|---|---|---|---|---|---|---|---|---|
| 604 | 796.3642 | -8.49 | 2 | 35.46 | 96.98 | 0 | 48-61 | R.QNGVLNSWTDQDSK.D | |
| 773 | 767.8647 | -5.41 | 2 | 40.54 | 98.43 | 0 | 62-75 | K.DSTYSMSSTLTLTK.D | |
| 484 | 775.8607 | -7.31 | 2 | 31.78 | 87.23 | 0 | 62-75 | K.DSTYSMSSTLTLTK.D | Oxidation: 6 |
| 94 | 416.7409 | -3.57 | 2 | 18.53 | 26.40 | 0 | 92-99 | K.TSTSPIVK.S | |

**Protein 6:** Ig heavy chain V region AC38 205.12 OS=Mus musculus PE=1 SV=1

| | | | |
|---|---|---|---|
| **Accession:** | HVM51_MOUSE | **Score:** | 214.94 |
| **Database:** | SwissProt | **MW [kDa]:** | 12.90 |
| **Seq. Coverage [%]:** | 36.40 % | **pI:** | 6.85 |
| | | **No. of Peptides:** | 3 |

**Modification(s):** Carbamidomethyl

10 20 30 40 50 60 70 80 90 100 110 120

EVQLQQSGPE LVKPGASVKI SCKASGYTFT DYYMNWVKQS HGKSLEWIGD INPNNGGTSY NQKFKGKATL TVDKSSSATY MELRSLTSED SAVYYCARGY GYDPFDVWGT GTTVTVSS

| Cmpd. | m/z meas. | Δ m/z [ppm] | z | Rt [min] | Score | P | Range | Sequence | Modification |
|---|---|---|---|---|---|---|---|---|---|
| 513 | 665.3638 | -7.45 | 3 | 32.63 | 67.25 | 0 | 1-19 | -.EVQLQQSGPELVKPGASVK.I | |
| 463 | 572.7642 | -7.94 | 2 | 31.06 | 41.30 | 0 | 75-84 | K.SSSATYMELR.S | |
| 533 | 811.3549 | -8.43 | 2 | 33.22 | 106.39 | 0 | 85-98 | R.SLTSEDSAVYYCAR.G | Carbamidomethyl: 12 |

**Protein 7:** Hemoglobin subunit beta-1 OS=Mus musculus GN=Hbb-b1 PE=1 SV=2

| | | | |
|---|---|---|---|
| **Accession:** | HBB1_MOUSE | **Score:** | 154.66 |
| **Database:** | SwissProt | **MW [kDa]:** | 15.80 |
| **Seq. Coverage [%]:** | 25.90 % | **pI:** | 7.13 |
| | | **No. of Peptides:** | 3 |

10 20 30 40 50 60 70 80 90 100 110 120

MVHLTDAEKA AVSCLWGKVN SDEVGGEALG RLLVVYPWTQ RYFDSFGDLS SASAIMGNAK VKAHGKKVIT AFNDGLNHLD SLKGTFASLS ELHCDKLHVD PENFRLLGNM IVIVLGHHLG

130 140 150

KDFTPAAQAA FQKVVAGVAT ALAHKYH

| Cmpd. | m/z meas. | Δ m/z [ppm] | z | Rt [min] | Score | P | Range | Sequence | Modification |
|---|---|---|---|---|---|---|---|---|---|
| 1100 | 637.8595 | -10.81 | 2 | 51.31 | 51.31 | 0 | 32-41 | R.LLVVYPWTQR.Y | |
| 959 | 586.3069 | -9.46 | 3 | 46.65 | 53.22 | 0 | 68-83 | K.VITAFNDGLNHLDSLK.G | |

| Cmpd. | m/z meas. | Δ m/z [ppm] | z | Rt [min] | Score | P | Range | Sequence | Modification |
|---|---|---|---|---|---|---|---|---|---|
| 582 | 647.8194 | -8.55 | 2 | 34.73 | 50.13 | 0 | 122-133 | K.DFTPAAQAAFQK.V | |

| | | | |
|---|---|---|---|
| **Protein 8:** | Hemoglobin subunit alpha OS=Mus musculus GN=Hba PE=1 SV=2 | | |
| **Accession:** | HBA_MOUSE | **Score:** | 139.16 |
| **Database:** | SwissProt | **MW [kDa]:** | 15.10 |
| **Seq. Coverage [%]:** | 19.00 % | **pI:** | 7.97 |
| | | **No. of Peptides:** | 2 |

```
        10         20         30         40         50         60         70         80         90        100        110        120
MVLSGEDKSN IKAAWGKIGG HGAEYGAEAL ERMFASFPTT KTYFPHFDVS HGSAQVKGHG KKVADALASA AGHLDDLPGA LSALSDLHAH KLRVDPVNFK LLSHCLLVTL ASHHPADFTP
       130        140        150
AVHASLDKFL ASVSTVLTSK YR
```

| Cmpd. | m/z meas. | Δ m/z [ppm] | z | Rt [min] | Score | P | Range | Sequence | Modification |
|---|---|---|---|---|---|---|---|---|---|
| 348 | 510.5784 | -8.85 | 3 | 27.61 | 73.71 | 0 | 18-32 | K.IGGHGAEYGAEALER.M | |
| 904 | 626.8552 | -9.27 | 2 | 44.79 | 65.45 | 0 | 129-140 | K.FLASVSTVLTSK.Y | |

| | | | |
|---|---|---|---|
| **Protein 9:** | Keratin, type II cytoskeletal 1 OS=Mus musculus GN=Krt1 PE=1 SV=4 | | |
| **Accession:** | K2C1_MOUSE | **Score:** | 138.22 |
| **Database:** | SwissProt | **MW [kDa]:** | 65.60 |
| **Seq. Coverage [%]:** | 3.60 % | **pI:** | 8.39 |
| | | **No. of Peptides:** | 2 |

| | | | | | | | | | | | |
|---|---|---|---|---|---|---|---|---|---|---|---|
| 10 | 20 | 30 | 40 | 50 | 60 | 70 | 80 | 90 | 100 | 110 | 120 |
| MSLQCSSRSL | CRGGGGSRNF | SSGSAGLVSF | QRRSTSSSMR | RSGGGGGGRF | SGGGFCGSSG | SGFGSKSLMN | LGGGRSISKS | VAGGGGSFCG | GFGGGSYGGG | GFGGGSYGGG | GFGGGSFGGG |
| 130 | 140 | 150 | 160 | 170 | 180 | 190 | 200 | 210 | 220 | 230 | 240 |
| GFGGSGFGGG | LGGGGGFGSG | GGFGGGRFGS | MGPVCPPGGI | QEVTINQSLL | QPLNVEVDPQ | IQKVKSQERE | QIKSLNDKFA | SFIDKVRFLE | QQNQVLQTKW | ELLQQVDTTT | RTQNLDPFFE |
| 250 | 260 | 270 | 280 | 290 | 300 | 310 | 320 | 330 | 340 | 350 | 360 |
| NYISILRRKV | DSLKSDQSRM | DSELKNMQDL | VEEYRTKYED | EINKRTNAEN | EFVTIKKDVD | SAYMTKVELQ | AKADALQQDI | DFFSALYQME | MSQMQTQISE | TNVVLSMDNN | RSLDLDGIIS |
| 370 | 380 | 390 | 400 | 410 | 420 | 430 | 440 | 450 | 460 | 470 | 480 |
| EVKAQYDSIC | QRSKAEAETF | YQSKYEELQI | TAGKHGDSVR | NTKMEISELN | RMIQRLRSEI | DGCKKQISQI | QQNINDAEQR | GEKALKDAQN | KLNEIEDALS | QCKEDLARLL | RDFQELMNTK |
| 490 | 500 | 510 | 520 | 530 | 540 | 550 | 560 | 570 | 580 | 590 | 600 |
| LALDMEIATY | KKLLEGEEIR | MSGECTPNVS | VSVSTSHTSM | SGSSSRGGGS | GGGRYGGGGS | YGGGSGGGSY | GGSSGGGGSG | GSYGGGSGGG | SYGGGSGGGS | SGSHRGGSGG | GGGSSGGSYG |
| 610 | 620 | 630 | 640 | | | | | | | | |
| GSSGGGRGGS | SSGGGGVKSS | GSSTVKFVST | SYSRGTK | | | | | | | | |

| Cmpd. | m/z meas. | Δ m/z [ppm] | z | Rt [min] | Score | P | Range | Sequence | Modification |
|---|---|---|---|---|---|---|---|---|---|
| 492 | 738.3906 | -7.67 | 2 | 32.05 | 76.19 | 0 | 208-219 | R.FLEQQNQVLQTK.W | |
| 580 | 633.3170 | -8.21 | 2 | 34.70 | 62.03 | 0 | 286-296 | R.TNAENEFVTIK.K | |

**Protein 10:** C4b-binding protein OS=Mus musculus GN=C4bpa PE=1 SV=3

**Accession:** C4BPA_MOUSE

**Database:** SwissProt

**Seq. Coverage [%]:** 8.50 %

**Score:** 123.83

**MW [kDa]:** 51.50

**pI:** 6.83

**No. of Peptides:** 3

**Modification(s):** Carbamidomethyl, Oxidation

| | | | | | | | | | | | |
|---|---|---|---|---|---|---|---|---|---|---|---|
| 10 | 20 | 30 | 40 | 50 | 60 | 70 | 80 | 90 | 100 | 110 | 120 |
| MCAKQQQTLL | PTRAAHGRLH | RNRDAVAWPF | STLCRVSGPT | LFQMTFTAAL | WVAVFGKCGP | PPAIPNALPA | SDVNRTDFES | HTTLKYECLP | GYGRGISRMM | VYCKPSGEWE | ISVSCAKKHC |
| 130 | 140 | 150 | 160 | 170 | 180 | 190 | 200 | 210 | 220 | 230 | 240 |
| RNPGYLDNGY | VNGETITFGS | QIEFSCQEGF | ILVGSSTSSC | EVRGKGVAWS | NPFPECVIVK | CGPPPDISNG | KHSGTEDFYP | YNHGISYTCD | PGFRLVGSPF | IGCTVVNKTV | PVWSSSPPTC |
| 250 | 260 | 270 | 280 | 290 | 300 | 310 | 320 | 330 | 340 | 350 | 360 |
| EKIICSQPNI | LHGVIVSGYK | ATYTHRDSVR | LACLNGTVLR | GRHVIECQGN | GNWSSLPTCE | FDCDLPPAIV | NGYYTSMVYS | KITLVTYECD | KGYRLVGKAI | ISCSFSKWKG | TAPQCKALCQ |
| 370 | 380 | 390 | 400 | 410 | 420 | 430 | 440 | 450 | 460 | 470 | |
| KPEVGNGTLS | DEKDQYVESE | NVTIQCDSGF | AMLGSQSISC | SESGTWYPEV | PRCEQEASED | LKPALTGNKT | MQYVPNSHDV | KMALEIYKLT | LEVELLQLQI | QKEKHTEAH | |

| Cmpd. | m/z meas. | Δ m/z [ppm] | z | Rt [min] | Score | P | Range | Sequence | Modification |
|---|---|---|---|---|---|---|---|---|---|
| 899 | 666.6943 | -7.84 | 3 | 44.59 | 35.97 | 0 | 243-260 | K.IICSQPNILHGVIVSGYK.A | Carbamidomethyl: 3 |
| 581 | 621.3025 | -8.52 | 2 | 34.72 | 58.80 | 0 | 322-331 | K.ITLVTYECDK.G | Carbamidomethyl: 8 |
| 129 | 478.8904 | -8.10 | 3 | 19.71 | 29.06 | 0 | 430-441 | K.TMQYVPNSHDVK.M | Oxidation: 2 |

| | | | |
|---|---|---|---|
| **Protein 11:** | Immunoglobulin J chain OS=Mus musculus GN=Jchain PE=1 SV=4 | | |
| **Accession:** | IGJ_MOUSE | **Score:** | 117.61 |
| **Database:** | SwissProt | **MW [kDa]:** | 18.00 |
| **Seq. Coverage [%]:** | 16.40 % | **pI:** | 4.74 |
| | | **No. of Peptides:** | 2 |
| **Modification(s):** | Carbamidomethyl | | |

```
         10         20         30         40         50         60         70         80         90        100        110        120
MKTHLLLWGV LAIFVKAVLV TGDDEATILA DNKCMCTRVT SRIIPSTEDP NEDIVERNIR IVVPLNNREN ISDPTSPLRR NFVYHLSDVC KKCDPVEVEL EDQVVTATQS NICNEDDGVP
        130        140        150        160
ETCYMYDRNK CYTTMVPLRY HGETKMVQAA LTPDSCYPD
```

| Cmpd. | m/z meas. | Δ m/z [ppm] | z | Rt [min] | Score | P | Range | Sequence | Modification |
|---|---|---|---|---|---|---|---|---|---|
| 540 | 863.9219 | -7.41 | 2 | 33.48 | 89.76 | 0 | 43-57 | R.IIPSTEDPNEDIVER.N | |
| 595 | 461.2207 | -6.69 | 3 | 35.20 | 27.85 | 0 | 81-91 | R.NFVYHLSDVCK.K | Carbamidomethyl: 10 |

| | | | |
|---|---|---|---|
| **Protein 12:** | Thrombospondin-1 OS=Mus musculus GN=Thbs1 PE=1 SV=1 | | |
| **Accession:** | TSP1_MOUSE | **Score:** | 113.18 |
| **Database:** | SwissProt | **MW [kDa]:** | 129.60 |
| **Seq. Coverage [%]:** | 2.00 % | **pI:** | 4.72 |
| | | **No. of Peptides:** | 2 |

| | | | | | | | | | | | |
|---|---|---|---|---|---|---|---|---|---|---|---|
| 10 | 20 | 30 | 40 | 50 | 60 | 70 | 80 | 90 | 100 | 110 | 120 |
| MELLRGLGVL | FLLHMCGSNR | IPESGGDNGV | FDIFELIGGA | RRGPGRRLVK | GQDLSSPAFR | IENANLIPAV | PDDKFQDLLD | AVWADKGFIF | LASLRQMKKT | RGTLLAVERK | DNTGQIFSVV |
| 130 | 140 | 150 | 160 | 170 | 180 | 190 | 200 | 210 | 220 | 230 | 240 |
| SNGKAGTLDL | SLSLPGKQQV | VSVEEALLAT | GQWKSITLFV | QEDRAQLYID | CDKMESAELD | VPIQSIFTRD | LASVARLRVA | KGDVNDNFQG | VLQNVRFVFG | TTPEDILRNK | GCSSSTNVLL |
| 250 | 260 | 270 | 280 | 290 | 300 | 310 | 320 | 330 | 340 | 350 | 360 |
| TLDNNVVNGS | SPAIRTNYIG | HKTKDLQAIC | GLSCDELSSM | VLELKGLRTI | VTTLQDSIRK | VTEENRELVS | ELKRPPLCFH | NGVQYKNNEE | WTVDSCTECH | CQNSVTICKK | VSCPIMPCSN |
| 370 | 380 | 390 | 400 | 410 | 420 | 430 | 440 | 450 | 460 | 470 | 480 |
| ATVPDGECCP | RCWPSDSADD | GWSPWSEWTS | CSATCGNGIQ | QRGRSCDSLN | NRCEGSSVQT | RTCHIQECDK | RFKQDGGWSH | WSPWSSCSVT | CGDGVITRIR | LCNSPSPQMN | GKPCEGEARE |
| 490 | 500 | 510 | 520 | 530 | 540 | 550 | 560 | 570 | 580 | 590 | 600 |
| TKACKKDACP | INGGWGPWSP | WDICSVTCGG | GVQRRSRLCN | NPTPQFGGKD | CVGDVTENQV | CNKQDCPIDG | CLSNPCFAGA | KCTSYPDGSW | KCGACPPGYS | GNGIQCKDVD | ECKEVPDACF |
| 610 | 620 | 630 | 640 | 650 | 660 | 670 | 680 | 690 | 700 | 710 | 720 |
| NHNGEHRCKN | TDPGYNCLPC | PPRFTGSQPF | GRGVEHAMAN | KQVCKPRNPC | TDGTHDCNKN | AKCNYLGHYS | DPMYRCECKP | GYAGNGIICG | EDTDLDGWPN | ENLVCVANAT | YHCKKDNCPN |
| 730 | 740 | 750 | 760 | 770 | 780 | 790 | 800 | 810 | 820 | 830 | 840 |
| LPNSGQEDYD | KDGIGDACDD | DDDNDKIPDD | RDNCPFHYNP | AQYDYDRDDV | GDRCDNCPYN | HNPDQADTDK | NGEGDACAVD | IDGDGILNER | DNCQYVYNVD | QRDTDMDGVG | DQCDNCPLEH |
| 850 | 860 | 870 | 880 | 890 | 900 | 910 | 920 | 930 | 940 | 950 | 960 |
| NPDQLDSDSD | LIGDTCDNNQ | DIDEDGHQNN | LDNCPYVPNA | NQADHDKDGK | GDACDHDDDN | DGIPDDRDNC | RLVPNPDQKD | SDGDGRGDAC | KDDFDHDNVP | DIDDICPENF | DISETDFRRF |
| 970 | 980 | 990 | 1000 | 1010 | 1020 | 1030 | 1040 | 1050 | 1060 | 1070 | 1080 |
| QMIPLDPKGT | SQNDPNWVVR | HQGKELVQTV | NCDPGLAVGY | DEFNAVDFSG | TFFINTERDD | DYAGFVFGYQ | SSSRFYVVMW | KQVTQSYWDT | NPTRAQGYSG | LSVKVVNSTT | GPGEHLRNAL |
| 1090 | 1100 | 1110 | 1120 | 1130 | 1140 | 1150 | 1160 | 1170 | 1180 | | |
| WHTGNTPGQV | RTLWHDPRHI | GWKDFTAYRW | RLSHRPKTGY | IRVVMYEGKK | IMADSGPIYD | KTYAGGRLGL | FVFSQEMVFF | SDMKYECRDS | | | |

| Cmpd. | m/z meas. | Δ m/z [ppm] | z | Rt [min] | Score | P | Range | Sequence | Modification |
|---|---|---|---|---|---|---|---|---|---|
| 1181 | 697.8620 | -10.57 | 2 | 54.02 | 42.20 | 0 | 217-228 | R.FVFGTTPEDILR.N | |
| 912 | 623.8471 | -10.64 | 2 | 45.04 | 70.98 | 0 | 289-299 | R.TIVTTLQDSIR.K | |

| | | | |
|---|---|---|---|
| **Protein 13:** | Ig kappa chain V-II region 7S34.1 OS=Mus musculus PE=1 SV=1 | | |
| **Accession:** | KV2A6_MOUSE | **Score:** | 103.16 |
| **Database:** | SwissProt | **MW [kDa]:** | 12.50 |
| **Seq. Coverage [%]:** | 11.50 % | **pI:** | 8.89 |
| | | **No. of Peptides:** | 1 |

| 10 | 20 | 30 | 40 | 50 | 60 | 70 | 80 | 90 | 100 | 110 | 120 |
|---|---|---|---|---|---|---|---|---|---|---|---|
| DIVMTQTAPS | ALVTPGESVS | ISCRSSKSLL | HSNGNTYLYW | FLQRPGQCPQ | LLIYRMSNLA | SGVPDRFSGS | GSGTAFTLRI | SRVEAEDVGV | YYCMQQREYP | YTFGGGTKLE | IKR |

| Cmpd. | m/z meas. | Δ m/z [ppm] | z | Rt [min] | Score | P | Range | Sequence | Modification |
|---|---|---|---|---|---|---|---|---|---|
| 613 | 644.3149 | -8.00 | 2 | 35.71 | 103.16 | 0 | 67-79 | R.FSGSGSGTAFTLR.I | |

**Protein 14:** CD5 antigen-like OS=Mus musculus GN=Cd5l PE=1 SV=3

**Accession:** CD5L_MOUSE

**Database:** SwissProt

**Seq. Coverage [%]:** 8.80 %

**Score:** 95.89

**MW [kDa]:** 38.80

**pI:** 5.01

**No. of Peptides:** 3

**Modification(s):** Carbamidomethyl

| 10 | 20 | 30 | 40 | 50 | 60 | 70 | 80 | 90 | 100 | 110 | 120 |
|---|---|---|---|---|---|---|---|---|---|---|---|
| MAPLFNLMLA | ILSIFVGSCF | SESPTKVQLV | GGAHRCEGRV | EVEHNGQWGT | VCDDGWDRRD | VAVVCRELNC | GAVIQTPRGA | SYQPPASEQR | VLIQGVDCNG | TEDTLAQCEL | NYDVFDCSHE |
| 130 | 140 | 150 | 160 | 170 | 180 | 190 | 200 | 210 | 220 | 230 | 240 |
| EDAGAQCENP | DSDLLFIPED | VRLVDGPGHC | QGRVEVLHQS | QWSTVCKAGW | NLQVSKVVCR | QLGCGRALLT | YGSCNKNTQG | KGPIWMGKMS | CSGQEANLRS | CLLSRLENNC | THGEDTWMEC |
| 250 | 260 | 270 | 280 | 290 | 300 | 310 | 320 | 330 | 340 | 350 | 360 |
| EDPFELKLVG | GDTPCSGRLE | VLHKGSWGSV | CDDNWGEKED | QVVCKQLGCG | KSLHPSPKTR | KIYGPGAGRI | WLDDVNCSGK | EQSLEFCRHR | LWGYHDCTHK | EDVEVICTDF | DV |

| Cmpd. | m/z meas. | Δ m/z [ppm] | z | Rt [min] | Score | P | Range | Sequence | Modification |
|---|---|---|---|---|---|---|---|---|---|
| 594 | 501.7679 | -8.11 | 2 | 35.12 | 20.09 | 0 | 168-176 | K.AGWNLQVSK.V | |
| 115 | 626.7707 | -5.33 | 2 | 19.29 | 12.90 | 0 | 209-219 | K.MSCSGQEANLR.S | Carbamidomethyl: 3 |
| 722 | 653.8034 | -7.72 | 2 | 38.98 | 62.90 | 0 | 310-320 | R.IWLDDVNCSGK.E | Carbamidomethyl: 8 |

**Protein 15:** Ig kappa chain V-II region 26-10 OS=Mus musculus PE=1 SV=1

| | | | |
|---|---|---|---|
| **Accession:** | KV2A7_MOUSE | **Score:** | 91.85 |
| **Database:** | SwissProt | **MW [kDa]:** | 12.30 |
| **Seq. Coverage [%]:** | 11.50 % | **pI:** | 9.04 |
| | | **No. of Peptides:** | 1 |

| 10 | 20 | 30 | 40 | 50 | 60 | 70 | 80 | 90 | 100 | 110 | 120 |
|---|---|---|---|---|---|---|---|---|---|---|---|
| DVVMTQTPLS | LPVSLGDQAS | ISCRSSQSLV | HSNGNTYLNW | YLQKAGQSPK | LLIYKVSNRF | SGVPDRFSGS | GSGTDFTLKI | SRVEAEDLGI | YFCSQTTHVP | PTFGGGTKLE | IKR |

| Cmpd. | m/z meas. | Δ m/z [ppm] | z | Rt [min] | Score | P | Range | Sequence | Modification |
|---|---|---|---|---|---|---|---|---|---|
| 564 | 652.3075 | -6.78 | 2 | 34.24 | 91.85 | 0 | 67-79 | R.FSGSGSGTDFTLK.I | |

**Protein 16:** Keratin, type II cytoskeletal 5 OS=Mus musculus GN=Krt5 PE=1 SV=1

| | | | |
|---|---|---|---|
| **Accession:** | K2C5_MOUSE | **Score:** | 86.58 |
| **Database:** | SwissProt | **MW [kDa]:** | 61.70 |
| **Seq. Coverage [%]:** | 2.10 % | **pI:** | 7.59 |
| | | **No. of Peptides:** | 1 |

| 10 | 20 | 30 | 40 | 50 | 60 | 70 | 80 | 90 | 100 | 110 | 120 |
|---|---|---|---|---|---|---|---|---|---|---|---|
| MSRQSSVSFR | SGGSRSFSAA | SAITPSVSRT | SFSSVSRSGG | GGGGRISLGG | ACGGAGGYGSR | SLYNVGGSKR | ISYSSGGGSF | RNQFGAGGFG | FGGGAGSGFG | FGGGAGSGFG | FGGGAGFGGG |
| 130 | 140 | 150 | 160 | 170 | 180 | 190 | 200 | 210 | 220 | 230 | 240 |
| YGGAGFPVCP | PGGIQEVTVN | QNLLTPLNLQ | IDPTIQRVRT | EEREQIKTLN | NKFASFIDKV | RFLEQQNKVL | DTKWALLQEQ | GTKTIKQNLD | PLFEQYINNL | RRQLDGVLGE | RGRLDSELRN |
| 250 | 260 | 270 | 280 | 290 | 300 | 310 | 320 | 330 | 340 | 350 | 360 |
| MQDLVEDYKN | KYEDEINKRT | TAENEFVMLK | KDVDAAYMNK | VELEARVDAL | MDEINFMKMF | FDAELSQMQT | HVSDTSVVLS | MDNNRSLDLD | SIIAEVKAQY | EDIANRSRTE | AESWYQTKYE |
| 370 | 380 | 390 | 400 | 410 | 420 | 430 | 440 | 450 | 460 | 470 | 480 |
| ELQQTAGRHG | DDLRNTKHEI | SEMNRMIQRL | RSEIDNVKKQ | CANLQNAIAE | AEQRGELALK | DARNKLTELE | EALQKAKQDM | ARLLREYQEL | MNTKLALDVE | IATYRKLLEG | EECRLSGEGV |
| 490 | 500 | 510 | 520 | 530 | 540 | 550 | 560 | 570 | 580 | 590 | |
| GPVNISVVTN | SVSSGYGGGS | SIGVGSGFGG | GLGSGFAGGL | GPRFTRGGGG | LGLGSGLSVG | GSGFSAGSSQ | GGMSFGSGGG | SGSSVKFVST | TSSSRRSFKS | | |

| Cmpd. | m/z meas. | Δ m/z [ppm] | z | Rt [min] | Score | P | Range | Sequence | Modification |
|---|---|---|---|---|---|---|---|---|---|
| 1322 | 651.8546 | -10.17 | 2 | 58.95 | 86.58 | 0 | 326-337 | R.SLDLDSIIAEVK.A | |

**Protein 17:** Keratin, type II cytoskeletal 79 OS=Mus musculus GN=Krt79 PE=1 SV=2

**Accession:** K2C79_MOUSE | **Score:** 61.62

**Database:** SwissProt | **MW [kDa]:** 57.50

**Seq. Coverage [%]:** 2.30 % | **pI:** 7.55

**No. of Peptides:** 1

| 10 | 20 | 30 | 40 | 50 | 60 | 70 | 80 | 90 | 100 | 110 | 120 |
|---|---|---|---|---|---|---|---|---|---|---|---|
| MRSSLSRQTF | STKGGFSSNS | ASGGGGSRMR | TSYSSVTMSR | GSGGGGGVRS | GSSSGGFGSR | SLYNLGGKNI | SVSMACGASS | GRALGGFGSG | AYVGLGASRQ | TFGPVCPPGG | IQEVTVNQSL |
| 130 | 140 | 150 | 160 | 170 | 180 | 190 | 200 | 210 | 220 | 230 | 240 |
| LTPLNVEIDP | EIQRVRTQER | EQIKTLNNKF | ASFIDKVRFL | EQQNKVLETK | WALLQEQSQN | TGVARSLEPF | FENYLSTLRR | QLDTKQSERG | RLDMELRNVQ | DNLEDFKNKY | EDEINKRTAL |
| 250 | 260 | 270 | 280 | 290 | 300 | 310 | 320 | 330 | 340 | 350 | 360 |
| ENEFVLLKKD | VDAAYMGRMD | LHGKVDSLTQ | EIDFLQQLFE | MELSQVQTNV | SDTNVILSMD | NNRNLDLDSI | IAEVKAQYEL | IAQKSRAEAE | SWYQTKYEEL | QVTAGKHGDS | LRDTKNEIAE |
| 370 | 380 | 390 | 400 | 410 | 420 | 430 | 440 | 450 | 460 | 470 | 480 |
| LTRTTQRLQG | EVDAAKKQCQ | QLQTAIAEAE | QNGEMALKDA | KKKLGDLDTA | LHQAKEDLAR | MLREYQDLVS | VKLALDMEIA | TYRKLLESEE | SRMSGDCPSA | ISISVTGNST | SVCAGGTAGF |
| 490 | 500 | 510 | 520 | 530 | 540 | | | | | | |
| GNGLSLGGAG | GASKGGFGSS | VSYGAAKGGQ | VSGGTSILRK | TTTVKTSSRR | Y | | | | | | |

| Cmpd. | m/z meas. | Δ m/z [ppm] | z | Rt [min] | Score | P | Range | Sequence | Modification |
|---|---|---|---|---|---|---|---|---|---|
| 1319 | 665.3596 | -10.60 | 2 | 58.83 | 61.62 | 0 | 304-315 | R.NLDLDSIIAEVK.A | |

**Protein 18:** Keratin, type I cytoskeletal 10 OS=Mus musculus GN=Krt10 PE=1 SV=3

**Accession:** K1C10_MOUSE | **Score:** 54.32

**Database:** SwissProt | **MW [kDa]:** 57.70

**Seq. Coverage [%]:** 2.10 % | **pI:** 5.04

**No. of Peptides:** 1

| 10 | 20 | 30 | 40 | 50 | 60 | 70 | 80 | 90 | 100 | 110 | 120 |
|---|---|---|---|---|---|---|---|---|---|---|---|
| MSVLYSSSSK | QFSSSRSGGG | GGGGSVRVSS | TRGSLGGGYS | SGGFSGGSFS | RGSSGGGCFG | GSSGGYGGFG | GGGSFGGGYG | GSSFGGGYGG | SSFGGGYGGS | SFGGAGFGGG | GSFGGGSFGG |
| 130 | 140 | 150 | 160 | 170 | 180 | 190 | 200 | 210 | 220 | 230 | 240 |
| GSYGGGFGGG | GFGGDGGSLL | SGNGRVTMQN | LNDRLASYMD | KVRALEESNY | ELEGKIKEWY | EKHGNSSQRE | PRDYSKYYKT | IEDLKGQILT | LTTDNANVLL | QIDNARLAAD | DFRLKYENEV |
| 250 | 260 | 270 | 280 | 290 | 300 | 310 | 320 | 330 | 340 | 350 | 360 |
| TLRQSVEADI | NGLRRVLDEL | TLSKSDLEMQ | IESLNEELAY | LKKNHEEEMR | DLQNVSTGDV | NVEMNAAPGV | DLTQLLNNMR | NQYEQLAEKN | RKDAEEWFNQ | KSKELTTEID | SNIEQMSSHK |
| 370 | 380 | 390 | 400 | 410 | 420 | 430 | 440 | 450 | 460 | 470 | 480 |
| SEITELRRTV | QGLEIELQSQ | LALKQSLEAS | LAETEGRYCV | QLSQIQSQIS | ALEEQLQQIR | AETECQNAEY | QQLLDIKTRL | ENEIQTYRSL | LEGEGSSSGG | GGGRRGGSGG | GSYGGSSGGG |
| 490 | 500 | 510 | 520 | 530 | 540 | 550 | 560 | 570 | 580 | | |
| SYGGSSGGGG | SYGGSSGGGG | SYGGGSSGGG | SHGGSSGGGY | GGGSSSGGAG | GHGGSSGGGY | GGGSSSGGQG | GSGGFKSSGG | GDQSSKGPRY | | | |

| Cmpd. | m/z meas. | Δ m/z [ppm] | z | Rt [min] | Score | P | Range | Sequence | Modification |
|---|---|---|---|---|---|---|---|---|---|
| 402 | 691.3228 | -7.08 | 2 | 29.19 | 56.00 | 0 | 164-175 | R.ALEESNYELEGK.I | |

**Protein 19:** Complement C4-B OS=Mus musculus GN=C4b PE=1 SV=3

**Accession:** CO4B_MOUSE

**Database:** SwissProt

**Seq. Coverage [%]:** 1.40 %

**Score:** 47.74

**MW [kDa]:** 192.80

**pI:** 7.38

**No. of Peptides:** 1

| | | | | | | | | | | | |
|---|---|---|---|---|---|---|---|---|---|---|---|
| 10 | 20 | 30 | 40 | 50 | 60 | 70 | 80 | 90 | 100 | 110 | 120 |
| MRLLWGLAWV | FSFCASSLQK | PRLLLFSPSV | VNLGTPLSVG | VQLLDAPPGQ | EVKGSVFLRN | PKGGSCSPKK | DFKLSSGDDF | VLLSLEVPLE | DVRSCGLFDL | RRAPHIQLVA | QSPWLRNTAF |
| 130 | 140 | 150 | 160 | 170 | 180 | 190 | 200 | 210 | 220 | 230 | 240 |
| KATETQGVNL | LFSSRRGHIF | VQTDQPIYNP | GQRVRYRVFA | LDQKMRPSTD | FLTITVENSH | GLRVLKKEIF | TSTSIFQDAF | TIPDISEPGT | WKISARFSDG | LESNRSTHFE | VKKYVLPNFE |
| 250 | 260 | 270 | 280 | 290 | 300 | 310 | 320 | 330 | 340 | 350 | 360 |
| VKITPWKPYI | LMVPSNSDEI | QLDIQARYIY | GKPVQGVAYT | RFALMDEQGK | RTFLRGLETQ | AKLVEGRTHI | SISKDQFQAA | LDKINIGVRD | LEGLRLYAAT | AVIESPGGEM | EEAELTSWRF |
| 370 | 380 | 390 | 400 | 410 | 420 | 430 | 440 | 450 | 460 | 470 | 480 |
| VSSAFSLDLS | RTKRHLVPGA | HFLLQALVQE | MSGSEASNVP | VKVSATLVSG | SDSQVLDIQQ | STNGIGQVSI | SFPIPPTVTE | LRLLVSAGSL | YPAIARLTVQ | APPSRGTGFL | SIEPLDPRSP |
| 490 | 500 | 510 | 520 | 530 | 540 | 550 | 560 | 570 | 580 | 590 | 600 |
| SVGDTFILNL | QPVGIPAPTF | SHYYYMIISR | GQIMAMGREP | RKTVTSVSVL | VDHQLAPSFY | FVAYFYHQGH | PVANSLLINI | QSRDCEGKLQ | LKVDGAKEYR | NADMMKLRIQ | TDSKALVALG |
| 610 | 620 | 630 | 640 | 650 | 660 | 670 | 680 | 690 | 700 | 710 | 720 |
| AVDMALYAVG | GRSHKPLDMS | KVFEVINSYN | VGCGPGGGDD | ALQVFQDAGL | AFSDGDRLTQ | TREDLSCPKE | KKSRQKRNVN | FQKAVSEKLG | QYSSPDAKRC | CQDGMTKLPM | KRTCEQRAAR |
| 730 | 740 | 750 | 760 | 770 | 780 | 790 | 800 | 810 | 820 | 830 | 840 |
| VPQQACREPF | LSCCKFAEDL | RRNQTRSQAH | LARNNHNMLQ | EEDLIDEDDI | LVRTSFPENW | LWRVEPVDSS | KLLTVWLPDS | MTTWEIHGVS | LSKSKGLCVA | KPTRVRVFRK | FHLHLRLPIS |
| 850 | 860 | 870 | 880 | 890 | 900 | 910 | 920 | 930 | 940 | 950 | 960 |
| IRRFEQFELR | PVLYNYLNDD | VAVSVHVTPV | EGLCLAGGGM | MAQQVTVPAG | SARPVAFSVV | PTAAANVPLK | VVARGVFDLG | DAVSKILQIE | KEGAIHREEL | VYNLDPLNNL | GRTLEIPGSS |
| 970 | 980 | 990 | 1000 | 1010 | 1020 | 1030 | 1040 | 1050 | 1060 | 1070 | 1080 |
| DPNIVPDGDF | SSLVRVTASE | PLETMGSEGA | LSPGGVASLL | RLPQGCAEQT | MIYLAPTLTA | SNYLDRTEQW | SKLSPETKDH | AVDLIQKGYM | RIQQFRKNDG | SFGAWLHRDS | STWLTAFVLK |
| 1090 | 1100 | 1110 | 1120 | 1130 | 1140 | 1150 | 1160 | 1170 | 1180 | 1190 | 1200 |
| ILSLAQEQVG | NSPEKLQETA | SWLLAQQLGD | GSFHDPCPVI | HRAMQGGLVG | SDETVALTAF | VVIALHHGLD | VFQDDDAKQL | KNRVEASITK | ANSFLGQKAS | AGLLGAHAAA | ITAYALTLTK |
| 1210 | 1220 | 1230 | 1240 | 1250 | 1260 | 1270 | 1280 | 1290 | 1300 | 1310 | 1320 |
| ASEDLRNVAH | NSLMAMAEET | GEHLYWGLVL | GSQDKVVLRP | TAPRSPTEPV | PQAPALWIET | TAYALLHLLL | REGKGKMADK | AASWLTHQGS | FHGAFRSTQD | TVVTLDALSA | YWIASHTTEE |
| 1330 | 1340 | 1350 | 1360 | 1370 | 1380 | 1390 | 1400 | 1410 | 1420 | 1430 | 1440 |
| KALNVTLSSM | GRNGLKTHGL | HLNNHQVKGL | EEELKFSLGS | TISVKVEGNS | KGTLKILRTY | NVLDMKNTTC | QDLQIEVKVT | GAVEYAWDAN | EDYEDYYDMP | AADDPSVPLQ | PVTPLQLFEG |
| 1450 | 1460 | 1470 | 1480 | 1490 | 1500 | 1510 | 1520 | 1530 | 1540 | 1550 | 1560 |
| RRSRRRREAP | KVVEEQESRV | QYTVCIWRNG | KLGLSGMAIA | DITLLSGFHA | LRADLEKLTS | LSDRYVSHFE | TDGPHVLLYF | DSVPTTRECV | GFGASQEVVV | GLVQPSSAVL | YDYYSPDHKC |
| 1570 | 1580 | 1590 | 1600 | 1610 | 1620 | 1630 | 1640 | 1650 | 1660 | 1670 | 1680 |
| SVFYAAPTKS | QLLATLCSGD | VCQCAEGKCP | RLLRSLERRV | EDKDGYRMRF | ACYYPRVEYG | FTVKVLREDG | RAAFRLFESK | ITQVLHFRKD | TMASIGQTRN | FLSRASCRLR | LEPNKEYLIM |
| 1690 | 1700 | 1710 | 1720 | 1730 | 1740 | | | | | | |
| GMDGETSDNK | GDPQYLLDSN | TWIEEMPSEQ | MCKSTRHRAA | CFQLKDFLME | FSSRGCQV | | | | | | |

| Cmpd. | m/z meas. | Δ m/z [ppm] | z | Rt [min] | Score | P | Range | Sequence | Modification |
|---|---|---|---|---|---|---|---|---|---|
| 1398 | 922.7743 | -13.95 | 3 | 61.61 | 47.74 | 0 | 1297-1321 | R.STQDTVVTLDALSAYWIASHTTEEK.A | |

**Protein 20:** Ig lambda-1 chain V region OS=Mus musculus PE=1 SV=2

| | | | |
|---|---|---|---|
| **Accession:** | LV1A_MOUSE | **Score:** | 37.63 |
| **Database:** | SwissProt | **MW [kDa]:** | 12.20 |
| **Seq. Coverage [%]:** | 7.70 % | **pI:** | 5.08 |
| | | **No. of Peptides:** | 1 |

```
        10         20         30         40         50         60         70         80         90        100        110        120
MAWISLILSL LALSSGGAIS QAVVTQESAL TTSPGETVTL TCRSSTGAVT TSNYANWVQE KPDHLFTGLI GGTNNRAPGV PARFSGSLIG DKAALTITGA QTEDEAIYFC ALWYSNH
```

| Cmpd. | m/z meas. | Δ m/z [ppm] | z | Rt [min] | Score | P | Range | Sequence | Modification |
|---|---|---|---|---|---|---|---|---|---|
| 445 | 462.2419 | -7.32 | 2 | 30.50 | 37.63 | 0 | 84-92 | R.FSGSLIGDK.A | |

**Protein 21:** Ig heavy chain V-III region A4 OS=Mus musculus PE=1 SV=1

| | | | |
|---|---|---|---|
| **Accession:** | HVM27_MOUSE | **Score:** | 37.24 |
| **Database:** | SwissProt | **MW [kDa]:** | 12.70 |
| **Seq. Coverage [%]:** | 8.00 % | **pI:** | 7.00 |
| | | **No. of Peptides:** | 1 |

```
        10         20         30         40         50         60         70         80         90        100        110        120
EVKLEESGGG LVQPGGSMKL SCVASGFTFS NYWMNWVRQS PEKGLEWVAE IRLKSHNYAT HYAESVKGRF TISRDDSKSS VYLQMNNLRA EDTGIYYCTT GFAYWGQGTL VTV
```

| Cmpd. | m/z meas. | Δ m/z [ppm] | z | Rt [min] | Score | P | Range | Sequence | Modification |
|---|---|---|---|---|---|---|---|---|---|
| 1027 | 536.7877 | -9.79 | 2 | 48.90 | 37.24 | 0 | 44-52 | K.GLEWVAEIR.L | |

| | | | |
|---|---|---|---|
| **Protein 22:** | Deubiquitinating protein VCIP135 OS=Mus musculus GN=Vcpip1 PE=1 SV=1 | | |
| **Accession:** | VCIP1_MOUSE | **Score:** | 28.67 |
| **Database:** | SwissProt | **MW [kDa]:** | 134.40 |
| **Seq. Coverage [%]:** | 1.20 % | **pI:** | 6.72 |
| | | **No. of Peptides:** | 1 |

| 10 | 20 | 30 | 40 | 50 | 60 | 70 | 80 | 90 | 100 | 110 | 120 |
|---|---|---|---|---|---|---|---|---|---|---|---|
| MSQPPPPPPL | PPPPPPPEAP | QTSSSLAAAA | SPGGLSKRRD | RRILSGSCPD | PKCQARLFFP | ASGSVSIECT | ECGQRHEQQQ | LLGVEEVTDP | DVVLHNLLRN | ALLGVTGAPK | KNTELVKVMG |
| 130 | 140 | 150 | 160 | 170 | 180 | 190 | 200 | 210 | 220 | 230 | 240 |
| LSNYHCKLLS | PILARYGMDK | QTGRAKLLRD | MNQGELFDCA | LLGDRAFLIE | PEHVNTVGYG | KDRSGSLLYL | HDTLEDIKRA | NKSQECLIPV | HVDGDGHCLV | HAVSRALVGR | ELFWHALREN |
| 250 | 260 | 270 | 280 | 290 | 300 | 310 | 320 | 330 | 340 | 350 | 360 |
| LKQHFQQHLA | RYQALFHDFI | DAAEWEDIIN | ECDPLFVPPE | GVPLGLRNIH | IFGLANVLHR | PIILLDSLSG | MRSSGDYSAT | FLPGLIPAEK | CTGRDGHLNK | PICIAWSSSG | RNHYIPLVGI |
| 370 | 380 | 390 | 400 | 410 | 420 | 430 | 440 | 450 | 460 | 470 | 480 |
| KGAALPKLPM | NLLPKAWGVP | QDLIKKYIKL | EEDGGCVIGG | DRSLQDKYLL | RLVAAMEEVF | MDKHGIHPSL | VADVHQYFYR | RTGVIGVQPE | EVTAAAKKAV | MDNRLHKCLL | YGALSELHVP |
| 490 | 500 | 510 | 520 | 530 | 540 | 550 | 560 | 570 | 580 | 590 | 600 |
| SEWLAPGGKL | YNLAKSTHGQ | LRPDKNYSFP | LNNLVCSYDP | VKDVLLPDYG | LSNLTACNWC | HGSSVRRVRG | DGSIVYLDGD | RTNSRSTGGK | CGCGFKHFWE | GKEYDNLPEA | FPITLEWGGR |
| 610 | 620 | 630 | 640 | 650 | 660 | 670 | 680 | 690 | 700 | 710 | 720 |
| VVRETVYWFQ | YESDPSLNSN | VYDVAMKLVT | KHFPGEFGSE | ILVQKVVHTI | LHQTAKKNPD | DYTPVNIDGA | HAQRVGDVQG | QELESQLPTK | IILTGQKTKT | LHKEELNMSK | TERTIQQNIT |
| 730 | 740 | 750 | 760 | 770 | 780 | 790 | 800 | 810 | 820 | 830 | 840 |
| EQASVMQKRK | TEKLKQEQKG | QPRTVSPSTI | RDGPSSAPAT | PTKAPYSPTT | SKEKKIRITT | NDGRQSMVTL | KPSTTFFELQ | ESIAREFNIP | PYLQCIRYGF | PPKELMPPQA | GMEKEPVPLQ |
| 850 | 860 | 870 | 880 | 890 | 900 | 910 | 920 | 930 | 940 | 950 | 960 |
| HGDRITIEIL | KGRAEGGPST | AAHSAHTVKQ | EEIAVTGKLS | SKELQEQADK | EMYSLCLLAT | LMGEDVWSYA | KGLPHMFQQG | GVFYNIMKKT | MGMADGKHCT | FPHLPGKTFV | YNASEDRLEL |
| 970 | 980 | 990 | 1000 | 1010 | 1020 | 1030 | 1040 | 1050 | 1060 | 1070 | 1080 |
| CVDAAGHFPI | GPDVEDLVKE | AVSQVRAEAT | TRSRESSPSH | GLLKLGSGGV | VKKKSEQLHN | VTAFQGKGHS | LGTASSHPHI | DPRARETLAV | RKHNTGTDFS | NSSIKTEPPV | FTAASSNSEL |
| 1090 | 1100 | 1110 | 1120 | 1130 | 1140 | 1150 | 1160 | 1170 | 1180 | 1190 | 1200 |
| IRIAPGVVTM | RDGRQIDPDV | VEAQRKKLQE | MVSSIQASMD | KHLRDQSAEQ | APSDLSQRKV | EVVSSVRPVN | LQTGLPEPFS | LTGGTENLNT | ETTDSHVADV | LGAAFATRSK | AQKENSMEEP |
| 1210 | 1220 | 1230 | | | | | | | | | |
| EEMDSQDAET | TNTTEPMDHS | | | | | | | | | | |

| Cmpd. | m/z meas. | Δ m/z [ppm] | z | Rt [min] | Score | P | Range | Sequence | Modification |
|---|---|---|---|---|---|---|---|---|---|
| 388 | 554.2862 | -17.20 | 3 | 28.80 | 28.67 | 2 | 980-994 | K.EAVSQVRAEATTRSR.E | |

**Protein 23:** Ig heavy chain V region B1-8/186-2 OS=Mus musculus GN=Ighv1-72 PE=1 SV=1

| | | | |
|---|---|---|---|
| **Accession:** | HVM07_MOUSE | **Score:** | 28.01 |
| **Database:** | SwissProt | **MW [kDa]:** | 15.40 |
| **Seq. Coverage [%]:** | 5.00 % | **pI:** | 8.94 |
| | | **No. of Peptides:** | 1 |

```
        10         20         30         40         50         60         70         80         90        100        110        120
MGWSCIMLFL AATATGVHSQ VQLQQPGAEL VKPGASVKLS CKASGYTFTS YWMHWVKQRP GRGLEWIGRI DPNSGGTKYN EKFKSKATLT VDKPSSTAYM QLSSLTSEDS AVYYCARYDY
       130        140
YGSSYFDYWG QGTTLTVSS
```

| Cmpd. | m/z meas. | Δ m/z [ppm] | z | Rt [min] | Score | P | Range | Sequence | Modification |
|---|---|---|---|---|---|---|---|---|---|
| 742 | 415.7250 | -10.96 | 2 | 39.61 | 28.01 | 0 | 63-69 | R.GLEWIGR.I | |

**Protein 24:** Complement C3 OS=Mus musculus GN=C3 PE=1 SV=3

| | | | |
|---|---|---|---|
| **Accession:** | CO3_MOUSE | **Score:** | 26.91 |
| **Database:** | SwissProt | **MW [kDa]:** | 186.40 |
| **Seq. Coverage [%]:** | 1.30 % | **pI:** | 6.29 |
| | | **No. of Peptides:** | 1 |

**Modification(s):** Carbamidomethyl

### Protein Report

| | | | | | | | | | | | |
|---|---|---|---|---|---|---|---|---|---|---|---|
| 10 | 20 | 30 | 40 | 50 | 60 | 70 | 80 | 90 | 100 | 110 | 120 |
| MGPASGSQLL | VLLLLLASSP | LALGIPMYSI | ITPNVLRLES | EETIVLEAHD | AQGDIPVTVT | VQDFLKRQVL | TSEKTVLTGA | SGHLRSVSIK | IPASKEFNSD | KEGHKYVTVV | ANFGETVVEK |
| 130 | 140 | 150 | 160 | 170 | 180 | 190 | 200 | 210 | 220 | 230 | 240 |
| AVMVSFQSGY | LFIQTDKTIY | TPGSTVLYRI | FTVDNNLLPV | GKTVVILIET | PDGIPVKRDI | LSSNNQHGIL | PLSWNIPELV | NMGQWKIRAF | YEHAPKQIFS | AEFEVKEYVL | PSFEVRVEPT |
| 250 | 260 | 270 | 280 | 290 | 300 | 310 | 320 | 330 | 340 | 350 | 360 |
| ETFYYIDDPN | GLEVSIIAKF | LYGKNVDGTA | FVIFGVQDGD | KKISLAHSLT | RVVIEDGVGD | AVLTRKVLME | GVRPSNADAL | VGKSLYVSVT | VILHSGSDMV | EAERSGIPIV | TSPYQIHFTK |
| 370 | 380 | 390 | 400 | 410 | 420 | 430 | 440 | 450 | 460 | 470 | 480 |
| TPKFFKPAMP | FDLMVFVTNP | DGSPASKVLV | VTQGSNAKAL | TQDDGVAKLS | INTPNSRQPL | TITVRTKKDT | LPESRQATKT | MEAHPYSTMH | NSNNYLHLSV | SRMELKPGDN | LNVNFHLRTD |
| 490 | 500 | 510 | 520 | 530 | 540 | 550 | 560 | 570 | 580 | 590 | 600 |
| PGHEAKIRYY | TYLVMNKGKL | LKAGRQVREP | GQDLVVLSLP | ITPEFIPSFR | LVAYYTLIGA | SGQREVVADS | VWVDVKDSCI | GTLVVKGDPR | DNHLAPGQQT | TLRIEGNQGA | RVGLVAVDKG |
| 610 | 620 | 630 | 640 | 650 | 660 | 670 | 680 | 690 | 700 | 710 | 720 |
| VFVLNKKNKL | TQSKIWDVVE | KADIGCTPGS | GKNYAGVFMD | AGLAFKTSQG | LQTEQRADLE | CTKPAARRRR | SVQLMERRMD | KAGQYTDKGL | RKCCEDGMRD | IPMRYSCQRR | ARLITQGENC |
| 730 | 740 | 750 | 760 | 770 | 780 | 790 | 800 | 810 | 820 | 830 | 840 |
| IKAFIDCCNH | ITKLREQHRR | DHVLGLARSE | LEEDIIPEED | IISRSHFPQS | WLWTIEELKE | PEKNGISTKV | MNIFLKDSIT | TWEILAVSLS | DKKGICVADP | YEIRVMQDFF | IDLRLPYSVV |
| 850 | 860 | 870 | 880 | 890 | 900 | 910 | 920 | 930 | 940 | 950 | 960 |
| RNEQVEIRAV | LFNYREQEEL | KVRVELLHNP | AFCSMATAKN | RYFQTIKIPP | KSSVAVPYVI | VPLKIGQQEV | EVKAAVFNHF | ISDGVKKTLK | VVPEGMRINK | TVAIHTLDPE | KLGQGGVQKV |
| 970 | 980 | 990 | 1000 | 1010 | 1020 | 1030 | 1040 | 1050 | 1060 | 1070 | 1080 |
| DVPAADLSDQ | VPDTDSETRI | ILQGSPVVQM | AEDAVDGERL | KHLIVTPAGC | GEQNMIGMTP | TVIAVHYLDQ | TEQWEKFGIE | KRQEALELIK | KGYTQQLAFK | QPSSAYAAFN | NRPPSTWLTA |
| 1090 | 1100 | 1110 | 1120 | 1130 | 1140 | 1150 | 1160 | 1170 | 1180 | 1190 | 1200 |
| YVVKVFSLAA | NLIAIDSHVL | CGAVKWLILE | KQKPDGVFQE | DGPVIHQEMI | GGFRNAKEAD | VSLTAFVLIA | LQEARDICEG | QVNSLPGSIN | KAGEYIEASY | MNLQRPYTVA | IAGYALALMN |
| 1210 | 1220 | 1230 | 1240 | 1250 | 1260 | 1270 | 1280 | 1290 | 1300 | 1310 | 1320 |
| KLEEPYLGKF | LNTAKDRNRW | EEPDQQLYNV | EATSYALLAL | LLLKDFDSVP | PVVRWLNEQR | YYGGGYGSTQ | ATFMVFQALA | QYQTDVPDHK | DLNMDVSFHL | PSRSSATTFR | LLWENGNLLR |
| 1330 | 1340 | 1350 | 1360 | 1370 | 1380 | 1390 | 1400 | 1410 | 1420 | 1430 | 1440 |
| SEETKQNEAF | SLTAKGKGRG | TLSVVAVYHA | KLKSKVTCKK | FDLRVSIRPA | PETAKKPEEA | KNTMFLEICT | KYLGDVDATM | SILDISMMTG | FAPDTKDLEL | LASGVDRYIS | KYEMNKAFSN |
| 1450 | 1460 | 1470 | 1480 | 1490 | 1500 | 1510 | 1520 | 1530 | 1540 | 1550 | 1560 |
| KNTLIIYLEK | ISHTEEDCLT | FKVHQYFNVG | LIQPGSVKVY | SYYNLEESCT | RFYHPEKDDG | MLSKLCHSEM | CRCAEENCFM | QQSQEKINLN | VRLDKACEPG | VDYVYKTELT | NIELLDDFDE |
| 1570 | 1580 | 1590 | 1600 | 1610 | 1620 | 1630 | 1640 | 1650 | 1660 | 1670 | |
| YTMTIQQVIK | SGSDEVQAGQ | QRKFISHIKC | RNALKLQKGK | KYLMWGLSSD | LWGEKPNTSY | IIGKDTWVEH | WPEAEECQDQ | KYQKQCEELG | AFTESMVVYG | CPN | |

| Cmpd. | m/z meas. | Δ m/z [ppm] | z | Rt [min] | Score | P | Range | Sequence | Modification |
|---|---|---|---|---|---|---|---|---|---|
| 1474 | 733.3948 | -13.73 | 3 | 64.36 | 26.91 | 0 | 1085-1105 | K.VFSLAANLIAIDSHVLCGAVK.W | Carbamidomethyl: 17 |

**Protein 25:** Ig kappa chain V-V region K2 (Fragment) OS=Mus musculus PE=1 SV=1

| | | | |
|---|---|---|---|
| **Accession:** | KV5A3_MOUSE | **Score:** | 23.99 |
| **Database:** | SwissProt | **MW [kDa]:** | 12.60 |
| **Seq. Coverage [%]:** | 8.70 % | **pI:** | 8.50 |
| | | **No. of Peptides:** | 1 |

```
        10         20         30         40         50         60         70         80         90        100        110        120
MSVLTQVLAL LLLWLTGARC DIQMTQSPAS LSASVGETVT ITCRASGNIH NYLAWYQQKQ GKSPQLLVYN AKTLADGVPS RFSGSGSGTQ YSLKINSLQP EDFGSYYCQH FWSTP
```

| Cmpd. | m/z meas. | Δ m/z [ppm] | z | Rt [min] | Score | P | Range | Sequence | Modification |
|---|---|---|---|---|---|---|---|---|---|
| 598 | 566.8166 | -9.03 | 2 | 35.24 | 23.99 | 0 | 63-72 | K.SPQLLVYNAK.T | |

**Protein 26:** Ig heavy chain V region RF OS=Mus musculus PE=1 SV=1

| | | | |
|---|---|---|---|
| **Accession:** | HVM53_MOUSE | **Score:** | 22.73 |
| **Database:** | SwissProt | **MW [kDa]:** | 12.90 |
| **Seq. Coverage [%]:** | 9.40 % | **pI:** | 9.38 |
| | | **No. of Peptides:** | 1 |

```
        10         20         30         40         50         60         70         80         90        100        110        120
MNFGLRLIFL VLVLKGVLCD VKLVESGGGL VKLGGSLKLS CAASGFTFSS YYMSWVRQTP EKRLELVAAI NSNGGSTYYP DTVKGRFTIS RDNAKNTLYL QMSSLKSEDT ALYYCAR
```

| Cmpd. | m/z meas. | Δ m/z [ppm] | z | Rt [min] | Score | P | Range | Sequence | Modification |
|---|---|---|---|---|---|---|---|---|---|
| 891 | 649.3376 | -10.82 | 2 | 44.33 | 22.73 | 0 | 96-106 | K.NTLYLQMSSLK.S | |

**Protein 27:** Ig kappa chain V19-17 OS=Mus musculus GN=Igk-V19-17 PE=1 SV=1

| | | | |
|---|---|---|---|
| **Accession:** | KV5A1_MOUSE | **Score:** | 22.30 |
| **Database:** | SwissProt | **MW [kDa]:** | 16.40 |
| **Seq. Coverage [%]:** | 6.00 % | **pI:** | 6.39 |
| | | **No. of Peptides:** | 1 |

| 10 | 20 | 30 | 40 | 50 | 60 | 70 | 80 | 90 | 100 | 110 | 120 |
|---|---|---|---|---|---|---|---|---|---|---|---|
| MHHTSMGIKM | ESQIQVFVFV | FLWLSGVDGD | IVMTQFAGVD | GDIVMTQSHK | FMSTSVGDRV | SITCKASQDV | STTVAWYQQK | PGQSPKLLIY | SASYRYTGVP | DRFTGSGSGT | DFTFTISSVQ |
| 130 | 140 | 150 | | | | | | | | | |
| AEDLAVYYCQ | QHYSTPPTFG | GGTKLEIKR | | | | | | | | | |

| Cmpd. | m/z meas. | Δ m/z [ppm] | z | Rt [min] | Score | P | Range | Sequence | Modification |
|---|---|---|---|---|---|---|---|---|---|
| 232 | 500.2279 | -7.84 | 2 | 24.12 | 22.30 | 0 | 51-59 | K.FMSTSVGDR.V | |

**Protein 28:** Tail-anchored protein insertion receptor WRB OS=Mus musculus GN=Wrb PE=1 SV=1

| | | | |
|---|---|---|---|
| **Accession:** | WRB_MOUSE | **Score:** | 20.68 |
| **Database:** | SwissProt | **MW [kDa]:** | 19.90 |
| **Seq. Coverage [%]:** | 5.20 % | **pI:** | 9.82 |
| | | **No. of Peptides:** | 1 |
| **Modification(s):** | Oxidation | | |

| 10 | 20 | 30 | 40 | 50 | 60 | 70 | 80 | 90 | 100 | 110 | 120 |
|---|---|---|---|---|---|---|---|---|---|---|---|
| MSASETDRWA | WLLVLSFVFG | CNLLRILLPS | LSSFISRVLQ | KDAEQESQMR | AEIQGMKQEL | STVNMMDEFA | RYARLERKIN | KMTDKLKTHV | KARTAQLAKI | KWFISVAFYI | LQAALMISLI |
| 130 | 140 | 150 | 160 | 170 | 180 | | | | | | |
| WKYYSVPVAV | VPSKWITPLD | RLVAFPTRVA | GGIGITCWIL | VCNKVVAIVL | HPFS | | | | | | |

| Cmpd. | m/z meas. | Δ m/z [ppm] | z | Rt [min] | Score | P | Range | Sequence | Modification |
|---|---|---|---|---|---|---|---|---|---|
| 730 | 555.2435 | 24.28 | 2 | 39.25 | 20.68 | 0 | 42-50 | K.DAEQESQMR.A | Oxidation: 8 |

| | | | |
|---|---|---|---|
| **Protein 29:** | Arf-GAP with SH3 domain, ANK repeat and PH domain-containing protein 2 OS=Mus musculus GN=Asap2 PE=1 SV=3 | | |
| **Accession:** | ASAP2_MOUSE | **Score:** | 19.85 |
| **Database:** | SwissProt | **MW [kDa]:** | 106.70 |
| **Seq. Coverage [%]:** | 0.80 % | **pI:** | 6.22 |
| | | **No. of Peptides:** | 1 |

| 10 | 20 | 30 | 40 | 50 | 60 | 70 | 80 | 90 | 100 | 110 | 120 |
|---|---|---|---|---|---|---|---|---|---|---|---|
| MPDQISVSEF | VAETHEDYKA | PTASSFTTRT | AQCRNTVAAI | EEALDVDRMV | LYKMKKSVKA | INISGLAHVE | NEEQYTQALE | KFGGNCVCRD | DPDLGSAFLK | FSVFTKELTA | LFKNLIQNMN |
| 130 | 140 | 150 | 160 | 170 | 180 | 190 | 200 | 210 | 220 | 230 | 240 |
| NIISFPLDSL | LKGDLKGVKG | DLKKPFDKAW | KDYETKITKI | EKEKKEHAKL | HGMIRTEISG | AEIAEEMEKE | RRFFQLQMCE | YLLKVNEIKV | KKGVDLLQNL | IKYFHAQCNF | FQDGLKAVES |
| 250 | 260 | 270 | 280 | 290 | 300 | 310 | 320 | 330 | 340 | 350 | 360 |
| LKPSIETLST | DLHTIKQAQD | EERRQLIQLR | DILKSALQVE | QKESRRDSQL | RQSTAYSLHQ | PQGNKEHGTE | RNGNLYKKSD | GIRKVWQKRK | CSVKNGFLTI | SHGTANRPPA | KLNLLTCQVK |
| 370 | 380 | 390 | 400 | 410 | 420 | 430 | 440 | 450 | 460 | 470 | 480 |
| TNPEEKKCFD | LISHDRTYHF | QAEDEQECQI | WMSVLQNSKE | EALNNAFKGD | DNTGENNIVQ | ELTKEIISEV | QRMTGNDVCC | DCGAPDPTWL | STNLGILTCI | ECSGIHRELG | VHYSRMQSLT |
| 490 | 500 | 510 | 520 | 530 | 540 | 550 | 560 | 570 | 580 | 590 | 600 |
| LDVLGTSELL | LAKNIGNAGF | NEIMECCLPS | EDPVKPNPGS | DMIARKDYIT | AKYMERRYAR | KKHADTAAKL | HSLCEAVKTR | DIFGLLQAYA | DGVDLTEKIP | LANGHEPDET | ALHLAVRSVD |
| 610 | 620 | 630 | 640 | 650 | 660 | 670 | 680 | 690 | 700 | 710 | 720 |
| RTSLHIVDFL | VQNSGNLDKQ | TGKGSTALHY | CCLTDNAECL | KLLLRGKASI | EIANESGETP | LDIAKRLKHE | HCEELLTQAL | SGRFNSHVHV | EYEWRLLHED | LDESDDDVDE | KLQPSPNRRE |
| 730 | 740 | 750 | 760 | 770 | 780 | 790 | 800 | 810 | 820 | 830 | 840 |
| DRPVSFYQLG | SSQFQSNAVS | LARDTANLTK | DKQRGFGPSI | LQNETYGAIL | SGSPPSSQSI | PPSTTSAPPL | PPRNVGKDPL | TTTPPPPVAK | TSGTLEAMNQ | PSKSSQPGTS | QSKPPPLPPQ |
| 850 | 860 | 870 | 880 | 890 | 900 | 910 | 920 | 930 | 940 | 950 | 960 |
| PPSRLPQKKP | ASGTDKPTPL | TNKGQPRGPE | ASGPLSNAMA | LQPPAPMPRK | SQATKSKPKR | VKALYNCVAD | NPDELTFSEG | DVIIVDGEED | QEWWIGHIDG | EPSRKGAFPV | SFVHFIAD |

| Cmpd. | m/z meas. | Δ m/z [ppm] | z | Rt [min] | Score | P | Range | Sequence | Modification |
|---|---|---|---|---|---|---|---|---|---|
| 491 | 487.2658 | -7.25 | 2 | 32.03 | 19.85 | 0 | 425-432 | K.EIISEVQR.M | |

| | | | |
|---|---|---|---|
| **Protein 30:** | STAM-binding protein OS=Mus musculus GN=Stambp PE=1 SV=1 | | |
| **Accession:** | STABP_MOUSE | **Score:** | 19.78 |
| **Database:** | SwissProt | **MW [kDa]:** | 48.50 |
| **Seq. Coverage [%]:** | 4.00 % | **pI:** | 6.20 |
| | | **No. of Peptides:** | 1 |

| 10 | 20 | 30 | 40 | 50 | 60 | 70 | 80 | 90 | 100 | 110 | 120 |
|---|---|---|---|---|---|---|---|---|---|---|---|
| MSDHGDVSLP | PQDRVRILSQ | LGSAVELNED | IPPRRYYRSG | VEIIRMASVY | SEEGNIEHAF | ILYNKYITLF | IEKLPKHRDY | KSAIIPEKKD | AVKKLKSVAF | PKAEELKTEL | LRRYTKEYEQ |
| 130 | 140 | 150 | 160 | 170 | 180 | 190 | 200 | 210 | 220 | 230 | 240 |
| YKERKKKEEE | ELARNIAIQQ | ELEKEKQRVA | QQKQKQLEQE | QFHAFEEMIQ | RQELEKERLK | IVQEFGKVDP | GPCGPLLPDL | EKPCVDVAPS | SPFSPTQTPD | CNTGMRPAKP | PVVDRSLKPG |
| 250 | 260 | 270 | 280 | 290 | 300 | 310 | 320 | 330 | 340 | 350 | 360 |
| ALSVIENVPT | IEGLRHIVVP | RNLCSEFLQL | ASANTAKGIE | TCGVLCGKLM | RNEFTITHVL | IPRQNGGPDY | CHTENEEEIF | FMQDDLGLLT | LGWIHTHPTQ | TAFLSSVDLH | THCSYQMMLP |
| 370 | 380 | 390 | 400 | 410 | 420 | 430 | | | | | |
| ESIAIVCSPK | FQETGFFKLT | DYGLQEISTC | RQKGFHPHGR | DPPLFCDCSH | VTVKDRIVTI | TDLR | | | | | |

| Cmpd. | m/z meas. | Δ m/z [ppm] | z | Rt [min] | Score | P | Range | Sequence | Modification |
|---|---|---|---|---|---|---|---|---|---|
| 686 | 681.3635 | 17.61 | 3 | 37.90 | 19.78 | 1 | 128-144 | K.EEEELARNIAIQQELEK.E | |

**Protein 31:** Prominin-1 OS=Mus musculus GN=Prom1 PE=1 SV=1

**Accession:** PROM1_MOUSE

**Database:** SwissProt

**Seq. Coverage [%]:** 1.20 %

**Score:** 17.65

**MW [kDa]:** 97.10

**pI:** 6.25

**No. of Peptides:** 1

**Modification(s):** Carbamidomethyl

| 10 | 20 | 30 | 40 | 50 | 60 | 70 | 80 | 90 | 100 | 110 | 120 |
|---|---|---|---|---|---|---|---|---|---|---|---|
| MALVFSALLL | LGLCGKISSE | GQPAFHNTPG | AMNYELPTTK | YETQDTFNAG | IVGPLYKMVH | IFLSVVQPND | FPLDLIKKLI | QNKKFDISVD | SKEPEIIVLA | LKIALYEIGV | LICAILGLLF |
| 130 | 140 | 150 | 160 | 170 | 180 | 190 | 200 | 210 | 220 | 230 | 240 |
| IILMPLVGCF | FCMCRCCNKC | GGEMHQRQKQ | NAPCRRKCLG | LSLLVICLLM | SLGIIYGFVA | NQQTRTRIKG | TQKLAKSNFR | DFQTLLTETP | KQIDYVVEQY | TNTKNKAFSD | LDGIGSVLGG |
| 250 | 260 | 270 | 280 | 290 | 300 | 310 | 320 | 330 | 340 | 350 | 360 |
| RIKDQLKPKV | TPVLEEIKAM | ATAIKQTKDA | LQNMSSSLKS | LQDAATQLNT | NLSSVRNSIE | NSLSSSDCTS | DPASKICDSI | RPSLSSLGSS | LNSSQLPSVD | RELNTVTEVD | KTDLESLVKR |
| 370 | 380 | 390 | 400 | 410 | 420 | 430 | 440 | 450 | 460 | 470 | 480 |
| GYTTIDEIPN | TIQNQTVDVI | KDVKNTLDSI | SSNIKDMSQS | IPIEDMLLQV | SHYLNNSNRY | LNQELPKLEE | YDSYWWLGGL | IVCFLLTLIV | TFFFLGLLCG | VFGYDKHATP | TRRGCVSNTG |
| 490 | 500 | 510 | 520 | 530 | 540 | 550 | 560 | 570 | 580 | 590 | 600 |
| GIFLMAGVGF | GFLFCWILMI | LVVLTFVVGA | NVEKLLCEPY | ENKKLLQVLD | TPYLLKEQWQ | FYLSGMLFNN | PDINMTFEQV | YRDCKRGRGI | YAAFQLENVV | NVSDHFNIDQ | ISENINTELE |
| 610 | 620 | 630 | 640 | 650 | 660 | 670 | 680 | 690 | 700 | 710 | 720 |
| NLNVNIDSIE | LLDNTGRKSL | EDFAHSGIDT | IDYSTYLKET | EKSPTEVNLL | TFASTLEAKA | NQLPEGKPKQ | AFLLDVQNIR | AIHQHLLPPV | QQSLNTLRQS | VWTLQQTSNK | LPEKVKKILA |
| 730 | 740 | 750 | 760 | 770 | 780 | 790 | 800 | 810 | 820 | 830 | 840 |
| SLDSVQHFLT | NNVSLIVIGE | TKKFGKTILG | YFEHYLHWVF | YAITEKMTSC | KPMATAMDSA | VNGILCGYVA | DPLNLFWFGI | GKATVLLLPA | VIIAIKLAKY | YRRMDSEDVY | DDVETVPMKN |
| 850 | 860 | 870 | | | | | | | | | |
| LEIGSNGYHK | DHLYGVHNPV | MTSPSRY | | | | | | | | | |

| Cmpd. | m/z meas. | Δ m/z [ppm] | z | Rt [min] | Score | P | Range | Sequence | Modification |
|---|---|---|---|---|---|---|---|---|---|
| 543 | 647.3263 | -4.22 | 2 | 33.60 | 17.65 | 1 | 515-524 | K.LLCEPYENKK.L | Carbamidomethyl: 3 |

| | | | |
|---|---|---|---|
| **Protein 32:** | Translation factor Guf1, mitochondrial OS=Mus musculus GN=Guf1 PE=1 SV=1 | | |
| **Accession:** | GUF1_MOUSE | **Score:** | 16.71 |
| **Database:** | SwissProt | **MW [kDa]:** | 72.40 |
| **Seq. Coverage [%]:** | 1.10 % | **pI:** | 8.87 |
| | | **No. of Peptides:** | 1 |

| 10 | 20 | 30 | 40 | 50 | 60 | 70 | 80 | 90 | 100 | 110 | 120 |
|---|---|---|---|---|---|---|---|---|---|---|---|
| MWALVGRALA | PWAAGARHAA | ASEPRAACRL | FSAAELKEKP | DMSRFPVEDI | RNFSIIAHVD | HGKSTLADRL | LELTGTIDKT | KKNKQVLDKL | QVERERGITV | KAQTASLFYS | FGGKQYLLNL |
| 130 | 140 | 150 | 160 | 170 | 180 | 190 | 200 | 210 | 220 | 230 | 240 |
| IDTPGHVDFS | YEVSRSLSAC | QGVLLVVDAN | EGIQAQTVAN | FFLAFEAQLS | VIPVINKIDL | KNADPERVGK | QIEKVFDIPS | EECIKISAKL | GTNVDSVLQA | VIERIPPPKV | HRENPLKALV |
| 250 | 260 | 270 | 280 | 290 | 300 | 310 | 320 | 330 | 340 | 350 | 360 |
| FDSTFDQYRG | VIANIALFDG | VVSKGDKIVS | AHTKKAYEVN | EVGILNPNEQ | PTHKLYAGQV | GFLIAGMKDV | TEAQIGDTLY | LHNHPVEPLP | GFKSAKPMVF | AGVYPIDQSE | YNNLKSAIEK |
| 370 | 380 | 390 | 400 | 410 | 420 | 430 | 440 | 450 | 460 | 470 | 480 |
| LTLNDSSVTV | HRDSSLALGA | GWRLGFLGLL | HMEVFNQRLE | QEYNASVILT | TPTVPYKAVL | SSAKLIKEYK | EKEITIINPA | QFPEKSQVTE | YLEPVVLGTV | ITPTEYTGKI | MALCQARRAI |
| 490 | 500 | 510 | 520 | 530 | 540 | 550 | 560 | 570 | 580 | 590 | 600 |
| QKNMTFIDEN | RVMLKYLFPL | NEIVVDFYDS | LKSLSSGYAS | FDYEDAGYQT | AELVKMDILL | NGNMVEELVT | VVHREKAYTV | GKSICERLKE | SLPRQLYEIA | IQAAVGSKVI | ARETVKAYRK |
| 610 | 620 | 630 | 640 | 650 | 660 | | | | | | |
| NVLAKCYGGD | ITRKMKLLKR | QSEGKKKLRK | IGNIEIPKDA | FIKVLKTQPN | K | | | | | | |

| Cmpd. | m/z meas. | Δ m/z [ppm] | z | Rt [min] | Score | P | Range | Sequence | Modification |
|---|---|---|---|---|---|---|---|---|---|
| 382 | 421.7567 | -3.89 | 2 | 28.67 | 16.71 | 1 | 568-574 | R.LKESLPR.Q | |

| | | | |
|---|---|---|---|
| **Protein 33:** | Putative Polycomb group protein ASXL3 OS=Mus musculus GN=Asxl3 PE=2 SV=3 | | |
| **Accession:** | ASXL3_MOUSE | **Score:** | 16.60 |
| **Database:** | SwissProt | **MW [kDa]:** | 243.50 |
| **Seq. Coverage [%]:** | 0.60 % | **pI:** | 5.77 |
| | | **No. of Peptides:** | 1 |

| | | | | | | | | | | | |
|---|---|---|---|---|---|---|---|---|---|---|---|
| 10 | 20 | 30 | 40 | 50 | 60 | 70 | 80 | 90 | 100 | 110 | 120 |
| MKDKRKKKDR | TWAEAARLAL | EKHPNSPMTA | KQILEVIQKE | GLKETGTSPL | ACLNAMLHTN | TRVGDGTFFK | IPGKSGLYAL | RKEESSCPVD | GTLDLVVDPD | LDGAEMAEAS | ANGEENRVCT |
| 130 | 140 | 150 | 160 | 170 | 180 | 190 | 200 | 210 | 220 | 230 | 240 |
| KQVTDEVSST | RDCSLTNTAV | QSKLVSSFQQ | HTKKALKQAL | RQQQKRRNGV | SMMVNKTVPR | VVLTPLKVSD | EQSDSPSGSE | SKNGEADSSD | KEMKHGQKSP | TGKQTSQHLK | RLKKSGLGHL |
| 250 | 260 | 270 | 280 | 290 | 300 | 310 | 320 | 330 | 340 | 350 | 360 |
| KWTKAEDIDI | ETPGSILVNT | NLRALINKHT | FASFPQHFQQ | YLLLLLPEVD | RQMGSDGILR | LSTSALNNEF | FAYAAQGWKQ | RLAEGEFTPE | MQLRIRQEIE | KEKKTEPWKE | KFFERFYGER |
| 370 | 380 | 390 | 400 | 410 | 420 | 430 | 440 | 450 | 460 | 470 | 480 |
| SGMSREESIK | LTSGPNHEGA | EGSSSHGDSG | IPGPSAQNAL | EEQQPKILKS | SASLEPDFCT | TVCPMLEVPV | KDVMTESETE | DIFIPEESVI | QEEVAEEVET | SIYECQDEHL | KTIPAFSEES |
| 490 | 500 | 510 | 520 | 530 | 540 | 550 | 560 | 570 | 580 | 590 | 600 |
| ESPATPCEEP | QVAAPEDSLE | SCVVMNDILH | TLPHIEVKIV | EKLECPQEEM | SVVIDQLEIC | DSLLPCPSSV | THILDVEQKE | QETTIETSAM | ALREGPSSLE | SQLPNEGIAV | DMELQSDPEE |
| 610 | 620 | 630 | 640 | 650 | 660 | 670 | 680 | 690 | 700 | 710 | 720 |
| QLSENACISE | TSFSSESPEG | ACASLPSPGG | ETQSTSEESC | TPASLETTFC | SEVSSTENTD | KYNQRNPTGE | SLHASLVSEV | SPLATSPEIS | EASLMSNLPL | TSEASPVSNL | PLTSEASPMS |
| 730 | 740 | 750 | 760 | 770 | 780 | 790 | 800 | 810 | 820 | 830 | 840 |
| DLPPTSETSS | ESSMPLTSET | PFVSSLPIPA | ETSPISNSSV | NERMVHQQRK | SPSGSEEANS | PQKEEPSIPT | KPLGESLVSH | PKPLSTIPEP | INMSSAMVPE | ALPPEGLHSQ | TLSQEPCNAH |
| 850 | 860 | 870 | 880 | 890 | 900 | 910 | 920 | 930 | 940 | 950 | 960 |
| VEMEKLYASS | IPELPSSEMT | KVKNHSVLQR | PEKKGLSAPL | EVPVFSEETE | TKGIELPPAK | LQDKQYAPSV | DKATFLEGSR | NKIHKQSSTL | NRLETSHTSK | VSEPSKSPDG | IRNDNRESEI |
| 970 | 980 | 990 | 1000 | 1010 | 1020 | 1030 | 1040 | 1050 | 1060 | 1070 | 1080 |
| SKRKTVEHSF | GICKEKRARI | EDDQSARSLA | SSSPPEKEQP | PREEPRVPPL | KIQLSKIGPP | FIIKSQPVSK | AESRASTSTS | VSSGRNTGAR | TLADIKARAQ | QARAQREAAA | AAAVAAAASI |
| 1090 | 1100 | 1110 | 1120 | 1130 | 1140 | 1150 | 1160 | 1170 | 1180 | 1190 | 1200 |
| VSGAMGSPGE | GGKARTLAHI | KEQTKAKLFA | KHQSRAHLFQ | TSKETRLPSV | STKEDSLNME | ASPTPETKME | GSTGVIIINP | NCRSPSSKPT | HIREITTVLQ | QPLNPPQIPE | TATDLSVHSS |
| 1210 | 1220 | 1230 | 1240 | 1250 | 1260 | 1270 | 1280 | 1290 | 1300 | 1310 | 1320 |
| DDNIPVSHLT | EKIVSSTSSE | NSSVPILHNK | SPINPIPMSV | CSTAMSGAIK | EHPFVSPVDK | SSVLMSVDSA | NSTISACNIS | MLKSIQGSDA | PCIALVPKCI | NRTPIPAAPE | GTGQSNSMDG |
| 1330 | 1340 | 1350 | 1360 | 1370 | 1380 | 1390 | 1400 | 1410 | 1420 | 1430 | 1440 |
| KALLVPSSKA | ANVISNQYTS | VPAPTIASNL | PNHLCTSSVL | IPPTGINNRF | VSEKIAMPGS | EEQAAVSIGA | TMRTALSCGD | SVAVTDSLVP | RSPIAMFAGN | MLTANSYNCP | PKLSGENLDN |
| 1450 | 1460 | 1470 | 1480 | 1490 | 1500 | 1510 | 1520 | 1530 | 1540 | 1550 | 1560 |
| NSGPLNRTDN | SEKPQQPAGG | FVPATINRSI | PCKVIVDHST | TLTSNLSLTV | SIESGDSSLD | SQTRSVRTDV | SIQPVACPQV | SVISRPEQAT | SEGLDHGSVF | IAAPTAKQDC | KTLQATCTSL |
| 1570 | 1580 | 1590 | 1600 | 1610 | 1620 | 1630 | 1640 | 1650 | 1660 | 1670 | 1680 |
| RELPLTLPDK | LNEVTVPTHG | FAEQARNSST | FKKETDTACS | NQYNPGNRIC | WSEDPMRNTA | PPVVSHSSSS | KQKEHPEQTG | LKAVKTEHVS | YAHVSDLHPR | NLITNVSLPV | KPEPHEVDKG |
| 1690 | 1700 | 1710 | 1720 | 1730 | 1740 | 1750 | 1760 | 1770 | 1780 | 1790 | 1800 |
| FRMDTEDFPG | PERPPPVTEV | TSSASVQPTQ | TMKPSTTSPV | EEAISLAPDT | LKRIPSASSS | SCRLSSVEAN | NPLVTQLLQG | NLPLEKVLPQ | PRLGAKLEIN | RLPLPLQTTS | VGKTGLERNM |
| 1810 | 1820 | 1830 | 1840 | 1850 | 1860 | 1870 | 1880 | 1890 | 1900 | 1910 | 1920 |
| VEMPSSSPNP | DGKGYLAGTL | APVQMRKREN | HPKKRAARTV | GDHAQVKCEP | GKMVMEPDVK | AVPCVISPSM | SQLGHNQPFK | QERLNKPSMA | NRIMPSPEVK | QQKRLLPACS | FQPSLFHVNK |
| 1930 | 1940 | 1950 | 1960 | 1970 | 1980 | 1990 | 2000 | 2010 | 2020 | 2030 | 2040 |
| NEGFHADTGT | SHRQQFYQMP | MAARGPLPTP | ALLQNSPKTP | VGCNAFAFNR | HLEQKALGDV | NLPTAPHQLR | LANMLSPNMP | IKEGEDGGGT | THTMPSKAVV | HAPLPPPPPP | PPPPPPPLAL |
| 2050 | 2060 | 2070 | 2080 | 2090 | 2100 | 2110 | 2120 | 2130 | 2140 | 2150 | 2160 |

| Cmpd. | m/z meas. | Δ m/z [ppm] | z | Rt [min] | Score | P | Range | Sequence | Modification |
|---|---|---|---|---|---|---|---|---|---|
| 764 | 688.8860 | 21.17 | 2 | 40.27 | 16.60 | 0 | 1814-1826 | K.GYLAGTLAPVQMR.K | |

| | | | |
|---|---|---|---|
| **Protein 34:** | Protein FAM208B OS=Mus musculus GN=Fam208b PE=1 SV=2 | | |
| **Accession:** | F208B_MOUSE | **Score:** | 16.36 |
| **Database:** | SwissProt | **MW [kDa]:** | 264.10 |
| **Seq. Coverage [%]:** | 0.30 % | **pI:** | 6.02 |
| | | **No. of Peptides:** | 1 |

| | | | | | | | | | | | |
|---|---|---|---|---|---|---|---|---|---|---|---|
| 10 | 20 | 30 | 40 | 50 | 60 | 70 | 80 | 90 | 100 | 110 | 120 |
| MAAPTSKIIL | ELNKNALISP | WKGQFIIQGC | LLCDITLWST | YGTVVPLQLP | RELDFKYVMD | VSSLKESLPE | AAFRRRSYLE | QKVCCQDLCF | DLYEVELTNK | QGENIDKLME | YVKNKELALI |
| 130 | 140 | 150 | 160 | 170 | 180 | 190 | 200 | 210 | 220 | 230 | 240 |
| KCLEDKSFFI | LFTSSALTPE | PGFGAEQMGL | HGLHLFHAPQ | TAGAKDLKVE | DGISLKVIPI | LPALSYALLE | AKKSLSEEGI | PPNILVKHSF | QELYKVDKSL | SLMAPPQDGV | EDTASTGKLS |
| 250 | 260 | 270 | 280 | 290 | 300 | 310 | 320 | 330 | 340 | 350 | 360 |
| HAFDLPPPLE | TCPSESLTHL | KCYFSDPAGY | TLDLSAALDL | LAEHPQFPCI | ADGVCDAGFS | LVMTPDPEFL | DSEMEIRKTE | TAEKSGRKLK | VKKKAVTPSS | NQRVQPKRKA | STTAVTLPSK |
| 370 | 380 | 390 | 400 | 410 | 420 | 430 | 440 | 450 | 460 | 470 | 480 |
| RVSLGRPTSK | RTVPRTDNRS | CNPTLKLVKG | QFPQKRKRGA | EVLAAQIVQK | TRLERKKQEA | SVSKDAPVPT | NTKRAKKQEK | SPGRIASQPK | PPMKKSPQKR | KVNVARGRRN | TRKQLQPAEK |
| 490 | 500 | 510 | 520 | 530 | 540 | 550 | 560 | 570 | 580 | 590 | 600 |
| EIALHLQSEI | SSDGQKDGLN | LSTSQQESIS | MIPKGPPENS | VISCDSQALN | MLADLALSSA | AASIPSCKPR | NLPCVSDLPR | NNVLLTKENP | LLGASDHEYH | KGVKSQKAVL | LPKPYSDEKI |
| 610 | 620 | 630 | 640 | 650 | 660 | 670 | 680 | 690 | 700 | 710 | 720 |
| SSESDLTRSQ | EENLVPCAQP | LPIAQPAPHS | EARELSDASQ | NSVVVEHSYA | LLLAEQSQKH | LHQRKLPSPA | FVKNGIKGPE | AGTPVGKVMP | FRHLQNTSPL | QKHSEDSLMK | HKSLFVSSTL |
| 730 | 740 | 750 | 760 | 770 | 780 | 790 | 800 | 810 | 820 | 830 | 840 |
| KEFFCSHTVL | KCDGSFKITF | KCEGEYVFSL | DSKYTSNPLE | KTVLRALHGP | WNTDLPENVD | DVKLLLHIWV | ALFYSKNNKV | IGSPRKVVEH | KNPAKYVCIN | RSLESLDHSE | IEAFSNVERP |
| 850 | 860 | 870 | 880 | 890 | 900 | 910 | 920 | 930 | 940 | 950 | 960 |
| SVEGSVDPLL | ETKETHIGHA | TNMTFPGPNR | VLPFINPPTT | RDLELCVQND | QKEVFTGECH | LDTSGNQNFI | YSCNTEVTGG | KTKQELSNKL | ETSNVVLSGV | VSAQSHGTCI | PSEDKTCQST |
| 970 | 980 | 990 | 1000 | 1010 | 1020 | 1030 | 1040 | 1050 | 1060 | 1070 | 1080 |
| KMVSYNDSVT | QATLTTAYDE | ASSELMCQKS | VFDNLENKVD | SFHPSPLIKT | DAVQDVIQHS | SHINNECQPS | VEKREDNVEC | MMVNLDPVTL | AFEKNASVPI | HTEVHTTDKP | TGFNIELVKR |
| 1090 | 1100 | 1110 | 1120 | 1130 | 1140 | 1150 | 1160 | 1170 | 1180 | 1190 | 1200 |
| VSPASSVQYP | MSALEEVQTQ | SSRDVPSLAM | SEYKDSKCLS | ASSVKKETPP | ESLCLLQKEI | PPLTSSADEG | LIMEALSLVK | SSSYSLASDK | TKCPQDDSLQ | TQNGLSMSLD | EVLEPSKVNV |
| 1210 | 1220 | 1230 | 1240 | 1250 | 1260 | 1270 | 1280 | 1290 | 1300 | 1310 | 1320 |
| VSSTSVTLRE | QPSPNCIPAM | SDVAGASTVI | MNSGSSSLNQ | EKILQTFSLV | FPKQTDLSLK | REEVSMELSG | EEADINLTLT | ISPPTSPSEE | IAAGEIEQFQ | KTPVSNVGQQ | SRSEEMVEPE |
| 1330 | 1340 | 1350 | 1360 | 1370 | 1380 | 1390 | 1400 | 1410 | 1420 | 1430 | 1440 |
| EEERLIRNRE | INSASCMSVY | PVESRELLKN | CTPEVTEKVN | VTLDFPFGPL | IEVSPASSPD | PIVQPGDRPS | SPCCLKLHSS | QSEKTNKFSQ | IKSGEVTIPE | KESLFLGPES | PKGQDKLAEV |
| 1450 | 1460 | 1470 | 1480 | 1490 | 1500 | 1510 | 1520 | 1530 | 1540 | 1550 | 1560 |
| QVQISAEMLQ | ITTNAEVEGR | VNMPGKVTKV | SVPSEHSENL | SFLEKVQCNT | ELNELSLPAK | YGGNFKPLEK | SGNSLEAGCM | ENRNVDVKHL | ALESSVPSCS | PRKVVENKSL | TDTLVSITTS |
| 1570 | 1580 | 1590 | 1600 | 1610 | 1620 | 1630 | 1640 | 1650 | 1660 | 1670 | 1680 |
| GIVNMSLKQT | SSKNIKKNVC | DSDVKTDSDV | KTEADSNMQT | EAVNSALIDK | TDVQAYSHPE | VSKFVSSSDS | AQCTYHAKPV | SVEPGFQTQE | IPVVRMASLL | KNIGVELHEE | KMDLSATGLQ |
| 1690 | 1700 | 1710 | 1720 | 1730 | 1740 | 1750 | 1760 | 1770 | 1780 | 1790 | 1800 |
| SNSMSAKDEQ | KTMHVLQDTI | CEVKEFLNGD | VFSQNAHSCQ | NTVDFSKSIS | EEPSASFVPE | FVDAICGVYK | EHTFNESPNM | VHETKADAET | LSRNTEISVN | ASMFCGPRSG | AYVQDSHDCK |
| 1810 | 1820 | 1830 | 1840 | 1850 | 1860 | 1870 | 1880 | 1890 | 1900 | 1910 | 1920 |
| SCKFDVENLR | GNHESQKDAV | KDSCDSFTSL | NNSDDTWACS | SKISTLETHI | PPRDQETEPR | LISPNKCIPR | YIQIPDSHGI | PKTYANFTIT | KEFKDTTRRL | HSLKRHRNLS | ANCNLLSSWT |
| 1930 | 1940 | 1950 | 1960 | 1970 | 1980 | 1990 | 2000 | 2010 | 2020 | 2030 | 2040 |
| STWHVTDDLT | QHTLDLEYLR | FDHKLKQIKK | GGSQQSSFPK | ESLVQISSGT | SPSTQTSEAS | GLHLPPESRS | PILVTVVRAD | TRQQSHHRRG | CSPSSLDGSS | SFWKKKCSQS | RNLKNSERSQ |
| 2050 | 2060 | 2070 | 2080 | 2090 | 2100 | 2110 | 2120 | 2130 | 2140 | 2150 | 2160 |

#### Protein Report

| Cmpd. | m/z meas. | Δ m/z [ppm] | z | Rt [min] | Score | P | Range | Sequence | Modification |
|---|---|---|---|---|---|---|---|---|---|
| 554 | 464.2703 | -28.28 | 2 | 33.87 | 16.36 | 2 | 462-469 | K.VNVARGRR.N | |

**Protein 35:** Centrosomal protein of 131 kDa OS=Mus musculus GN=Cep131 PE=1 SV=2

**Accession:** CP131_MOUSE

**Score:** 15.87

**Database:** SwissProt

**MW [kDa]:** 120.20

**Seq. Coverage [%]:** 1.10 %

**pI:** 8.88

**No. of Peptides:** 1

**Modification(s):** Oxidation

| | | | | | | | | | | | |
|---|---|---|---|---|---|---|---|---|---|---|---|
| 10 | 20 | 30 | 40 | 50 | 60 | 70 | 80 | 90 | 100 | 110 | 120 |
| MKGSRTITAT | PEGSPEAVDL | SLIGLPPPMS | QRPGSASATR | SIFRSMSVAT | GSEPRKKALE | ATGPGGPRAI | NNLRRSNSTT | QVNQSWTGSP | RPAEPTDFLM | LFEGSTSGRR | RVASLSKASS |
| 130 | 140 | 150 | 160 | 170 | 180 | 190 | 200 | 210 | 220 | 230 | 240 |
| EKEATWNVLD | EQPRGLALPA | SAQSPSTLDS | ALGPRRKECP | LAPSFTANNR | SNKGAVGNCV | TTMVHNHYAS | SKMVSPPKSS | NQTAPSLNNI | VKAAAREGGE | GSDLGKPRKN | LSSASQSARG |
| 250 | 260 | 270 | 280 | 290 | 300 | 310 | 320 | 330 | 340 | 350 | 360 |
| TTGLLRRREV | TEEEAERFIH | QVNQAAVTIQ | RWYRCQVQRR | RAGAAALEHL | LASKREGQRQ | RLGGGNLLEL | HRQEEAARKK | AREEKARQAR | QAAIQVLQQK | RAQKASEAEH | RRPKDRPETR |
| 370 | 380 | 390 | 400 | 410 | 420 | 430 | 440 | 450 | 460 | 470 | 480 |
| APEQPRPMQE | PGCVTHPKAN | NAGASIYPTG | PADPCPPASE | SSPEQWQSPE | DKPQDIHSQG | EARQDLAVSG | SSRGKARARA | TLDDLLDTLK | LLEEEPEPLP | HPKAYHKDRY | AWTDEEEDAN |
| 490 | 500 | 510 | 520 | 530 | 540 | 550 | 560 | 570 | 580 | 590 | 600 |
| SLTADNLEKF | GKLSAAPGPP | DDGTLLSEAK | LQSIMTFLDE | MEKSGQERPA | PWRESLVLEA | GSGSEGSTSV | MRLKLELEEK | KQAMALLQRA | LAQQRDLTVR | RVKETEKELT | RQLRQQKEQY |
| 610 | 620 | 630 | 640 | 650 | 660 | 670 | 680 | 690 | 700 | 710 | 720 |
| EATIQRHLSF | IDQLIEDKKV | LSEKCEAVVA | ELKHGDQRCR | ERVAQMQEQH | ELEIKKLKEL | MSATEKIRRE | KWINEKTKKI | KEITVRGLEP | EIQKLIAKHK | QEVRRLRGLH | EAELQQREEQ |
| 730 | 740 | 750 | 760 | 770 | 780 | 790 | 800 | 810 | 820 | 830 | 840 |
| AAQRHLRQAE | ELRQHLDRER | EVLGQQERER | AQQRFEQHLE | QEQRALEQQR | RRLYNEVAEE | KERLGQQAAR | QRAELEELRQ | QLEESSAALT | RALRAEFERS | REEQERRHQM | ELKALKDQLE |
| 850 | 860 | 870 | 880 | 890 | 900 | 910 | 920 | 930 | 940 | 950 | 960 |
| AERQAWVASC | AKKEEAWLLT | RERELKEEIR | KGRDQEIELV | IHRLEADMTL | AKEESERAAE | SRVKRVRDKY | ETELSELEQS | ERKLQERCSE | LKGRLGEAEG | EKERLQSLVR | QKEKELEDLR |
| 970 | 980 | 990 | 1000 | 1010 | 1020 | 1030 | 1040 | 1050 | 1060 | 1070 | |
| AVNTQMCSER | ASLAQVVRQE | FAEQLAASQE | ETQRVKVELA | ELQARQQVEL | DEVHRRVKTA | LARKEAAVNS | LRKQHEAAVK | RADHLEELLE | QHKGSSLSSK | | |

| Cmpd. | m/z meas. | Δ m/z [ppm] | z | Rt [min] | Score | P | Range | Sequence | Modification |
|---|---|---|---|---|---|---|---|---|---|
| 360 | 633.3236 | 19.39 | 2 | 27.97 | 15.87 | 1 | 45-56 | R.SMSVATGSEPRK.K | Oxidation: 2 |

**Protein 36:** Taste receptor type 2 member 7 OS=Mus musculus GN=Tas2r7 PE=2 SV=1

**Accession:** TA2R7_MOUSE

**Database:** SwissProt

**Seq. Coverage [%]:** 1.90 %

**Score:** 15.33

**MW [kDa]:** 35.50

**pI:** 9.58

**No. of Peptides:** 1

**Modification(s):** Oxidation

| 10 | 20 | 30 | 40 | 50 | 60 | 70 | 80 | 90 | 100 | 110 | 120 |
|---|---|---|---|---|---|---|---|---|---|---|---|
| MTYETDTTLM | LVAVGEALVG | ILGNAFIALV | NFMGWMKNRK | IASIDLILSS | VAMSRICLQC | IILLDCIILV | QYPDTYNRGK | EMRTVDFFWT | LTNHLSVWFA | TCLSIFYLFK | IANFFHPLFL |
| 130 | 140 | 150 | 160 | 170 | 180 | 190 | 200 | 210 | 220 | 230 | 240 |
| WIKWRIDKLI | LRTLLACVII | SLCFSLPVTE | NLSDDFRRCV | KTKERINSTL | RCKVNKAGHA | SVKVNLNLVM | LFPFSVSLVS | FLLLILSLWR | HTRQIQLSVT | GYKDPSTTAH | VKAMKAVISF |
| 250 | 260 | 270 | 280 | 290 | 300 | 310 | 320 | | | | |
| LALFVVYCLA | FLIATSSYFM | PESELAVIWG | ELIALIYPSS | HSFILILGSS | KLKQASVRVL | CRVKTMLKGK | KY | | | | |

| Cmpd. | m/z meas. | Δ m/z [ppm] | z | Rt [min] | Score | P | Range | Sequence | Modification |
|---|---|---|---|---|---|---|---|---|---|
| 692 | 368.2208 | -12.33 | 2 | 38.05 | 15.33 | 1 | 303-308 | R.VKTMLK.G | Oxidation: 4 |

### Table S1. Raw data of MS analysis for mouse BiEVs (TS)

#### Protein Report

BRUKER

##### Project Info

| | | | |
|---|---|---|---|
| **Name:** | NAWA | **Date:** | October 6, 2015 |

##### Sample Info & Protocols

| | | | |
|---|---|---|---|
| | | **Date:** | March 1, 2018 |
| **Name:** | 2018-3 | | |

##### Search Result Info

| Search Result | Location | Search Engine | Database | Ident. Compounds |
|---|---|---|---|---|
| maXis_mouse-2016_Mascot_2018-03-02 16:26:36 | /NAWA/2018-3/0302-kotani-S_Tray01-E2_01_6613.mgf | Mascot, 2.6.0 | SwissProt, SwissProt_2017_07.fasta | 56/1507 (FDR:4.05%) |

| | | | |
|---|---|---|---|
| **Protein 1:** | Ig mu chain C region OS=Mus musculus GN=Ighm PE=1 SV=2 | | |
| **Accession:** | IGHM_MOUSE | **Score:** | 671.06 |
| **Database:** | SwissProt | **MW [kDa]:** | 49.90 |
| **Seq. Coverage [%]:** | 31.30 % | **pI:** | 6.56 |
| | | **No. of Peptides:** | 10 |
| **Modification(s):** | Carbamidomethyl | | |

| 10 | 20 | 30 | 40 | 50 | 60 | 70 | 80 | 90 | 100 | 110 | 120 |
|---|---|---|---|---|---|---|---|---|---|---|---|
| SQSFPNVFPL | VSCESPLSDK | NLVAMGCLAR | DFLPSTISFT | WNYQNNTEVI | QGIRTFPTLR | TGGKYLATSQ | VLLSPKSILE | GSDEYLVCKI | HYGGKNRDLH | VPIPAVAEMN | PNVNVFVPPR |
| 130 | 140 | 150 | 160 | 170 | 180 | 190 | 200 | 210 | 220 | 230 | 240 |
| DGFSGPAPRK | SKLICEATNF | TPKPITVSWL | KDGKLVESGF | TTDPVTIENK | GSTPQTYKVI | STLTISEIDW | LNLNVYTCRV | DHRGLTFLKN | VSSTCAASPS | TDILTFTIPP | SFADIFLSKS |
| 250 | 260 | 270 | 280 | 290 | 300 | 310 | 320 | 330 | 340 | 350 | 360 |
| ANLTCLVSNL | ATYETLNISW | ASQSGEPLET | KIKIMESHPN | GTFSAKGVAS | VCVEDWNNRK | EFVCTVTHRD | LPSPQKKFIS | KPNEVHKHPP | AVYLLPPARE | QLNLRESATV | TCLVKGFSPA |
| 370 | 380 | 390 | 400 | 410 | 420 | 430 | 440 | 450 | 460 | | |
| DISVQWLQRG | QLLPQEKYVT | SAPMPEPGAP | GFYFTHSILT | VTEEEWNSGE | TYTCVVGHEA | LPHLVTERTV | DKSTGKPTLY | NVSLIMSDTG | GTCY | | |

| Cmpd. | m/z meas. | Δ m/z [ppm] | z | Rt [min] | Score | P | Range | Sequence | Modification |
|---|---|---|---|---|---|---|---|---|---|
| 700 | 552.7776 | -15.70 | 2 | 37.93 | 42.99 | 0 | 21-30 | K.NLVAMGCLAR.D | Carbamidomethyl: 7 |
| 744 | 660.3715 | -16.07 | 2 | 39.24 | 89.94 | 0 | 65-76 | K.YLATSQVLLSPK.S | |
| 806 | 756.8532 | -17.08 | 2 | 41.28 | 82.32 | 0 | 77-89 | K.SILEGSDEYLVCK.I | Carbamidomethyl: 12 |

| Cmpd. | m/z meas. | Δ m/z [ppm] | z | Rt [min] | Score | P | Range | Sequence | Modification |
|---|---|---|---|---|---|---|---|---|---|
| 1322 | 842.4299 | -23.04 | 3 | 58.42 | 61.37 | 0 | 98-120 | R.DLHVPIPAVAEMNPNVNVFVPPR.D | |
| 1159 | 740.0537 | -21.39 | 3 | 52.79 | 31.56 | 0 | 133-151 | K.LICEATNFTPKPITVSWLK.D | Carbamidomethyl: 3 |
| 771 | 875.4331 | -18.10 | 2 | 40.10 | 136.10 | 0 | 155-170 | K.LVESGFTTDPVTIENK.G | |
| 659 | 753.3327 | -14.62 | 2 | 36.57 | 66.16 | 0 | 287-299 | K.GVASVCVEDWNNR.K | Carbamidomethyl: 6 |
| 669 | 444.2524 | -15.22 | 3 | 36.93 | 29.73 | 0 | 328-339 | K.HPPAVYLLPPAR.E | |
| 393 | 554.2809 | -15.33 | 2 | 28.62 | 44.68 | 0 | 346-355 | R.ESATVTCLVK.G | Carbamidomethyl: 7 |
| 1266 | 802.3961 | -23.56 | 2 | 56.50 | 86.21 | 0 | 356-369 | K.GFSPADISVQWLQR.G | |

| | | | |
|---|---|---|---|
| **Protein 2:** | Actin, cytoplasmic 1 OS=Mus musculus GN=Actb PE=1 SV=1 | | |
| **Accession:** | ACTB_MOUSE | **Score:** | 153.82 |
| **Database:** | SwissProt | **MW [kDa]:** | 41.70 |
| **Seq. Coverage [%]:** | 9.10 % | **pI:** | 5.29 |
| | | **No. of Peptides:** | 2 |

| 10 | 20 | 30 | 40 | 50 | 60 | 70 | 80 | 90 | 100 | 110 | 120 |
|---|---|---|---|---|---|---|---|---|---|---|---|
| MDDDIAALVV | DNGSGMCKAG | FAGDDAPRAV | FPSIVGRPRH | QGVMVGMGQK | DSYVGDEAQS | KRGILTLKYP | IEHGIVTNWD | DMEKIWHHTF | YNELRVAPEE | HPVLLTEAPL | NPKANREKMT |
| 130 | 140 | 150 | 160 | 170 | 180 | 190 | 200 | 210 | 220 | 230 | 240 |
| QIMFETFNTP | AMYVAIQAVL | SLYASGRTTG | IVMDSGDGVT | HTVPIYEGYA | LPHAILRLDL | AGRDLTDYLM | KILTERGYSF | TTTAEREIVR | DIKEKLCYVA | LDFEQEMATA | ASSSSLEKSY |
| 250 | 260 | 270 | 280 | 290 | 300 | 310 | 320 | 330 | 340 | 350 | 360 |
| ELPDGQVITI | GNERFRCPEA | LFQPSFLGME | SCGIHETTFN | SIMKCDVDIR | KDLYANTVLS | GGTTMYPGIA | DRMQKEITAL | APSTMKIKII | APPERKYSVW | IGGSILASLS | TFQQMWISKQ |
| 370 | 380 | | | | | | | | | | |
| EYDESGPSIV | HRKCF | | | | | | | | | | |

| Cmpd. | m/z meas. | Δ m/z [ppm] | z | Rt [min] | Score | P | Range | Sequence | Modification |
|---|---|---|---|---|---|---|---|---|---|
| 766 | 652.0144 | -18.36 | 3 | 39.95 | 61.06 | 0 | 96-113 | R.VAPEEHPVLLTEAPLNPK.A | |
| 1015 | 895.9289 | -23.09 | 2 | 48.00 | 92.76 | 0 | 239-254 | K.SYELPDGQVITIGNER.F | |

**Protein 3:** Pregnancy zone protein OS=Mus musculus GN=Pzp PE=1 SV=3

| | | | |
|---|---|---|---|
| **Accession:** | PZP_MOUSE | **Score:** | 87.89 |
| **Database:** | SwissProt | **MW [kDa]:** | 165.70 |
| **Seq. Coverage [%]:** | 2.10 % | **pI:** | 6.24 |
| | | **No. of Peptides:** | 2 |

**Modification(s):** Carbamidomethyl

| | | | | | | | | | | | |
|---|---|---|---|---|---|---|---|---|---|---|---|
| 10 | 20 | 30 | 40 | 50 | 60 | 70 | 80 | 90 | 100 | 110 | 120 |
| MRRNQLPTPA | FLLLFLLLPR | DATTATAKPQ | YVVLVPSEVY | SGVPEKACVS | LNHVNETVML | SLTLEYAMQQ | TKLLTDQAVD | KDSFYCSPFT | ISGSPLPYTF | ITVEIKGPTQ | RFIKKKSIQI |
| 130 | 140 | 150 | 160 | 170 | 180 | 190 | 200 | 210 | 220 | 230 | 240 |
| IKAESPVFVQ | TDKPIYKPGQ | IVKFRVVSVD | ISFRPLNETF | PVVYIETPKR | NRIFQWQNIH | LAGGLHQLSF | PLSVEPALGI | YKVVVQKDSG | KKIEHSFEVK | EYVLPKFEVI | IKMQKTMAFL |
| 250 | 260 | 270 | 280 | 290 | 300 | 310 | 320 | 330 | 340 | 350 | 360 |
| EEELPITACG | VYTYGKPVPG | LVTLRVCRKY | SRYRSTCHNQ | NSMSICEEFS | QQADDKGCFR | QVVKTKVFQL | RQKGHDMKIE | VEAKIKEEGT | GIELTGIGSC | EIANALSKLK | FTKVNTNYRP |
| 370 | 380 | 390 | 400 | 410 | 420 | 430 | 440 | 450 | 460 | 470 | 480 |
| GLPFSGQVLL | VDEKGKPIPN | KNITSVVSPL | GYLSIFTTDE | HGLANISIDT | SNFTAPFLRV | VVTYKQNHVC | YDNWWLDEFH | TQADHSATLV | FSPSQSYIQL | ELVFGTLACG | QTQEIRIHYL |
| 490 | 500 | 510 | 520 | 530 | 540 | 550 | 560 | 570 | 580 | 590 | 600 |
| LNEDIMKNEK | DLTFYYLIKA | RGSIFNLGSH | VLSLEQGNMK | GVFSLPIQVE | PGMAPEAQLL | IYAILPNEEL | VADAQNFEIE | KCFANKVNLS | FPSAQSLPAS | DTHLKVKAAP | LSLCALTAVD |
| 610 | 620 | 630 | 640 | 650 | 660 | 670 | 680 | 690 | 700 | 710 | 720 |
| QSVLLLKPEA | KLSPQSIYNL | LPGKTVQGAF | FGVPVYKDHE | NCISGEDITH | NGIVYTPKHS | LGDNDAHSIF | QSVGINIFTN | SKIHKPRFCQ | EFQHYPAMGG | VAPQALAVAA | SGPGSSFRAM |
| 730 | 740 | 750 | 760 | 770 | 780 | 790 | 800 | 810 | 820 | 830 | 840 |
| GVPMMGLDYS | DEINQVVEVR | ETVRKYFPET | WIWDLVPLDV | SGDGELAVKV | PDTITEWKAS | AFCLSGTTGL | GLSSTISLQA | FQPFFLELTL | PYSVVRGEAF | TLKATVLNYM | SHCIQIRVDL |
| 850 | 860 | 870 | 880 | 890 | 900 | 910 | 920 | 930 | 940 | 950 | 960 |
| EISPDFLAVP | VGGHENSHCI | CGNERKTVSW | AVTPKSLGEV | NFTATAEALQ | SPELCGNKLT | EVPALVHKDT | VVKSVIVEPE | GIEKEQTYNT | LLCPQDTELQ | DNWSLELPPN | VVEGSARATH |
| 970 | 980 | 990 | 1000 | 1010 | 1020 | 1030 | 1040 | 1050 | 1060 | 1070 | 1080 |
| SVLGDILGSA | MQNLQNLLQM | PYGCGEQNMV | LFVPNIYVLN | YLNETQQLTE | AIKSKAINYL | ISGYQRQLNY | QHSDGSYSTF | GNHGGGNTPG | NTWLTAFVLK | AFAQAQSHIF | IEKTHITNAF |
| 1090 | 1100 | 1110 | 1120 | 1130 | 1140 | 1150 | 1160 | 1170 | 1180 | 1190 | 1200 |
| NWLSMKQKEN | GCFQQSGYLL | NNAMKGGVDD | EVTLSAYITI | ALLEMPLPVT | HSAVRNALFC | LETAWASISQ | SQESHVYTKA | LLAYAFALAG | NKAKRSELLE | SLNKDAVKEE | DSLHWQRPGD |
| 1210 | 1220 | 1230 | 1240 | 1250 | 1260 | 1270 | 1280 | 1290 | 1300 | 1310 | 1320 |
| VQKVKALSFY | QPRAPSAEVE | MTAYVLLAYL | TSESSRPTRD | LSSSDLSTAS | KIVKWISKQQ | NSHGGFSSTQ | DTVVALQALS | KYGAATFTRS | QKEVLVTIES | SGTFSKTFHV | NSGNRLLLQE |
| 1330 | 1340 | 1350 | 1360 | 1370 | 1380 | 1390 | 1400 | 1410 | 1420 | 1430 | 1440 |
| VRLPDLPGNY | VTKGSGSGCV | YLQTSLKYNI | LPVADGKAPF | ALQVNTLPLN | FDKAGDHRTF | QIRINVSYTG | ERPSSNMVIV | DVKMVSGFIP | MKPSVKKLQD | QPNIQRTEVN | TNHVLIYIEK |
| 1450 | 1460 | 1470 | 1480 | 1490 | 1500 | | | | | | |
| LTNQTLGFSF | AVEQDIPVKN | LKPAPIKVYD | YYETDEFTVE | EYSAPFSDGS | EQGNA | | | | | | |

| Cmpd. | m/z meas. | Δ m/z [ppm] | z | Rt [min] | Score | P | Range | Sequence | Modification |
|---|---|---|---|---|---|---|---|---|---|
| 1433 | 836.7841 | -29.03 | 3 | 62.28 | 59.99 | 0 | 588-611 | K.AAPLSLCALTAVDQSVLLLKPEAK.L | Carbamidomethyl: 7 |
| 603 | 435.7670 | -15.98 | 2 | 34.83 | 27.90 | 0 | 1316-1322 | R.LLLQEVR.L | |

| | | | |
|---|---|---|---|
| **Protein 4:** | 14-3-3 protein zeta/delta OS=Mus musculus GN=Ywhaz PE=1 SV=1 | | |
| **Accession:** | 1433Z_MOUSE | **Score:** | 79.00 |
| **Database:** | SwissProt | **MW [kDa]:** | 27.80 |
| **Seq. Coverage [%]:** | 12.20 % | **pI:** | 4.73 |
| | | **No. of Peptides:** | 3 |
| **Modification(s):** | Carbamidomethyl | | |

| 10 | 20 | 30 | 40 | 50 | 60 | 70 | 80 | 90 | 100 | 110 | 120 |
|---|---|---|---|---|---|---|---|---|---|---|---|
| MDKNELVQKA | KLAEQAERYD | DMAACMKSVT | EQGAELSNEE | RNLLSVAYKN | VVGARRSSWR | VVSSIEQKTE | GAEKKQQMAR | EYREKIETEL | RDICNDVLSL | LEKFLIPNAS | QPESKVFYLK |
| **130** | **140** | **150** | **160** | **170** | **180** | **190** | **200** | **210** | **220** | **230** | **240** |
| MKGDYYRYLA | EVAAGDDKKG | IVDQSQQAYQ | EAFEISKKEM | QPTHPIRLGL | ALNFSVFYYE | ILNSPEKACS | LAKTAFDEAI | AELDTLSEES | YKDSTLIMQL | LRDNLTLWTS | DTQGDEAEAG |
| **250** | | | | | | | | | | | |
| EGGEN | | | | | | | | | | | |

| Cmpd. | m/z meas. | Δ m/z [ppm] | z | Rt [min] | Score | P | Range | Sequence | Modification |
|---|---|---|---|---|---|---|---|---|---|
| 689 | 454.2583 | -17.02 | 2 | 37.56 | 14.06 | 0 | 42-49 | R.NLLSVAYK.N | |
| 1456 | 709.8440 | -27.32 | 2 | 63.13 | 37.34 | 0 | 92-103 | R.DICNDVLSLLEK.F | Carbamidomethyl: 3 |
| 1370 | 595.3181 | -26.81 | 2 | 60.03 | 27.60 | 0 | 213-222 | K.DSTLIMQLLR.D | |

| | | | |
|---|---|---|---|
| **Protein 5:** | Tubulin alpha-1B chain OS=Mus musculus GN=Tuba1b PE=1 SV=2 | | |
| **Accession:** | TBA1B_MOUSE | **Score:** | 68.06 |
| **Database:** | SwissProt | **MW [kDa]:** | 50.10 |
| **Seq. Coverage [%]:** | 3.10 % | **pI:** | 4.94 |
| | | **No. of Peptides:** | 1 |

| | | | | | | | | | | | |
|---|---|---|---|---|---|---|---|---|---|---|---|
| 10 | 20 | 30 | 40 | 50 | 60 | 70 | 80 | 90 | 100 | 110 | 120 |
| MRECISIHVG | QAGVQIGNAC | WELYCLEHGI | QPDGQMPSDK | TIGGGDDSFN | TFFSETGAGK | HVPRAVFVDL | EPTVIDEVRT | GTYRQLFHPE | QLITGKEDAA | NNYARGHYTI | GKEIIDLVLD |
| 130 | 140 | 150 | 160 | 170 | 180 | 190 | 200 | 210 | 220 | 230 | 240 |
| RIRKLADQCT | GLQGFLVFHS | FGGGTGSGFT | SLLMERLSVD | YGKKSKLEFS | IYPAPQVSTA | VVEPYNSILT | THTTLEHSDC | AFMVDNEAIY | DICRRNLDIE | RPTYTNLNRL | ISQIVSSITA |
| 250 | 260 | 270 | 280 | 290 | 300 | 310 | 320 | 330 | 340 | 350 | 360 |
| SLRFDGALNV | DLTEFQTNLV | PYPRIHFPLA | TYAPVISAEK | AYHEQLSVAE | ITNACFEPAN | QMVKCDPRHG | KYMACCLLYR | GDVVPKDVNA | AIATIKTKRS | IQFVDWCPTG | FKVGINYQPP |
| 370 | 380 | 390 | 400 | 410 | 420 | 430 | 440 | 450 | 460 | | |
| TVVPGGDLAK | VQRAVCMLSN | TTAIAEAWAR | LDHKFDLMYA | KRAFVHWYVG | EGMEEGEFSE | AREDMAALEK | DYEEVGVDSV | EGEGEEEGEE | Y | | |

| Cmpd. | m/z meas. | Δ m/z [ppm] | z | Rt [min] | Score | P | Range | Sequence | Modification |
|---|---|---|---|---|---|---|---|---|---|
| 1454 | 744.4232 | -26.95 | 2 | 63.01 | 68.06 | 0 | 230-243 | R.LISQIVSSITASLR.F | |

**Protein 6:** Ig kappa chain C region OS=Mus musculus PE=1 SV=1

**Accession:** IGKC_MOUSE

**Database:** SwissProt

**Seq. Coverage [%]:** 13.20 %

**Score:** 56.93

**MW [kDa]:** 11.80

**pI:** 5.23

**No. of Peptides:** 1

| | | | | | | | | | | |
|---|---|---|---|---|---|---|---|---|---|---|
| 10 | 20 | 30 | 40 | 50 | 60 | 70 | 80 | 90 | 100 | 110 |
| ADAAPTVSIF | PPSSEQLTSG | GASVVCFLNN | FYPKDINVKW | KIDGSERQNG | VLNSWTDQDS | KDSTYSMSST | LTLTKDEYER | HNSYTCEATH | KTSTSPIVKS | FNRNEC |

| Cmpd. | m/z meas. | Δ m/z [ppm] | z | Rt [min] | Score | P | Range | Sequence | Modification |
|---|---|---|---|---|---|---|---|---|---|
| 779 | 767.8552 | -17.81 | 2 | 40.37 | 56.93 | 0 | 62-75 | K.DSTYSMSSTLTLTK.D | |

| | | | |
|---|---|---|---|
| **Protein 7:** | Complement C4-B OS=Mus musculus GN=C4b PE=1 SV=3 | | |
| **Accession:** | CO4B_MOUSE | **Score:** | 48.65 |
| **Database:** | SwissProt | **MW [kDa]:** | 192.80 |
| **Seq. Coverage [%]:** | 1.40 % | **pI:** | 7.38 |
| | | **No. of Peptides:** | 1 |

| 10 | 20 | 30 | 40 | 50 | 60 | 70 | 80 | 90 | 100 | 110 | 120 |
|---|---|---|---|---|---|---|---|---|---|---|---|
| MRLLWGLAWV | FSFCASSLQK | PRLLLFSPSV | VNLGTPLSVG | VQLLDAPPGQ | EVKGSVFLRN | PKGGSCSPKK | DFKLSSGDDF | VLLSLEVPLE | DVRSCGLFDL | RRAPHIQLVA | QSPWLRNTAF |
| 130 | 140 | 150 | 160 | 170 | 180 | 190 | 200 | 210 | 220 | 230 | 240 |
| KATETQGVNL | LFSSRRGHIF | VQTDQPIYNP | GQRVRYRVFA | LDQKMRPSTD | FLTITVENSH | GLRVLKKEIF | TSTSIFQDAF | TIPDISEPGT | WKISARFSDG | LESNRSTHFE | VKKYVLPNFE |
| 250 | 260 | 270 | 280 | 290 | 300 | 310 | 320 | 330 | 340 | 350 | 360 |
| VKITPWKPYI | LMVPSNSDEI | QLDIQARYIY | GKPVQGVAYT | RFALMDEQGK | RTFLRGLETQ | AKLVEGRTHI | SISKDQFQAA | LDKINIGVRD | LEGLRLYAAT | AVIESPGGEM | EEAELTSWRF |
| 370 | 380 | 390 | 400 | 410 | 420 | 430 | 440 | 450 | 460 | 470 | 480 |
| VSSAFSLDLS | RTKRHLVPGA | HFLLQALVQE | MSGSEASNVP | VKVSATLVSG | SDSQVLDIQQ | STNGIGQVSI | SFPIPPTVTE | LRLLVSAGSL | YPAIARLTVQ | APPSRGTGFL | SIEPLDPRSP |
| 490 | 500 | 510 | 520 | 530 | 540 | 550 | 560 | 570 | 580 | 590 | 600 |
| SVGDTFILNL | QPVGIPAPTF | SHYYYMIISR | GQIMAMGREP | RKTVTSVSVL | VDHQLAPSFY | FVAYFYHQGH | PVANSLLINI | QSRDCEGKLQ | LKVDGAKEYR | NADMMKLRIQ | TDSKALVALG |
| 610 | 620 | 630 | 640 | 650 | 660 | 670 | 680 | 690 | 700 | 710 | 720 |
| AVDMALYAVG | GRSHKPLDMS | KVFEVINSYN | VGCGPGGGDD | ALQVFQDAGL | AFSDGDRLTQ | TREDLSCPKE | KKSRQKRNVN | FQKAVSEKLG | QYSSPDAKRC | CQDGMTKLPM | KRTCEQRAAR |
| 730 | 740 | 750 | 760 | 770 | 780 | 790 | 800 | 810 | 820 | 830 | 840 |
| VPQQACREPF | LSCCKFAEDL | RRNQTRSQAH | LARNNHNMLQ | EEDLIDEDDI | LVRTSFPENW | LWRVEPVDSS | KLLTVWLPDS | MTTWEIHGVS | LSKSKGLCVA | KPTRVRVFRK | FHLHLRLPIS |
| 850 | 860 | 870 | 880 | 890 | 900 | 910 | 920 | 930 | 940 | 950 | 960 |
| IRRFEQFELR | PVLYNYLNDD | VAVSVHVTPV | EGLCLAGGGM | MAQQVTVPAG | SARPVAFSVV | PTAAANVPLK | VVARGVFDLG | DAVSKILQIE | KEGAIHREEL | VYNLDPLNNL | GRTLEIPGSS |
| 970 | 980 | 990 | 1000 | 1010 | 1020 | 1030 | 1040 | 1050 | 1060 | 1070 | 1080 |
| DPNIVPDGDF | SSLVRVTASE | PLETMGSEGA | LSPGGVASLL | RLPQGCAEQT | MIYLAPTLTA | SNYLDRTEQW | SKLSPETKDH | AVDLIQKGYM | RIQQFRKNDG | SFGAWLHRDS | STWLTAFVLK |
| 1090 | 1100 | 1110 | 1120 | 1130 | 1140 | 1150 | 1160 | 1170 | 1180 | 1190 | 1200 |
| ILSLAQEQVG | NSPEKLQETA | SWLLAQQLGD | GSFHDPCPVI | HRAMQGGLVG | SDETVALTAF | VVIALHHGLD | VFQDDDAKQL | KNRVEASITK | ANSFLGQKAS | AGLLGAHAAA | ITAYALTLTK |
| 1210 | 1220 | 1230 | 1240 | 1250 | 1260 | 1270 | 1280 | 1290 | 1300 | 1310 | 1320 |
| ASEDLRNVAH | NSLMAMAEET | GEHLYWGLVL | GSQDKVVLRP | TAPRSPTEPV | PQAPALWIET | TAYALLHLLL | REGKGKMADK | AASWLTHQGS | FHGAFRSTQD | TVVTLDALSA | YWIASHTTEE |
| 1330 | 1340 | 1350 | 1360 | 1370 | 1380 | 1390 | 1400 | 1410 | 1420 | 1430 | 1440 |
| KALNVTLSSM | GRNGLKTHGL | HLNNHQVKGL | EEELKFSLGS | TISVKVEGNS | KGTLKILRTY | NVLDMKNTTC | QDLQIEVKVT | GAVEYAWDAN | EDYEDYYDMP | AADDPSVPLQ | PVTPLQLFEG |
| 1450 | 1460 | 1470 | 1480 | 1490 | 1500 | 1510 | 1520 | 1530 | 1540 | 1550 | 1560 |
| RRSRRRREAP | KVVEEQESRV | QYTVCIWRNG | KLGLSGMAIA | DITLLSGFHA | LRADLEKLTS | LSDRYVSHFE | TDGPHVLLYF | DSVPTTRECV | GFGASQEVVV | GLVQPSSAVL | YDYYSPDHKC |
| 1570 | 1580 | 1590 | 1600 | 1610 | 1620 | 1630 | 1640 | 1650 | 1660 | 1670 | 1680 |
| SVFYAAPTKS | QLLATLCSGD | VCQCAEGKCP | RLLRSLERRV | EDKDGYRMRF | ACYYPRVEYG | FTVKVLREDG | RAAFRLFESK | ITQVLHFRKD | TMASIGQTRN | FLSRASCRLR | LEPNKEYLIM |
| 1690 | 1700 | 1710 | 1720 | 1730 | 1740 | | | | | | |
| GMDGETSDNK | GDPQYLLDSN | TWIEEMPSEQ | MCKSTRHRAA | CFQLKDFLME | FSSRGCQV | | | | | | |

| Cmpd. | m/z meas. | Δ m/z [ppm] | z | Rt [min] | Score | P | Range | Sequence | Modification |
|---|---|---|---|---|---|---|---|---|---|
| 1410 | 922.7611 | -28.21 | 3 | 61.44 | 48.65 | 0 | 1297-1321 | R.STQDTVVTLDALSAYWIASHTTEEK.A | |

**Protein 8:** Fatty acid-binding protein, epidermal OS=Mus musculus GN=Fabp5 PE=1 SV=3

| | | | |
|---|---|---|---|
| **Accession:** | FABP5_MOUSE | **Score:** | 40.94 |
| **Database:** | SwissProt | **MW [kDa]:** | 15.10 |
| **Seq. Coverage [%]:** | 6.70 % | **pI:** | 6.14 |
| | | **No. of Peptides:** | 1 |

```
        10         20         30         40         50         60         70         80         90        100        110        120
MASLKDLEGK WRLMESHGFE EYMKELGVGL ALRKMAAMAK PDCIITCDGN NITVKTESTV KTTVFSCNLG EKFDETTADG RKTETVCTFQ DGALVQHQQW DGKESTITRK LKDGKMIVEC
       130        140
VMNNATCTRV YEKVQ
```

| Cmpd. | m/z meas. | Δ m/z [ppm] | z | Rt [min] | Score | P | Range | Sequence | Modification |
|---|---|---|---|---|---|---|---|---|---|
| 770 | 464.2773 | -16.08 | 2 | 40.09 | 40.94 | 0 | 25-33 | K.ELGVGLALR.K | |

**Protein 9:** Thrombospondin-1 OS=Mus musculus GN=Thbs1 PE=1 SV=1

| | | | |
|---|---|---|---|
| **Accession:** | TSP1_MOUSE | **Score:** | 38.88 |
| **Database:** | SwissProt | **MW [kDa]:** | 129.60 |
| **Seq. Coverage [%]:** | 0.90 % | **pI:** | 4.72 |
| | | **No. of Peptides:** | 1 |

| 10 | 20 | 30 | 40 | 50 | 60 | 70 | 80 | 90 | 100 | 110 | 120 |
|---|---|---|---|---|---|---|---|---|---|---|---|
| MELLRGLGVL | FLLHMCGSNR | IPESGGDNGV | FDIFELIGGA | RRGPGRRLVK | GQDLSSPAFR | IENANLIPAV | PDDKFQDLLD | AVWADKGFIF | LASLRQMKKT | RGTLLAVERK | DNTGQIFSVV |
| 130 | 140 | 150 | 160 | 170 | 180 | 190 | 200 | 210 | 220 | 230 | 240 |
| SNGKAGTLDL | SLSLPGKQQV | VSVEEALLAT | GQWKSITLFV | QEDRAQLYID | CDKMESAELD | VPIQSIFTRD | LASVARLRVA | KGDVNDNFQG | VLQNVRFVFG | TTPEDILRNK | GCSSSTNVLL |
| 250 | 260 | 270 | 280 | 290 | 300 | 310 | 320 | 330 | 340 | 350 | 360 |
| TLDNNVVNGS | SPAIRTNYIG | HKTKDLQAIC | GLSCDELSSM | VLELKGLRTI | VTTLQDSIRK | VTEENRELVS | ELKRPPLCFH | NGVQYKNNEE | WTVDSCTECH | CQNSVTICKK | VSCPIMPCSN |
| 370 | 380 | 390 | 400 | 410 | 420 | 430 | 440 | 450 | 460 | 470 | 480 |
| ATVPDGECCP | RCWPSDSADD | GWSPWSEWTS | CSATCGNGIQ | QRGRSCDSLN | NRCEGSSVQT | RTCHIQECDK | RFKQDGGWSH | WSPWSSCSVT | CGDGVITRIR | LCNSPSPQMN | GKPCEGEARE |
| 490 | 500 | 510 | 520 | 530 | 540 | 550 | 560 | 570 | 580 | 590 | 600 |
| TKACKKDACP | INGGWGPWSP | WDICSVTCGG | GVQRRSRLCN | NPTPQFGGKD | CVGDVTENQV | CNKQDCPIDG | CLSNPCFAGA | KCTSYPDGSW | KCGACPPGYS | GNGIQCKDVD | ECKEVPDACF |
| 610 | 620 | 630 | 640 | 650 | 660 | 670 | 680 | 690 | 700 | 710 | 720 |
| NHNGEHRCKN | TDPGYNCLPC | PPRFTGSQPF | GRGVEHAMAN | KQVCKPRNPC | TDGTHDCNKN | AKCNYLGHYS | DPMYRCECKP | GYAGNGIICG | EDTDLDGWPN | ENLVCVANAT | YHCKKDNCPN |
| 730 | 740 | 750 | 760 | 770 | 780 | 790 | 800 | 810 | 820 | 830 | 840 |
| LPNSGQEDYD | KDGIGDACDD | DDDNDKIPDD | RDNCPFHYNP | AQYDYDRDDV | GDRCDNCPYN | HNPDQADTDK | NGEGDACAVD | IDGDGILNER | DNCQYVYNVD | QRDTDMDGVG | DQCDNCPLEH |
| 850 | 860 | 870 | 880 | 890 | 900 | 910 | 920 | 930 | 940 | 950 | 960 |
| NPDQLDSDSD | LIGDTCDNNQ | DIDEDGHQNN | LDNCPYVPNA | NQADHDKDGK | GDACDHDDDN | DGIPDDRDNC | RLVPNPDQKD | SDGDGRGDAC | KDDFDHDNVP | DIDDICPENF | DISETDFRRF |
| 970 | 980 | 990 | 1000 | 1010 | 1020 | 1030 | 1040 | 1050 | 1060 | 1070 | 1080 |
| QMIPLDPKGT | SQNDPNWVVR | HQGKELVQTV | NCDPGLAVGY | DEFNAVDFSG | TFFINTERDD | DYAGFVFGYQ | SSSRFYVVMW | KQVTQSYWDT | NPTRAQGYSG | LSVKVVNSTT | GPGEHLRNAL |
| 1090 | 1100 | 1110 | 1120 | 1130 | 1140 | 1150 | 1160 | 1170 | 1180 | | |
| WHTGNTPGQV | RTLWHDPRHI | GWKDFTAYRW | RLSHRPKTGY | IRVVMYEGKK | IMADSGPIYD | KTYAGGRLGL | FVFSQEMVFF | SDMKYECRDS | | | |

| Cmpd. | m/z meas. | Δ m/z [ppm] | z | Rt [min] | Score | P | Range | Sequence | Modification |
|---|---|---|---|---|---|---|---|---|---|
| 918 | 623.8416 | -19.43 | 2 | 44.90 | 38.88 | 0 | 289-299 | R.TIVTTLQDSIR.K | |

| | | | |
|---|---|---|---|
| **Protein 10:** | Arf-GAP with SH3 domain, ANK repeat and PH domain-containing protein 2 OS=Mus musculus GN=Asap2 PE=1 SV=3 | | |
| **Accession:** | ASAP2_MOUSE | **Score:** | 27.60 |
| **Database:** | SwissProt | **MW [kDa]:** | 106.70 |
| **Seq. Coverage [%]:** | 0.80 % | **pI:** | 6.22 |
| | | **No. of Peptides:** | 1 |

| 10 | 20 | 30 | 40 | 50 | 60 | 70 | 80 | 90 | 100 | 110 | 120 |
|---|---|---|---|---|---|---|---|---|---|---|---|
| MPDQISVSEF | VAETHEDYKA | PTASSFTTRT | AQCRNTVAAI | EEALDVDRMV | LYKMKKSVKA | INISGLAHVE | NEEQYTQALE | KFGGNCVCRD | DPDLGSAFLK | FSVFTKELTA | LFKNLIQNMN |
| 130 | 140 | 150 | 160 | 170 | 180 | 190 | 200 | 210 | 220 | 230 | 240 |
| NIISFPLDSL | LKGDLKGVKG | DLKKPFDKAW | KDYETKITKI | EKEKKEHAKL | HGMIRTEISG | AEIAEEMEKE | RRFFQLQMCE | YLLKVNEIKV | KKGVDLLQNL | IKYFHAQCNF | FQDGLKAVES |
| 250 | 260 | 270 | 280 | 290 | 300 | 310 | 320 | 330 | 340 | 350 | 360 |
| LKPSIETLST | DLHTIKQAQD | EERRQLIQLR | DILKSALQVE | QKESRRDSQL | RQSTAYSLHQ | PQGNKEHGTE | RNGNLYKKSD | GIRKVWQKRK | CSVKNGFLTI | SHGTANRPPA | KLNLLTCQVK |
| 370 | 380 | 390 | 400 | 410 | 420 | 430 | 440 | 450 | 460 | 470 | 480 |
| TNPEEKKCFD | LISHDRTYHF | QAEDEQECQI | WMSVLQNSKE | EALNNAFKGD | DNTGENNIVQ | ELTKEIISEV | QRMTGNDVCC | DCGAPDPTWL | STNLGILTCI | ECSGIHRELG | VHYSRMQSLT |
| 490 | 500 | 510 | 520 | 530 | 540 | 550 | 560 | 570 | 580 | 590 | 600 |
| LDVLGTSELL | LAKNIGNAGF | NEIMECCLPS | EDPVKPNPGS | DMIARKDYIT | AKYMERRYAR | KKHADTAAKL | HSLCEAVKTR | DIFGLLQAYA | DGVDLTEKIP | LANGHEPDET | ALHLAVRSVD |
| 610 | 620 | 630 | 640 | 650 | 660 | 670 | 680 | 690 | 700 | 710 | 720 |
| RTSLHIVDFL | VQNSGNLDKQ | TGKGSTALHY | CCLTDNAECL | KLLLRGKASI | EIANESGETP | LDIAKRLKHE | HCEELLTQAL | SGRFNSHVHV | EYEWRLLHED | LDESDDDVDE | KLQPSPNRRE |
| 730 | 740 | 750 | 760 | 770 | 780 | 790 | 800 | 810 | 820 | 830 | 840 |
| DRPVSFYQLG | SSQFQSNAVS | LARDTANLTK | DKQRGFGPSI | LQNETYGAIL | SGSPPSSQSI | PPSTTSAPPL | PPRNVGKDPL | TTTPPPPVAK | TSGTLEAMNQ | PSKSSQPGTS | QSKPPPLPPQ |
| 850 | 860 | 870 | 880 | 890 | 900 | 910 | 920 | 930 | 940 | 950 | 960 |
| PPSRLPQKKP | ASGTDKPTPL | TNKGQPRGPE | ASGPLSNAMA | LQPPAPMPRK | SQATKSKPKR | VKALYNCVAD | NPDELTFSEG | DVIIVDGEED | QEWWIGHIDG | EPSRKGAFPV | SFVHFIAD |

| Cmpd. | m/z meas. | Δ m/z [ppm] | z | Rt [min] | Score | P | Range | Sequence | Modification |
|---|---|---|---|---|---|---|---|---|---|
| 493 | 487.2636 | -11.70 | 2 | 31.62 | 27.60 | 0 | 425-432 | K.EIISEVQR.M | |

**Protein 11:** Potassium voltage-gated channel subfamily S member 1 OS=Mus musculus GN=Kcns1 PE=1 SV=2

**Accession:** KCNS1_MOUSE

**Database:** SwissProt

**Seq. Coverage [%]:** 3.00 %

**Score:** 26.19

**MW [kDa]:** 54.90

**pI:** 6.65

**No. of Peptides:** 1

**Modification(s):** Oxidation

BRUKER

| 10 | 20 | 30 | 40 | 50 | 60 | 70 | 80 | 90 | 100 | 110 | 120 |
|---|---|---|---|---|---|---|---|---|---|---|---|
| MVSEFPGPGS | RVPWRPRDEA | LRVNVGGVRR | LLSARALARF | PGTRLGRLQA | AASEEQARRL | CDDYDAAAHE | FYFDRHPGFF | LGLLHFYRTG | HLHVLDELCV | FAFGQEADYW | GLGENALATC |
| **130** | **140** | **150** | **160** | **170** | **180** | **190** | **200** | **210** | **220** | **230** | **240** |
| CRARYLERRV | ARPRAWDEDS | DAPSSVDPCP | DEISDVQREL | ARYGAARCGR | LRRRLWLTME | NPGYSLPSKL | FSCVSIGVVL | ASIAAMCIHS | LPEYQAREAA | AAVAAVAAGR | SAEEVRDDPV |
| **250** | **260** | **270** | **280** | **290** | **300** | **310** | **320** | **330** | **340** | **350** | **360** |
| LRRLEYFCIA | WFSFEVSSRL | LLAPSTRNFF | CHPLNLIDIV | SVLPFYLTLL | AGAALGDQRG | ASGEELGDLG | KVVQVFRLMR | IFRVLKLARH | STGLRSLGAT | LKHSYREVGI | LLLYLAVGVS |
| **370** | **380** | **390** | **400** | **410** | **420** | **430** | **440** | **450** | **460** | **470** | **480** |
| VFSGVAYTAE | EENEGFHTIP | ACWWWGTVSM | TTVGYGDVVP | ETVGGKLAAS | GCILGGILVV | ALPITIIFNK | FSHFYRRQKA | LEAAVRSSGQ | REFEDLLSSV | DGVSDVSLET | SRDTSQEGRS |
| **490** | **500** | | | | | | | | | | |
| TDLETQAPRE | PAKSHSY | | | | | | | | | | |

| Cmpd. | m/z meas. | Δ m/z [ppm] | z | Rt [min] | Score | P | Range | Sequence | Modification |
|---|---|---|---|---|---|---|---|---|---|
| 1209 | 584.6104 | -28.94 | 3 | 54.54 | 11.00 | 0 | 175-189 | R.LWLTMENPGYSLPSK.L | Oxidation: 5 |

**Protein 12:** ATPase family AAA domain-containing protein 3 OS=Mus musculus GN=Atad3 PE=1 SV=1

**Accession:** ATAD3_MOUSE

**Database:** SwissProt

**Seq. Coverage [%]:** 1.70 %

**Score:** 23.16

**MW [kDa]:** 66.70

**pI:** 9.32

**No. of Peptides:** 1

| 10 | 20 | 30 | 40 | 50 | 60 | 70 | 80 | 90 | 100 | 110 | 120 |
|---|---|---|---|---|---|---|---|---|---|---|---|
| MSWLFGIKGP | KGEGTGPPLP | LPPAQPGAEG | GGDRGAGDRP | SPKDKWSNFD | PTGLERAAKA | ARELEHSRHA | KEALSLAQMQ | EQTLQLEQQS | KLKEYEAAVE | QLKSEQIRVQ | AEERRKTLTE |
| **130** | **140** | **150** | **160** | **170** | **180** | **190** | **200** | **210** | **220** | **230** | **240** |
| ETRQHQARAQ | YQDKLARQRY | EDQLKQQQLL | NEENLRKQEE | SVQKQEAIRR | ATVEREMELR | HKNEMLRVEA | EARARAKADR | ENADIIREQI | RLKAAEHRQT | ILESIRTAGT | LLGEGFRAFV |
| **250** | **260** | **270** | **280** | **290** | **300** | **310** | **320** | **330** | **340** | **350** | **360** |
| TDWDKVTATV | AGLTLLAVGV | YSAKNATSVA | GRYIEARLGK | PSLVRETSRI | SVLEALRHPI | QVSRRLVSRP | QDALEGVILS | PSLEARVRDI | AIATRNTKKN | KSLYRNVLMY | GPPGTGKTLF |
| **370** | **380** | **390** | **400** | **410** | **420** | **430** | **440** | **450** | **460** | **470** | **480** |
| AKKLALHSGM | DYAIMTGGDV | APMGREGVTA | MHKVFDWAST | SRRGLLLFVD | EADAFLRKRA | TEKISEDLRA | TLNAFLHRTG | QHSSKFMLVL | ASNQPEQFDW | AINDRIDEMV | CFALPQREER |
| **490** | **500** | **510** | **520** | **530** | **540** | **550** | **560** | **570** | **580** | **590** | **600** |
| ERLVRMYFDK | YVLKPATEGK | QRLKVAQFDY | GKKCSEVAQL | TEGMSGREIA | QLAVAWQAMA | YSSEDGVLTE | AMMDARVQDA | VQQHQQKMQW | LKVERPDSQT | NKPPHPSLLS | C |

| Cmpd. | m/z meas. | Δ m/z [ppm] | z | Rt [min] | Score | P | Range | Sequence | Modification |
|---|---|---|---|---|---|---|---|---|---|
| 530 | 590.2965 | -2.91 | 2 | 32.65 | 23.16 | 0 | 94-103 | K.EYEAAVEQLK.S | |
